# RAPTOR: A Five-Safes approach to a secure, cloud native and serverless genomics data repository

**DOI:** 10.1101/2022.10.27.514127

**Authors:** Chih Chuan Shih, Jieqi Chen, Ai Shan Lee, Nicolas Bertin, Maxime Hebrard, Chiea Chuen Khor, Zheng Li, Joanna Hui Juan Tan, Wee Yang Meah, Su Qin Peh, Shi Qi Mok, Kar Seng Sim, Jianjun Liu, Ling Wang, Eleanor Wong, Jingmei Li, Aung Tin, Ching-Yu Cheng, Chew-Kiat Heng, Jian-Min Yuan, Woon-Puay Koh, Seang Mei Saw, Yechiel Friedlander, Xueling Sim, Jin Fang Chai, Yap Seng Chong, Sonia Davila, Liuh Ling Goh, Eng Sing Lee, Tien Yin Wong, Neerja Karnani, Khai Pang Leong, Khung Keong Yeo, John C Chambers, Su Chi Lim, Rick Siow Mong Goh, Patrick Tan, Rajkumar Dorajoo

**Affiliations:** Genome Institute of Singapore, Agency for Science, Technology, and Research, Singapore; Singapore Eye Research institute, Singapore National Eye Centre, Singapore; Yong Loo Lin School of Medicine, National University of Singapore; Duke-NUS Medical School, Singapore; Singapore National Eye Centre, Singapore; Department of Paediatrics, Yong Loo Lin School of Medicine, National University of Singapore; Khoo Teck Puat - National University Children’s Medical Institute, National University Health System, Singapore; Department of Epidemiology, Graduate School of Public Health, University of Pittsburgh, Pittsburgh, Pennsylvania, USA; UPMC Hillman Cancer Center, University of Pittsburgh, Pittsburgh, Pennsylvania, USA; Healthy Longevity Translational Research Programme, Yong Loo Lin School of Medicine, National University of Singapore; Singapore Institute for Clinical Sciences, Agency for Science Technology and Research, Singapore; Saw Swee Hock School of Public Health, National University of Singapore; Braun School of Public Health, Hebrew University of Jerusalem, Jerusalem, Israel; Precision Health Research, Singapore; SingHealth Duke-NUS Institute of Precision Medicine, Singapore; Molecular Diagnostic Laboratory, Tan Tock Seng Hospital, Singapore; Clinical Research Unit, National Healthcare Group Polyclinics, Singapore; Lee Kong Chian School of Medicine, Nanyang Technological University, Singapore; Bioinformatics Institute, Agency for Science, Technology, and Research, Singapore; Personalised Medicine Service, Tan Tock Seng Hospital, Singapore; Department of Cardiology, National Heart Centre Singapore, Singapore; School of Public Health, Imperial College London, United Kingdom; Clinical Research Unit, Khoo Teck Puat Hospital, Singapore; Diabetes Centre, Admiralty Medical Centre, Singapore; Institute of High Performance Computing, Agency for Science, Technology, and Research, Singapore; Cancer Science Institute of Singapore, National University of Singapore, Singapore

**Author notes:** Senior Authors (Patrick Tan, Rajkumar Dorajoo). Corresponding Authors: Rajkumar Dorajoo, Chih Chuan Shih : Genome Institute of Singapore, 60 Biopolis Street, Genome, #02-01, Singapore 138672.

## Abstract

Genomic researchers are increasingly utilizing commercial cloud platforms (CCPs) to manage their data and analytics needs. Commercial clouds allow researchers to grow their storage and analytics capacity on demand, keeping pace with expanding project data footprints and enabling researchers to avoid large capital expenditures while paying only for IT capacity consumed by their project. Cloud computing also allows researchers to overcome common network and storage bottlenecks encountered when combining or re-analysing large datasets. However, cloud computing presents a new set of challenges. Without adequate security controls, the risk of unauthorised access may be higher for data stored on the cloud. In addition, regulators are increasingly mandating data access patterns and specific security protocols on the storage and use of genomic data to safeguard rights of the study participants. While CCPs provide tools for security and regulatory compliance, utilising these tools to build the necessary controls required for cloud solutions is not trivial as such skill sets are not commonly found in a genomics lab. The Research Assets Provisioning and Tracking Online Repository (RAPTOR) by the Genome Institute of Singapore is a cloud native genomics data repository and analytics platform focusing on security and regulatory compliance. Using a “five-safes” framework (Safe Purpose, Safe People, Safe Settings, Safe Data and Safe Output), RAPTOR provides security and governance controls to data contributors and users leveraging cloud computing for sharing and analysis of large genomic datasets without the risk of security breaches or running afoul of regulations. RAPTOR can also enable data federation with other genomic data repositories using GA4GH community-defined standards, allowing researchers to boost the statistical power of their work and overcome geographic and ancestry limitations of data sets

## Introduction

Data footprint for genomics projects has increased rapidly. The UK Biobank for example, holds about 11 petabytes of genomic data, which is projected to grow above 40 petabytes by 2025^1^. To effectively store and process data at petabyte-scale requires significant IT capacity, operated by experienced specialists. These capacities are often out-of-reach for small to medium size companies and academic labs. There is also the added issue of effectively sharing large-scale data with collaborators. To transfer 1 petabyte of data across a network which allows actual sustained throughput (not line speed) of 1024Mbps will take more than 90 days. If data is moved through the public internet, the transfer time is estimated to be at least 5 times longer^2^.

Public cloud computing platforms, such as Amazon Web Services (AWS), Google Cloud Platform (GCP) and Microsoft Azure, provide feasible ways around these constraints. Commercial clouds provide elastic and scalable IT resources (i.e servers and storage) allowing users to grow their IT infrastructure in tandem with data generation without loss in reliability, availability or performance^3–5^. Operators can also adjust the scale and subsequent costs of cloud-based IT operations on demand, to match different project phases. In contrast, operators building on-premise data centres must build-in sufficient capacity to take on the peak load of the project and not the most common load level. Therefore, an on-premise system operator will have to bear the cost of maintaining the entire system designed for peak usage, even during lull periods when most of the servers are idle. In addition, once data is available on the cloud, collaboration and sharing can be achieved by having users run analytics within the same “cloud region” where the data resides, effectively side-stepping the challenge of moving large data across networks. The AnVIL project, for example, leverages GCP to host and share more than 3 petabytes of genomic data^6^. For analysis, AnVIL provides its users with the ability to work directly on cloud by integrating with various analytics platforms, including Galaxy, Juypter and Dockstore. These platforms enable users to bring computation to the data repository, effectively “inverting the model of data sharing”^6^. However, cloud computing can introduce significant security risks. Without well designed access controls, data placed on cloud can potentially be accessed from anywhere and by anyone with internet access. Active data on cloud may spend considerable time moving through storage and servers shared with many other users, allowing data to be silently replicated many times over while in flight. Furthermore, cloud data sharing requires setting up an endpoint which is accessible from the internet. As evidenced by remote attacks through OpenSSH with GNU Bash, this may create opportunities for malicious actors who can either break in using forged credentials or vulnerabilities in the software used to host the data^7^.

There have been several initiatives to address these security concerns of genomic data. The Global Alliance for Genomics and Health (GA4GH), has developed protocols, tools, and policies for responsible sharing of genomics data^8^. In particular, the GA4GH Data Security workstream has delivered a data security infrastructure policy which outlines recommendations for securing IT infrastructures used for genomic and clinical data^9^. The Authentication and Authorisation Infrastructure guide additionally provides a comprehensive framework for cloud users to safely authenticate users and assign authorisations for use of data^10^. The Ministry of Health of Singapore has also issued a HealthTech Instruction Manual providing instructions for IT and data governance^11^. These include standard procedures and algorithms for data encryption, configuration of network partition and data access points, and management and securing user accounts^12^.

Major Cloud Service Providers (CSP) already supply many tools needed to build a secure platform following best practices and compliance to regulations^13–15^. However, this remains a complex task requiring deep information technology expertise and knowledge of relevant regulations that may not be available in a genomics lab^16^. The Research Assets Provisioning and Tracking Online Repository (RAPTOR) was developed to fill this need. RAPTOR is a serverless, cloud native, genomics data repository and analytics platform that focuses on data security and regulatory compliance (Figure 1). AWS is the current CSP for RAPTOR. The goal of RAPTOR is to provide a secure, regulatory compliant platform for researchers to leverage elasticity and scalability of cloud computing for large-scale data analysis. With data-in-place analytics, RAPTOR also circumvents the challenge of transferring large data sets across public networks while providing accountability of data usage.

**Figure 1:**
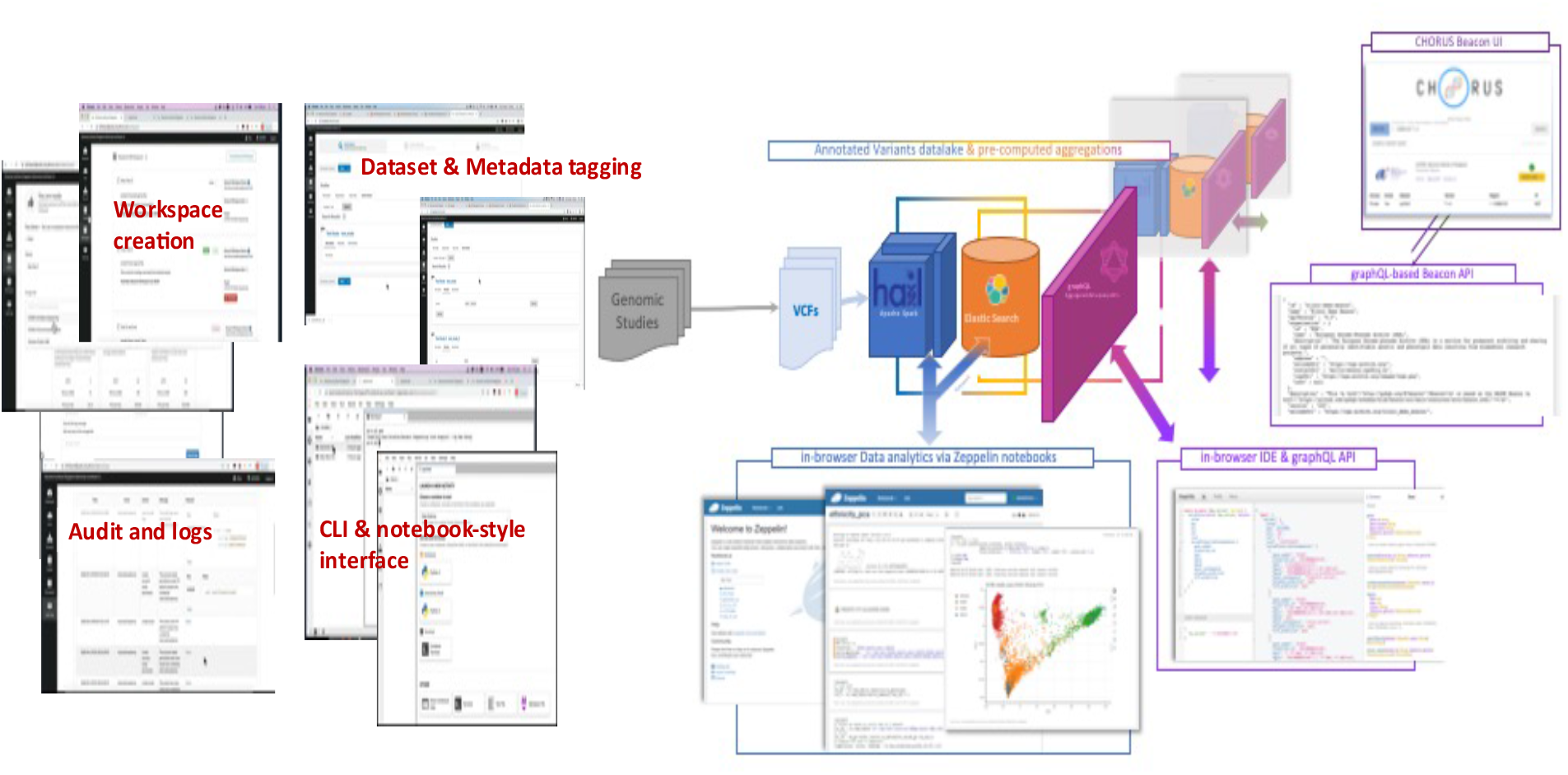
RAPTOR Overview and the modes of data analysis supported. RAPTOR provides access via Standalone Linux machines with sudo access, Juypter Notebooks, and EMR Cluster with preconfigured Hail tools.

## Results

RAPTOR is designed to be foundationally secure and embedded with all elements essential for data governance. Security and data governance strategies and procedures are considered and designed into the platform using a “5 Safes” framework–Safe Purpose, Safe People, Safe Settings, Safe Data and Safe Output.

### Safe Purpose

Safe purpose refers to measures adopted to ensure data contributors’ control of data deposited with RAPTOR. Users who wish to access a dataset must submit an access request on the platform. Mandatory fields of this request include their proposal and the duration of their access. Users who wish to access datasets beyond the original deadline must submit an extension request through the platform.

It is mandatory to provide at least one data access committee (DAC) contact when depositing data onto RAPTOR and RAPTOR will forward any access request to the relevant DAC. After evaluation, the DAC has the option to approve or deny the access request using RAPTOR’s data management console. The DAC may also choose to grant access to specific subsets of files or allow access with modified parameters. For instance, a DAC may choose to modify certain data access expiry dates for specific sub-datasets on the same console.

RAPTOR hosts data on AWS S3. The hosting buckets are configured to block all access except those coming through a specific S3 Endpoint. An S3 Endpoint is analogous to a proxy for a webserver, providing an interface for the bucket to interact with the outside world without any direct connection. Access policies can be applied to the endpoint to govern traffic to the bucket. For example, in the case of allowing read access from a specific EC2 instance, when data users request to work on a dataset, a customised policy is generated on the fly based on permissions granted by the DAC.

Once access has been granted, users can analyse the dataset using RAPTOR’s Analytics Workspace. This is an on-demand, dedicated virtual network where all interactions by a user with the chosen data set can occur. The Analytics Workspace comes in three flavours: single node AWS EC2 virtual machine instance, elastic spark cluster and high-performance computing cluster. Through this design, RAPTOR automates the provisioning of selected data, creation and configuration of virtual machines, and the enforcement of security policies, on one computational resource.

Within the EC2 instance, users have full administrator rights by default and can install tools directly from the internet. To save operating costs, workspaces can be shut down when not in use, with the virtual server retaining its mount points, tools, and environment. Only the terminate command will destroy the workspace completely. Users have the option of exporting their virtual machine, the Amazon Machine Image (AMI), to be shared with collaborators. AMIs flagged for export will be reviewed by RAPTOR administrators to ensure the image is safe and does not contain unwanted data. Second, RAPTOR’s elastic spark cluster invokes AWS Elastic MapReduce Service to provision an Apache SPARK cluster (https://spark.apache.org/) with a Zeppelin notebook (https://zeppelin.apache.org/) serving as the front end. This workspace also comes with common genomic analysis tools, such as Hail (https://hail.is/) pre-installed to facilitate data analytics. Third, within the high-performance computing cluster, users can create a dedicated linux computing cluster complete with SLURM scheduler (https://www.schedmd.com/) and a share storage for processing data requiring heavy computational demands.

Workspaces do not create local copies of datasets. The operating system mount points read data directly from where selected datasets are housed on RAPTOR, thereby allowing users to side-step the issue of moving large data files across the network. RAPTOR uses AWS FSX to provision Lustre filesystem (https://www.lustre.org/) for both runtime scratch and analysis outputs. Outputs and results are flushed into S3 for persistent storage when the workspace is shut down. AWS Service Endpoints are used to route function calls to native AWS services. This allows the workspace to function even if all external network access is disabled.

Importantly, the DAC can modify the conditions of data being shared even after RAPTOR has granted user access. For instance, DACs may revoke a user’s permission to modify the original dataset while still allowing the user to access data for specific analysis. Additionally, the DAC may stop all access immediately with a kill switch. i.e., the mounted drive containing the selected data set will immediately disconnect from the analytics workspace. Under Safe Usage, data contributors are thus able to have effective and direct control of datasets housed on RAPTOR.

### Safe User

RAPTOR ensures all users within the system are properly validated and does not allow anonymised access. As part of the user registration process, RAPTOR administrators verify the identity of the applicant. This typically involves running checks with the applicant’s institute or collaborator.

User management and authentication controls in RAPTOR are handled by AWS Cognito due to its compliance with key security standards including ISO 27001, HIPPA BAA and Multi-Tiered Cloud Security (MTCS)^17,18^. RAPTOR does not store user credentials. All user records are encrypted both at rest and in transit. A 2-Factor authentication is mandatory for all users for better protection, which also discourages users from sharing accounts. Additionally, Cognito allows integration with major identity providers, including active directory, and supports protocols including OAuth2, SAML and OpenID Connect^19^, facilitating future integration with systems that conform to GA4GH Authentication and Authorisation Infrastructure^20^.

RAPTOR’s Safe user protocols can also extend to the platform’s core administration and development team. For example, in the case of Singapore, developers and system administrators working on RAPTOR must receive security clearance from Singapore’s Ministry of Home Affairs to work with restricted data. These measures ensure that data deposited in RAPTOR will only be accessed by approved individuals.

### Safe Settings

RAPTOR employs a multi-layer strategy to protect hosted data against unauthorised access, starting with the host infrastructure. RAPTOR is a serverless, cloud native application that leverages CSP with the Platform-as-a-Service (PaaS) model. RAPTOR’s features are constructed from scripts and functions by hosting CSP’s native services (Figure 2). For example, RAPTOR’s graphical user interface is built from a set of Java Scripts hosted on AWS CloudFront, meaning that RAPTOR does not manage or operate any servers. Notably, AWS is a CSP adopted to host public services for the Singaporean government^21^, and was one of the first CSPs to attain MTCS level 3 (Highest level) under SS584:2020, a security standard designed by Infocomm Media Development Authority (IMDA) Singapore^22^, which is required for a CSP to host data and services for the Singapore government. Thus, adopting AWS in RAPTOR for application and data security allows us to provide stringent operating procedures for infrastructure security.

**Figure 2:**
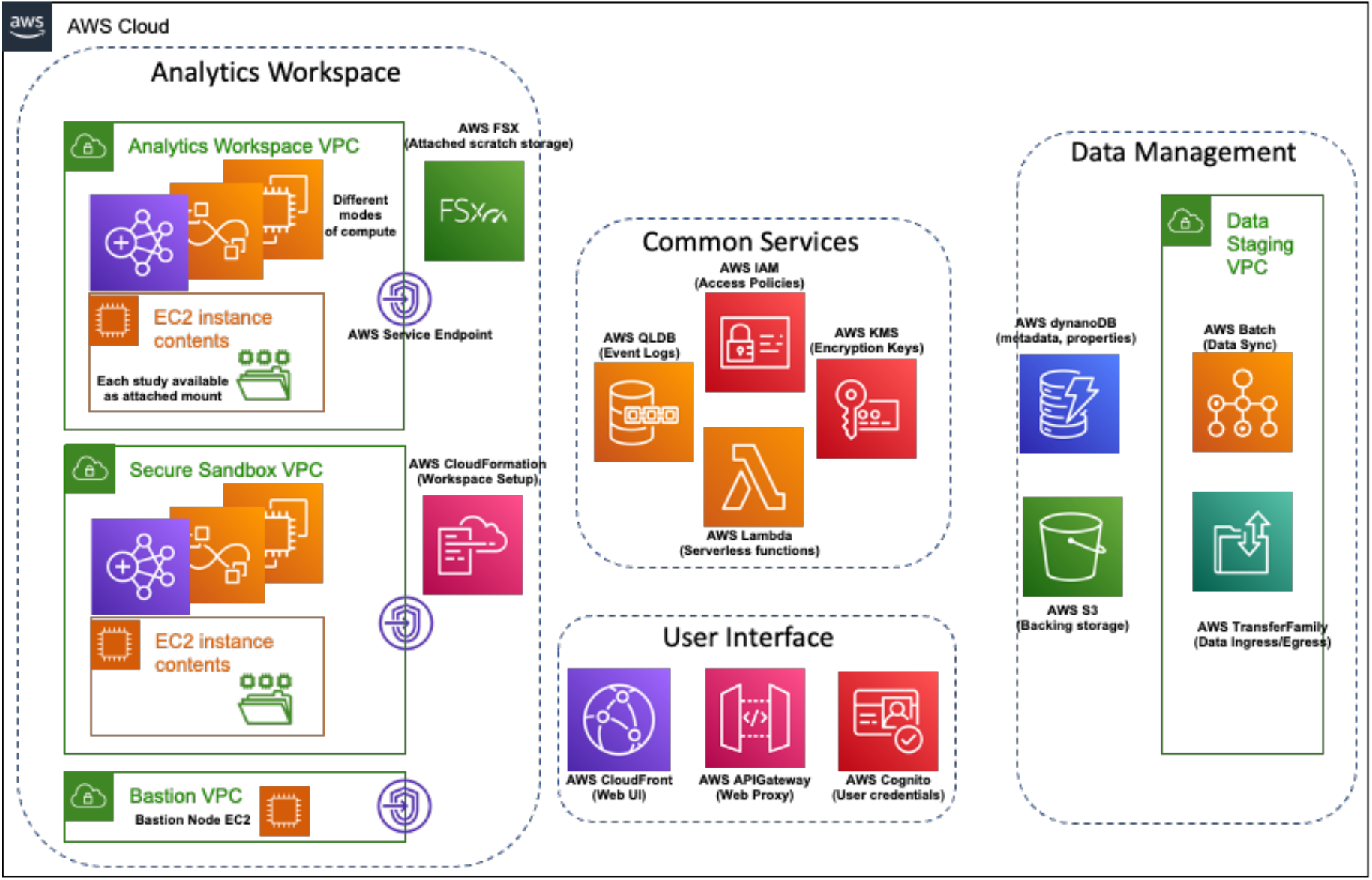
As a serverless application, RAPTOR is composed of native AWS services integrated together with Lambda functions. User Interfaces are composed of CloudFront hosting graphical user interfaces made with java scripts. User authentications are managed with Cognito. Hosted data sets sit on S3 (with automated tiering) while all meta-data are stored on DynamoDB. Data staging activities are managed using S3 Batch. Data ingress and egress are managed through TransferFamily. The Analytics workspace relies on FSX to provide scratch storage and depending on the mode of compute, either EC2, Elastic Map Reduce or Parallel Cluster will provide computing power. Data access from the nodes is regulated by Service Endpoints. All permissions and authorisations are managed using IAM. Encryption keys used by S3, EBS and DynamoDB are stored within KMS. All RAPTOR activities are written into AWS QLDB.

Data encryption provides an added layer of data protection. RAPTOR encrypts all data with symmetric AES256. This extends to data stored on interim server stores within the analytics workspace, dataset metadata and search indexes. To guard against accidental leakage and to provide additional protection against malicious actors, data encryption keys are stored in a separate system, the AWS Key management system (KMS). RAPTOR leverages AWS Parameters Store to encrypt key IDs to prevent accidental leakage of key identifiers when invoking routines. All connections between the user’s computer and RAPTOR are encrypted with SSH2, RC4 or TLS version no older than 1.2.

RAPTOR further provides enhanced data protection for data requiring egress restrictions with the Secure Analytics Workspace (Secure Sandbox). This is a locked-down version of a regular analytics workspace. It provides complete network isolation (no internet), blocking all data egress. A specially provisioned bastion node using Remote Desktop Protocol (RDP) with copy and print redirection disabled is the only way for users to access a Secure Analytics Workspace. Bastion nodes can further restrict its access to whitelisted IPs and subnets.

To validate the effectiveness of RAPTOR’s security measures, RAPTOR undergoes penetration tests and vulnerability assessment by CREST certified assessors at least once every twelve months. During the annual assessment, RAPTOR will be assessed against well-known exploits (published CVEs) and potential weakness in any of RAPTOR’s system dependencies.

Beyond the direct security measures, RAPTOR has an extensive event logging system which tracks all actions performed on datasets, including each time it comes up during a search, when a user submits an access request, and every instance the dataset is provisioned to a workspace. RAPTOR’s logs provide network (TCP/IP) level granularity and are encrypted to protect against tampering. Logs are kept for at least 12 months and can be made available for evaluations upon request.

Taken collectively, RAPTOR ensures that there are adequate mechanisms to protect hosted data, all system vulnerabilities will be promptly patched, and makes available extensive logs to allow reconstruction of events to identify issues or support audits.

### Safe Data and Safe Output

Safe Data and Safe Output refers to RAPTOR’s data protection mechanism for ingress and egress. RAPTOR’s ingress procedure requires the data contributors to deposit their data into a pre-set staging *S3* bucket. Data going into RAPTOR is both screened for malware and checked for authorisation from the project’s DAC before it becomes available for use on RAPTOR. Data hosted on RAPTOR cannot be modified without authorisation from the data contributor. Data on RAPTOR is transparently spread across multiple AWS S3 storage tiers to optimise between cost and access efficiency. The use of AWS S3 also provides data with 11 9s in data durability and 2 9s in data availability^23^. For data contributors who desire higher levels of assurance, options such as data immutability, file level versioning and cross-region backups are available.

To enable egress protection, RAPTOR allows data contributors to mandate the use of Secure Analytics Workspace for users working on their data. To copy files out of a Secure Analytics Workspace, users must submit an egress request on RAPTOR. A copy of the data for egress will be made in a bucket. The requestors will not have any access to this bucket, and requestors cannot change the contents of the files to be egressed after submitting a request. RAPTOR will notify the data contributor of the egress request. The data contributor will then have full file level access to the files RAPTOR had duplicated. The contributor can spin up a “Review workspace” to run content filtering or validation routines on the files submitted. The contributor can subsequently approve the egress request from their data management console and the requestors can export these approved files. If the egress request consists of several datasets that have been combined, all contributors of the respective datasets are required to approve the egress request before the requestor will be allowed to perform data egress.

### Case-study: Secure imputation analysis on RAPTOR

Utilizing population-specific reference panels, during imputation can significantly improve accuracy of low frequency variants (MAF < 1%) for the relevant study population^24–27^. We evaluated the performance of imputing additional genotypes on the Singapore Chinese Health Study (SCHS) dataset (N = 23,600) ^27,28, 29^ and the Asian datasets from the Breast Cancer Association Consortium (BCAC, N = 40,001) using local population-specific reference panels from the SG10K whole-genome sequencing initiative (SG10K Health)^30^ on the RAPTOR platform. Alleles for all SNPs were coded to the forward strand and mapped to HG38. Minimac4 (version 1.0.0) was used to impute variants in the SCHS study using 9,770 local Singaporean population sample reference panels from the SG10K study^31^. Additional imputation on the same SCHS dataset was performed using Trans-Omics in Precision Medicine (TOPMed)^32^ imputation reference panel (version r2) that includes data from 97,256 reference samples (https://imputation.biodatacatalyst.nhlbi.nih.gov). The quality of imputed SNPs from both analyses were determined by impute r^2^ values and high-quality common SNPs (MAF ≥ 1%) were those with an impute r^2^ > 0.3 and high-quality rare SNPs (MAF < 1%) were those with an impute r^2^ > 0.6.

The SG10K data deposited in RAPTOR was flagged as sensitive and thus, the SCHS genotyped data was linked to the SG10K reference panels in the Secure Sandbox. Access to this instance was restricted to a Windows bastion node and users could only connect to the bastion node with the Windows Remote Desktop Protocol (RDP) that has print and clip-board function disabled. Within the bastion node, users will find Secure Shell (ssh) credentials to access the imputation server, with both the SCHS study and the SG10K panels data available as mount points. After the end of imputation, we submitted an egress request for the folder containing imputed dosage data.

### Ancestry Matched Reference Panels Improve Imputation of Rare Variants

We compared the imputation performance in the SCHS after imputing for additional variants using the Trans-Omics in Precision Medicine (TOPMed)^32^ and local SG10K reference panels using the RAPTOR platform. Expectedly, high quality common variants (MAF ≥ 1%) obtained after imputation on the TOPMed and SG10K reference panels were similar (7,236,027 and 7,263,376 bi-allelic SNPs obtained after TOPMed and SG10K imputations, respectively, Table 1). However, a substantially higher number of high-quality rare variants (MAF < 1%) were obtained in the SCHS study through imputation with local SG10K reference panels as compared to the TOPMed imputed data (1,271,426 additional rare SNPs from SG10K imputation procedures, Table 1).

**Table 1:** Numbers of total, common and rare SNPs obtained after imputation of the SCHS dataset with TOPMed and SG10K imputation panels.

|  | SCHS (N=23,756) |  |
| --- | --- | --- |
|  | TOPMed<br>Imputation | SG10K<br>Imputation |
| Total SNPs obtained after imputation | 271,221,018 | 42,602,074 |
| Common SNPs (MAF ≥ 1%) with high imputation quality ($r^2 > 0.3$) | 7,236,027 | 7,263,376 |
| Rare SNPs (MAF < 1%) with high imputation quality ($r^2 > 0.6$) | 9,496,227 | 10,767,653 |

### Five Safes Framework enabled efficient usage of consortia level data in RAPTOR

BCAC data consisted of South Asian and South-East Asian ancestry subjects that were genotyped on the Infinium OncoArray (N = 27, 501) and the iCOGS array (N = 12,500). Data was highlighted as sensitive and similarly required mandatory use of the Secure Sandbox, as well as requiring limited usage for only imputation protocols. The BCAC study folder was linked to the SG10K reference panels in the Secure Sandbox and access to this instance was similarly restricted to a Windows bastion node paired exclusively to the instance running imputation service. Of note, even on the a single AWS EC2 instance (r5.8xLarge), this entire work for imputation of the relatively large BCAC dataset (N = 40,001) in a secure setting, took only around 20 days (end-to-end), and incurred modest costs of about USD 3,000 worth of AWS utilisation, highlighting that the RAPTOR platform enables for effective and secure usage of large-scale consortia level data.

### Remote Data Retrieval Using GA4GH Standards

Data integration with other data platforms, such as combining RAPTOR’s genomic data with another repository holding phenotype information, is a key feature of RAPTOR. GA4GH work streams have defined several standards and APIs to facilitate data federation across different genomic repositories. The first task in any data exchange is to discover and list datasets available in the remote host. This is most effective when performed before initialising user authentication and authorisation. Hence common standards are crucial in enabling two different data sources to exchange “data catalogues” securely without authentication. The GA4GH Discovery workstream’s Beacon V2 standard^33^ provides an efficient mechanism for such activity. We do not anticipate major roadblocks adding this functionality to RAPTOR as the Discovery workstream has provided a working reference for implementation of the Beacon v2 standard. Furthermore, as RAPTOR is serverless, every conceptual “layer” of the application is exposed via APIs and a thin integration layer between the Beacon v2 reference implementation and RAPTOR’s hosting services can be readily incorporated. The integration layer will serve to translate functions and calls from the Beacon v2 implementation into native service calls for RAPTOR, allowing RAPTOR to reuse the Beacon v2 reference implementation with minimal modifications. The GA4GH Data Repository Service (DRS)^34^ is a set of APIs providing consumers (both users and workflows) with direct access to data in a repository. A key feature of DRS is the provision of a Universal Resource Identifier (URI) to provide an exclusive identification for a single file or group of files within a repository. This will allow data consumers to access resources without prior knowledge of repository’s data organisation or file hierarchy. RAPTOR also enables data contributors to define groups of files or directories termed a Collection. RAPTOR Collections can be shared and referenced by data consumers independently of Collection’s parent data set. Therefore, it may be possible to extend RAPTOR Collections to a DRS compliant resource. DAC approval can be integrated into authentication and authorisation workflows before retrieving data using DRS. Destination endpoint IP address for remote retrieval can also be whitelisted.

### Federated analysis with Data-In-Place

Even with DRS approval and IP whitelisting, remote data retrieval may inevitably weaken RAPTOR’s Safe Purpose assurance. In addition, when working with data sets on the scale of multiple terabytes or more, cost and latency for replicating large data volume across networks can quickly become prohibitive. A more feasible approach, would be to send the compute job to the data and return outputs of the job. GA4GH has defined three standards for sending compute jobs to be executed on remote sites, including the Tools Registry Service (TRS), Workflow Execution Service (WES) and Task Execution Service (TES)^35^. TRS provides standardization for tools discovery, providing standardize descriptions of docker-based tools and popular workflow engines^36^. Dockstore (https://dockstore.org/) is an example of a TRS compliant container repository. WES then build on top of TRS to provide a common interface to interact with TRS tools and workflows. TES is similar to WES in that it also defines an interface defining and running compute tasks^37^. TES is differentiated from WES in that a TES task can be modelled as a single job execution such as executing a single script or a command, while a WES is designed for executing a pre-composed workflow. It is possible for a TES to be “nested” within a WES. For example, a Nextflow workflow can serve as a TES client submitting tasks to a TES server^38^.

These three services, however, are insufficient for RAPTOR to provide federation services to third party repositories. As a serverless application, RAPTOR does not have any ready virtual servers to execute remote workflows or tasks requests. The Analytics Workspace is also not suitable as it is designed for direct interactions with end-users and would be unwieldy to automate. In addition, these standards do not provide an easy way to inform RAPTOR of the type of virtual machines required for execution. As an example, while TES allows for resource specification within a task request, the resource request is defined in the form of cores, ram, disk size and name or URI of a docker container. These parameters alone are not sufficient to identify an instance type. Specifically, on AWS, just specifying “2 cores and 8 GB ram” maps to at least 7 different instance types, each with a unique hardware profile, optimised for different use-case. There is also no means of specifying a type of platform (x86 or ARM, Memory optimised, or GPU enabled), and loading containers or images from third party repository is not compatible with RAPTOR’s operations. To preserve RAPTOR’s Safe Purpose assurance, RAPTOR administrators would have to manually clear an AMI or container before allowing it for use, which would require the container or image to be hosted within a restricted repository trusted by RAPTOR administrators. The remote user would thus have to perform an “out-of-band” communication with the data contributor on RAPTOR to learn the identifier of the AMI or container to be referenced with TES. This may complicate workflow scripting or automation.

To overcome these limitations, RAPTOR’s development team evaluated an early concept where we could pre-define and associate an AMI with a data set. AMIs allow users to define key features for a virtual server, such as platform configured tools (x86 or ARM, ROCm or CUDA) and even reference files. At the same time, AMIs provide some flexibility for users to determine the appropriate sizing (core counts, amount of ram, volume of attached disks) during runtime. The exchange only involves the universal identifier of the AMI, not the actual image file. When a remote compute request has been received, RAPTOR will initialise a virtual machine using AWS Batch, with core count and ram size matching the values from TES. Batch will initialise the virtual machine, execute the command, write-back the output to requestor and terminate the machine.

### Case-study: prototype of federated analysis on RAPTOR

As a case study, the RAPTOR development team recognized the challenge of performing these activities within the defined set of GA4GH APIs. We thus implemented an early-stage proof of concept where a RAPTOR instance received a remote computation request together with an AMI and docker identifier, with task execution and termination of the virtual server after the results were returned.

In the prototype, we evaluated a federated imputation workflow in RAPTOR by performing imputation on a Primary Open Angle Glaucoma dataset (POAG) using Minimac4 (version 1.0.0) with the same SG10K health reference panels. However, in this scenario, two RAPTOR instances were created, one hosting POAG and the other SG10K health panels. Each customised instance had added functions, providing a minimal implementation of TES and task invocation via the API Gateway and Batch. The team limited the POC’s scope to TES, since the initial DRS exchange only involved inserting additional data content under “alias” section of response to a DRS “Get” call. Hence, we configured the RAPTOR instances assuming that information can be packed within the single DRS get call. Specifically, these include the AMI id, minimac4 imputation commands, mount paths and customised AWS IAM role (Role created for this workflow only, will be destroyed after the end of workflow) and egress IP address. The IAM role and IP were used to configure POAG RAPTOR to allow NFS read-write mounts from the SG10K RAPTOR. This allowed SG10K RAPTOR to read and write data to POAG RAPTOR.

The complete imputation workflow could then be broken down into four main TES tasks, 1. Creating unphased VCFs, 2. Conversion to phased VCFs, 3. Addition of prefix “chr” to entries in phased VCFs and 4. Imputation. During the evaluation, the POAG RAPTOR submitted the four TES tasks creation calls in sequence. Each task was invoked after the previous one had been completed successfully (validated via TES “Get task” call). With every task received, the SG10K RAPTOR will submit a job to AWS Batch service using its account with a predefined Virtual Private Network (VPC) using parameters received from TES calls. To allow reading both data sets during analysis and writing of outputs to either (or both) of RAPTOR instances, a virtual machine was set-up to run FUSE mounts to both POAG RAPTOR and SG10K RAPTOR. AWS Batch service will run the TES tasks and write outputs back into POAG RAPTOR. After the end of an AWS Batch job, AWS Batch Service will automatically stop and delete all resources associate with the task (Except files written to output directories). Users on POAG RAPTOR can monitor the task status by issuing TES “Get task” calls to SG10K RAPTOR. Once a task was completed, the user will submit the subsequent TES task until all 4 tasks are completed (figure 3).

**Figure 3:**
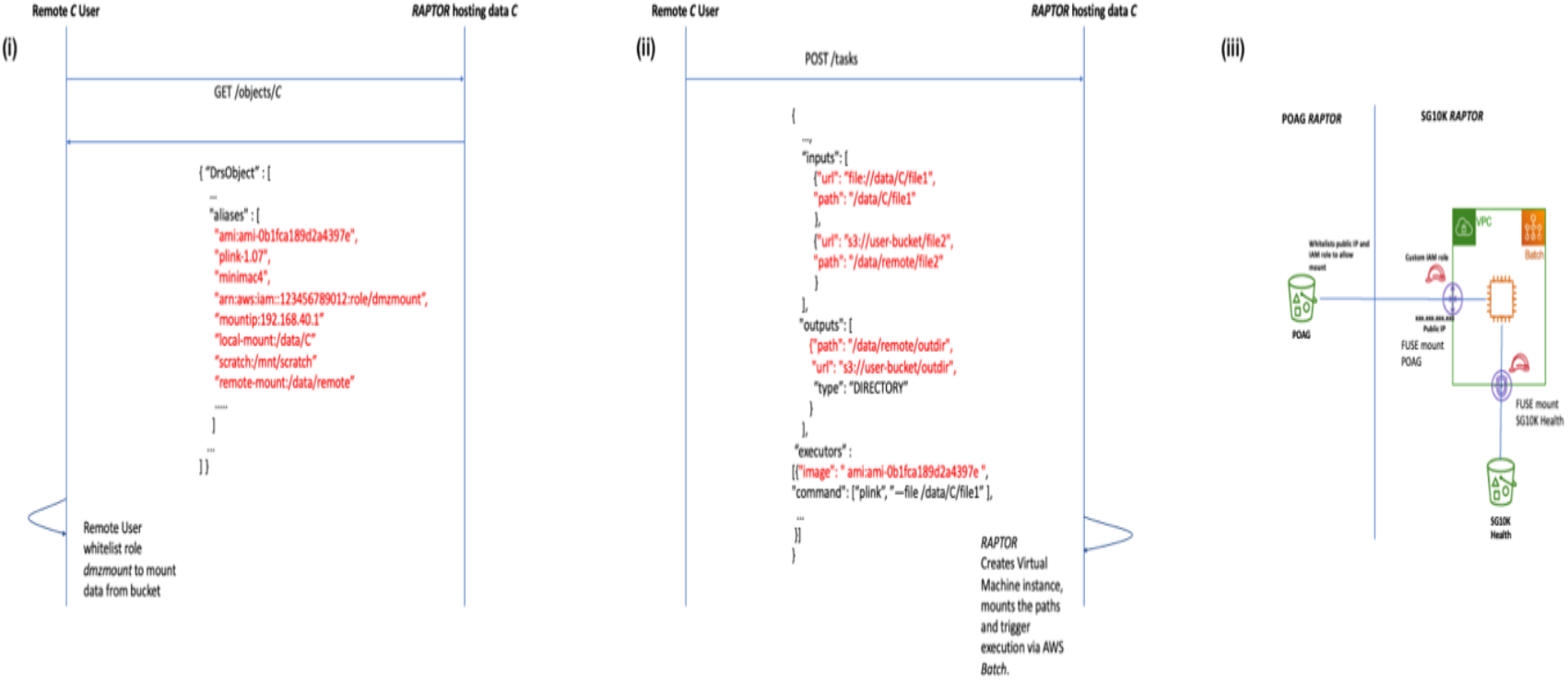
Using existing GA4GH DRS and TES for federated computation on RAPTOR. (i) A client invokes Get to describe an existing data collection ‘C’ on RAPTOR. In addition to content and access descriptions, RAPTOR also sends identity of the AMI associated with C under ‘alias’. The provided information includes AMI id, pre-installed tools, mount points for accessing data and the customised IAM role and end-point IP – these are used by RAPTOR to read and write data from the remote client. (ii) The remote client invokes a task using TES. In the call, inputs are used for the client to inform RAPTOR which URI is to be mapped to which mount point. Under executors, the remote client will inform RAPTOR which AMI is to be used. (iii) Batch instantiates an EC2 machine AMI and parameters provided by TES task. The machine will mount paths from POAG RAPTOR and SG10K RAPTOR within the same machine using the customised IAM role.

Using the POC system, we completed the imputation of POAG chr21 with SG10K Health. Results from the POC system were consistent with results from work done on our production RAPTOR (Whole-genome imputation of POAG with SG10K Health panel). With this approach, the “host” site controls the tools, security policies and the virtual machines, while the remote user directs where the outputs are written. The use of *Batch* ensures all interim data in scratch will be deleted. Hence, we believe this protype approach can provide both the remote user and the host strong protections against data leaks.

## Discussion

### Value of RAPTOR-like platforms for genomics research

A cornerstone to precision and personalised medicine is the ability to effectively access and thoroughly evaluate a multitude of genomic datasets to improve our molecular understanding of health and disease processes. Managing large scale data analysis across multiple datasets, however, remains challenging. Hyper-scale computational capabilities provided by CSPs such as AWS provide a full suite of computational resources to enable concurrent in-data-analysis data on an AWS S3 Bucket. Nevertheless, harnessing these resources requires knowledge on programming and cloud computing expertise. Additionally, the sensitive nature of genomics data and increased emphasis on data governance and security limits capabilities of individual labs to setup a cloud platform for effective large-scale genomic collaboration studies. Data contributors from public sectors also routinely request for specific controls to be put in place, such as AES256 symmetric for data encryption at rest and the version of TLS/SSH protocols for data encryption during transit. Data hosting facilities may also require certifications such as ISO 27001 or MCTS Level 3.

RAPTOR provides users with a platform which addresses these computation and security concerns. Through RAPTOR, users seamlessly run their analysis as they would have on an on-premise system or their private cloud machine while, still staying compliant to overarching governance and security regulations. Our case example on performing imputation on large-scale and consortia level datasets demonstrates these capabilities in RAPTOR and highlights the potential of integrating various genetic resources to improve genetic studies, while, at the same time, remaining compliant to required restrictions of sensitive genetic data.

Beyond security, datasets hosted on RAPTOR become FAIR (Findable, Accessible, Interoperable and Repeatable)^39^. RAPTOR allows contributors to add meta-data as tags to their data sets, allowing for quick way to filter out relevant data sets using RAPTOR’s simple search form (Example: Show all data sets where “Country = Singapore and ethnicity = Malay”). RAPTOR users can submit request to access any dataset hosted on RAPTOR, with levels of access determined by the respective DACs. As RAPTOR manages all data provisioning tasks, users can start working on the requested data sets immediately upon receiving approval. RAPTOR’s AMI sharing feature further allows users to save and share their intermediate data and work environment (i.e. Juypter-style Notebook or even the whole Linux machine) with other RAPTOR users, enabling interoperability and repeatability.

RAPTOR went live in 2021 and has uniquely built a collection of Asian genetic SNP array-based genetic datasets that are primarily from the local Singapore ethnic population groups (Singapore Chinese, Malay, and Indian datasets). As of October 2022, RAPTOR is hosting close to 100,000 samples with more than half from Singapore (Table 2).

**Table 2:** Datasets hosted on RAPTOR. # denotes studies with samples from Singapore.

| **Studies** | **N** | **Ethnicity** | **Data available** |
| --- | --- | --- | --- |
| Singapore Chinese Health Study<sup>#</sup> | 23,756 | Chinese | GWAS Array / CNV data |
| SCHS Coronary Artery Disease Cohort<sup>#</sup> | 2,003 | Chinese | GWAS Array |
| Singapore Chinese Eye Study<sup>#</sup> | 1,889 | Chinese | GWAS Array |
| Singapore Malay Eye Study<sup>#</sup> | 2,542 | Malay | GWAS Array |
| Singapore Indian Eye Study<sup>#</sup> | 2,538 | Indians | GWAS Array |
| Singapore Coronary Artery Disease Genetics Study<sup>#</sup> | 1,943 | Chinese, Malay, and Indians | GWAS Array |
| Study of Macro-angiopathy and Micro-vascular Reactivity in Type 2 Diabetes (SMART2D) dataset <sup>#</sup> | 1,893 | Chinese, Malay, and Indians | GWAS Array |
| Diabetic Nephropathy dataset <sup>#</sup> | 2,563 | Chinese, Malay, and Indians | GWAS Array |
| SG10K Health r5.5 <sup>#</sup> | 9,770 | Chinese, Malay, and Indians | Imputation Reference Panel |
| Jerusalem Perinatal study | 2,593 | Israeli | GWAS Array |
| Singapore Prospective Study Program <sup>#</sup> | 2,434 | Chinese | GWAS Array |
| Primary Open Angle Glaucoma Study<sup>#</sup> | 3,580 | Chinese | GWAS Array |
| Singapore Cohort Of the Risk factors for Myopia <sup>#</sup> | 1,029 | Chinese | GWAS Array |
| Breast Cancer Association Consortium | 40,001 | Pan Asian | GWAS Array |

In the long-term, platforms such as RAPTOR are likely to represent options as trusted custodians for national-scale genomic data. RAPTOR’s next phase will also focus on expanding the types of data hosted on the platform. Various transcriptomic datasets from local population datasets are expected to be housed in RAPTOR, facilitating larger-scale transcriptomic studies and combinatorial expression quantitative trail loci studies. To do so, RAPTOR will continuously update security measures and operating procedures to ensure compliance with relevant data and IT governance policies. While we have not extensively reviewed RAPTOR’s alignment to similar policies from other jurisdictions, we note that RAPTOR’s core features facilitates alignment to 5 of the 7 core principles of European Union’s data protection law (GDPR)^40^. These include lawfulness, fairness and transparency and purpose limitation by providing the DAC with control over how data is used and who can access data, data minimisation where data contributors may create cub-collections of data for sharing, integrity and confidentiality through RAPTOR’s security capabilities and encryption adoptions as well as accountability through the storage of logs of every activity on RAPTOR. The two remaining principles on Accuracy and Storage limitation are perhaps, beyond the scope for a data and analysis platform such as RAPTOR and may be adopted by the data contributor and DAC.

### Federated analysis and data sharing using GA4GH protocols

For RAPTOR, the ability to integrate with other repositories (either via data movement or federated analysis) will be essential for the next phase of development. GA4GH protocols will be the key interface between RAPTOR and other repositories and platforms. The key challenge, as highlighted, is ensuring this implementation continues while retaining the 5 Safes assurances. One current limitation in our study was that for our DRS implementation, there required be a mandatory, out-of-band approval seeking with dataset’s DAC before a data set can be retrieved to remote site. Notably, while DRS’s standards require the use of OAuth2 tokens for authentication, the standard is silent on the authentication and authorisation procedures. RAPTOR is already utilising OAuth2 for user authentication. RAPTOR’s authentication mechanism can be extended to applications and scripts (i.e., automated flow triggered by user or software), with IP whitelisting as replacement for 2FA. However, once data has egressed, there would be no effective way for DAC to track subsequent usage or enforce additional compliances. It should, however, be noted that RAPTOR Data marked as sensitive (mandatory use of secure analysis workspace) will not be valid for retrieval using DRS.

We propose that RAPTOR’s federated analysis will provide stronger adherence to the 5 Safes. The federated analysis environment and tools are pre-determined by the data contributors (i.e., the remote data site) and are immutable. All endpoints are IP-locked, with the pre-determined write out process. The data owners therefore have full control of how the data is to be used and the outputs which will be shared with the remote user. Data access expiry can be automatically enforced by RAPTOR.

RAPTOR’s federated analysis is implemented with GA4GH DRS and TES API interfaces. RAPTOR utilises free text sections within the DRS and TEST schemas to exchange essential integration information, including the data paths and tools available for use. The goal of using DRS and TES APIs is not to allow “ad-hoc” invocations from remote users. The goal of using GA4GH APIs is mostly to reduce the implementation and customisation overheads.

### RAPTOR’s technical advantage and roadmap

A key advantage from building RAPTOR as a “serverless” application is that RAPTOR is completely abstracted from both the system hardware and software, including the operating system. Instead, core RAPTOR services including the user interface and user management are plugged directly into AWS’s hyper-scale compute and storage fabric. All button clicks and function calls are distributed to a managed cluster of servers that ensures consistent performance that scales with load while keeping the cost of upkeep low, since RAPTOR is billed only for the resources for the functions executed, not for the number of servers it ran on. Apart from cost efficiency and performance scaling, a serverless resource also provides an easier pathway for integration with existing applications and services. The reason is that major CSP such as AWS reduces entire system infrastructure as a service to a series of functions calls that are by design consistent with cloud native service calls utilised for core RAPTOR functions. This means it is programmatically similar for us to deploy a complete system of applications as to implement a button on the user interface, hence appreciably reducing the effort and time required to integrate 3^rd^ party applications. For example, we have plans to enhance hosted data’s findability by providing improvements on metadata capture and adopting GA4GH’s Data Use Ontology (DUO)^41^. However, instead of building more native services, we are evaluating ways to integrate with 3^rd^ party tool suites such as those from Centre for Expanded Data Annotation and Retrieval (CEDAR)^42^. We will package the entire system into a software-defined infrastructure that will be provisioned on-demand.

Based from learnings from this prototype, the next developmental goal for RAPTOR is put to implement a feature complete data exchange with third party data sources (data federation) using combinations of community defined standards, including GA4GH Beacon v2, Data Repository Service (DRS) and WES.

In conclusion, we propose that the RAPTOR computational platform provides researchers a significant resource that enables power of cloud-based computing while, ensuring a safe and secure environment to meet regulatory requirements for genomic data. Additionally, RAPTOR enables flexibility in data federation and offers potentials to integrate multiple data repositories that would enable for more effective analysis of large-scale genomic data.

## Acknowledgements

We would like to thank Charissa Chang of AWS Singapore, Kenny Max, and Smit Callum of AWS Envision Engineering. RAPTOR grew out of an early prototype Kenny and Smit built on top of AWS Service Workbench. Charissa secured technical support from AWS that was invaluable for RAPTOR to transit from proof-of-concept to an operational platform.

The Singapore Chinese Health Study was supported by grants from the National Medical Research Council, Singapore (NMRC/CIRG/1456/2016 and NMRC/CSA/0055/2013), the Saw Swee Hock School of Public Health, National University of Singapore, and the USA National Institutes of Health (R01 CA144034 and UM1 CA182876). The Singapore Coronary Artery Disease Genetics Study (SCADGENS) and genotyping of the SCHS-CAD cohort was supported by the HUJ-CREATE Programme of the National Research Foundation, Singapore (Project Number 370062002). The SCORM study is supported by the National Medical Research Council Grant NMRC/0975/2005. The Jerusalem Perinatal study was supported by Israeli Science Foundation grants 201/98-1 and 407/17 and by National Institutes of Health research grant R01HL088884. The SP2 study is supported by individual research and clinical scientist award schemes from the National Medical Research Council (NMRC) and the Biomedical Research Council (BMRC) of Singapore, the Singapore Ministry of Health, National University of Singapore and National University Health System, Singapore. The Singapore Study of Macro-angiopathy and Micro-vascular Reactivity in Type 2 Diabetes (SMART2D) cohort was supported by grants from the National Medical Research Council, Singapore (NMRC/PPG/AH(KTPH)/2011 and NMRC/CIRG/1398/2014). The Diabetic Nephropathy (DN) cohort was supported by grants from Alexandra Health Fund Private Limited (SIG II/15205). The Singapore POAG study was supported by a Biomedical Research Council (BMRC) grant in Singapore. This research was also partly supported by a grant (NMRC/TCR/008-SERI/2013) from the Singapore National Research Foundation under its Translational and Clinical Research Flagship Programme and administered by the Singapore Ministry of Health’s National Medical Research Council.

This study made use of data generated as part of the Singapore National Precision Medicine program funded by the Industry Alignment Fund (Pre-Positioning) (IAF-PP: H17/01/a0/007). We also acknowledge the following source of funding support for recruitment and genotyping of population-based cohorts SIMES and SCES: National Medical Research Council, Singapore (NMRC/TCR/002-SERI/2008, (R626/47/2008TCR), CSA R613/34/2008, NMRC 0796/2003, STaR/0003/2008), the National Research Foundation of Singapore, the Biomedical Research Council, Singapore (BMRC 09/1/35/ 19/616 and 08/1/35/19/550) and Genome Institute of Singapore.

This study made use of data / samples (Request: NPM00038) collected in the following cohorts in Singapore:

1. The Health for Life in Singapore (HELIOS) study at the Lee Kong Chian School of Medicine, Nanyang Technological University, Singapore (supported by grants from a Strategic Initiative at Lee Kong Chian School of Medicine, the Singapore Ministry of Health (MOH) under its Singapore Translational Research Investigator Award (NMRC/STaR/0028/2017) and the IAF-PP: H18/01/a0/016);
2. The Growing up in Singapore Towards Healthy Outcomes (GUSTO) study, which is jointly hosted by the National University Hospital (NUH), KK Women’s and Children’s Hospital (KKH), the National University of Singapore (NUS) and the Singapore Institute for Clinical Sciences (SICS), Agency for Science Technology and Research (A*STAR) (supported by the Singapore National Research Foundation under its Translational and Clinical Research (TCR) Flagship Programme and administered by the Singapore Ministry of Health’s National Medical Research Council (NMRC), Singapore - NMRC/TCR/004-NUS/2008; NMRC/TCR/012-NUHS/2014. Additional funding is provided by SICS and IAF-PP H17/01/a0/005);
3. The Singapore Epidemiology of Eye Diseases (SEED) cohort at Singapore Eye Research Institute (SERI) (supported by NMRC/CIRG/1417/2015; NMRC/CIRG/1488/2018; NMRC/OFLCG/004/2018);
4. The Multi-Ethnic Cohort (MEC) cohort (supported by NMRC grant 0838/2004; BMRC grant 03/1/27/18/216; 05/1/21/19/425; 11/1/21/19/678, Ministry of Health, Singapore, National University of Singapore and National University Health System, Singapore);
5. The SingHealth Duke-NUS Institute of Precision Medicine (PRISM) cohort (supported by NMRC/CG/M006/2017_NHCS; NMRC/STaR/0011/2012, NMRC/STaR/ 0026/2015, Lee Foundation and Tanoto Foundation);
6. The TTSH Personalised Medicine Normal Controls (TTSH) cohort funded (supported by NMRC/CG12AUG17 and CGAug16M012).

The views expressed are those of the author(s) are not necessarily those of the National Precision Medicine investigators, or institutional partners. We thank all investigators, staff members and study participants who made the National Precision Medicine Project possible.

## Abbreviations

AAI: Authentication and Authorisation Infrastructure
AMI: Amazon Machine Image
API: Application Programming Interface
AWS: Amazon Web Services
BAA: Business Associate Agreement
CCP: Commercial Cloud Platform
CREST: Council of Registered Ethical Security Testers
CSP: Cloud Service Provider
CVE: Common Vulnerabilities and Exposure
DAC: Data Access Committee
DRS: Data Repository Service
DUO: Data Use Ontology
EBS: Elastic Block Storage
EC2: Elastic Computing Cloud
EMR: Elastic Map Reduce
GA4GH: Global Alliance for Genomics and Health
GCP: Google Cloud Platform
HIPPA: Health Insurance Portability and Accountability Act
IAM: Identity Access Management
IMDA: Inforcomm Media Development Authority
KMS: Key Management System
MAF: Minor Allele Frequency
MTCS: Multi-Tiered Cloud Security
POAG: Primary Open Angle Glaucoma
QLDB: Quantum Ledger Database
RAPTOR: Research Assets Provisioning and Tracking Online Repository
RDP: Remote Desktop Protocol
S3: Simple Storage Service
SCHS: Singapore Chinese Health Study
SSH: Secure Shell
TCP/IP: Transmission Control Protocol/Internet Protocol
TES: Task Execution Service
TOPMed: Trans-Omics in Precision Medicine
URI: Universal Resource Identifier
VPC: Virtual Private Cloud
WES: Workflow Execution Service

