## Supplementary for "RAPTOR: A Five-Safes approach to a secure, cloud native and serverless genomics data repository"

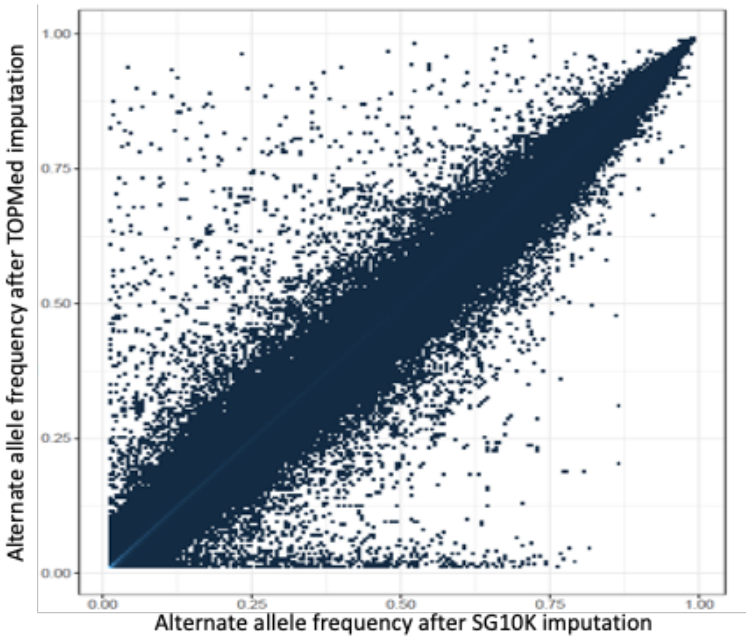


Supplementary figure 1: Strong correlation (r=0.9986) between alternate allele frequencies for common variants (MAF ≥ 1%) imputed in the SCHS study using TOPMed and SG10K reference population panels.

### On Access to Data Sets on *RAPTOR*

#### Request for Access

1. Go to the **Studies** tab

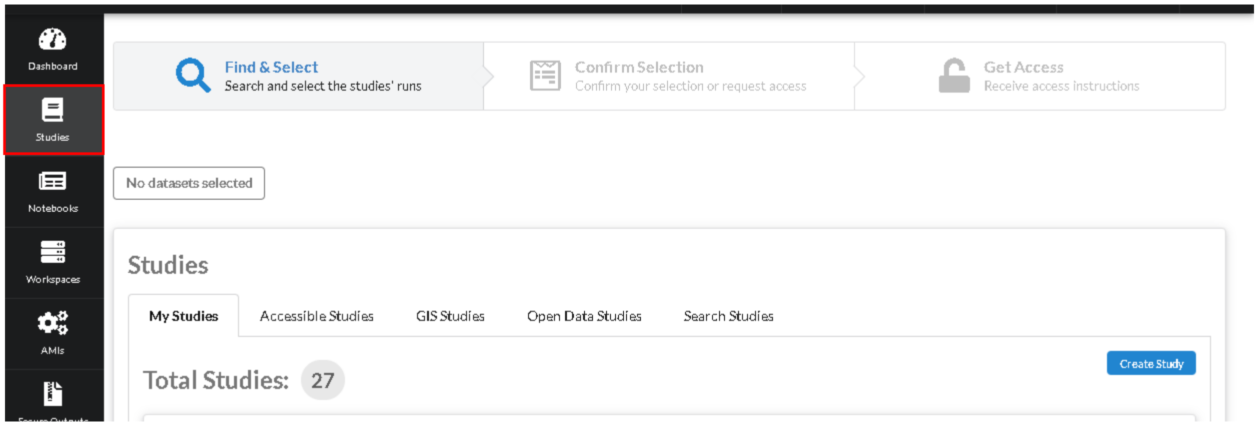


2. Under **GIS Studies** tab, click on **Request Access** button of the study you would like access to.

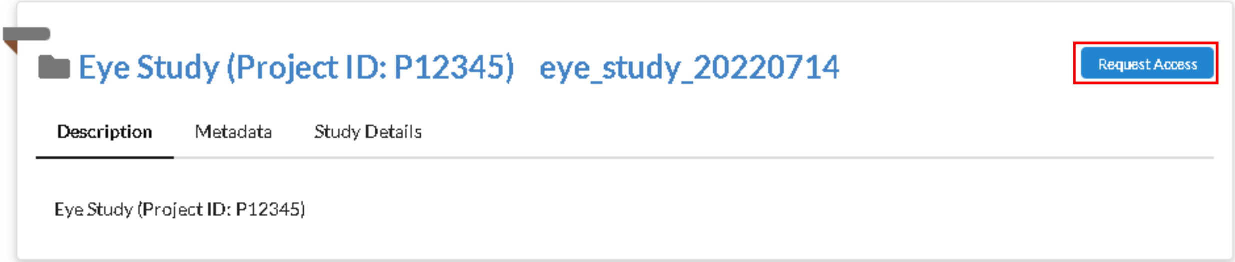


3. Specify your access details and click **Request Access**.

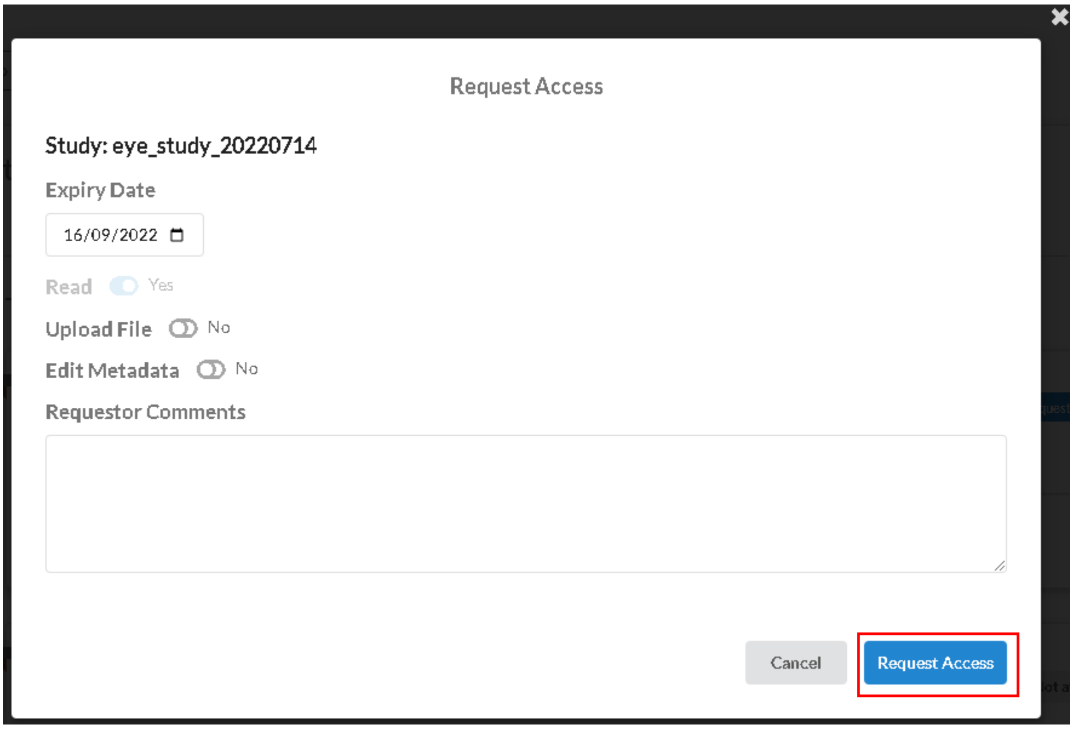

#### Approve/Reject Study Access Requests

Please note that you need to be an admin of the study to perform this action.

1. Go to the **Access Requests** tab.

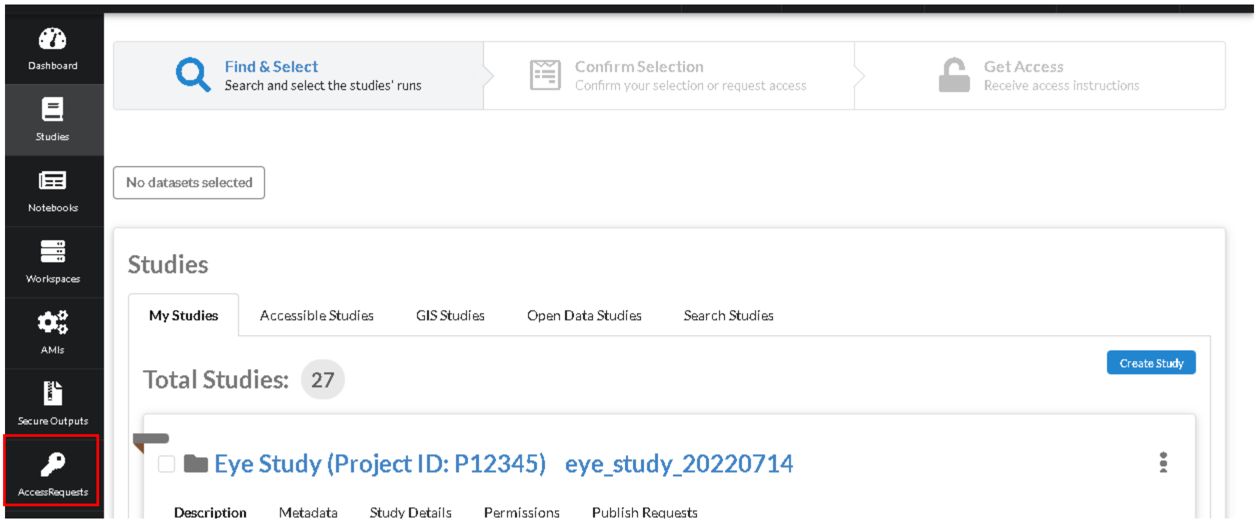


2. Navigate to the study. Click on the corresponding buttons to approve/deny the request.

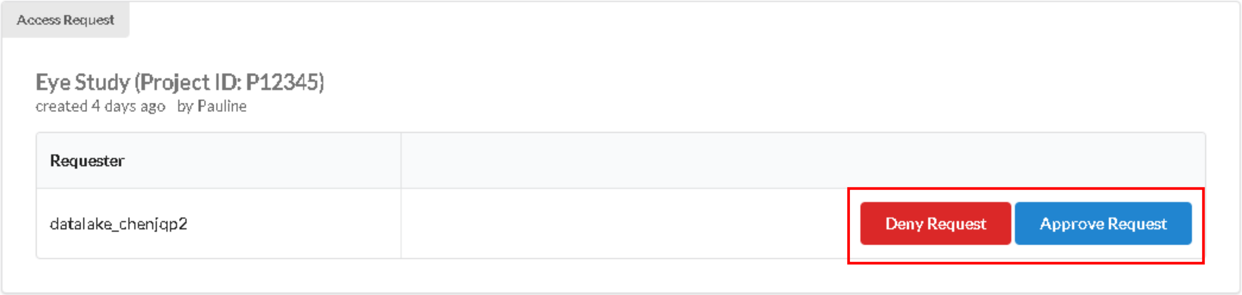

3. On clicking **Deny Request**, you will be presented with a field to provide your comments on rejection (compulsory). Once ready, click **Deny Request**.

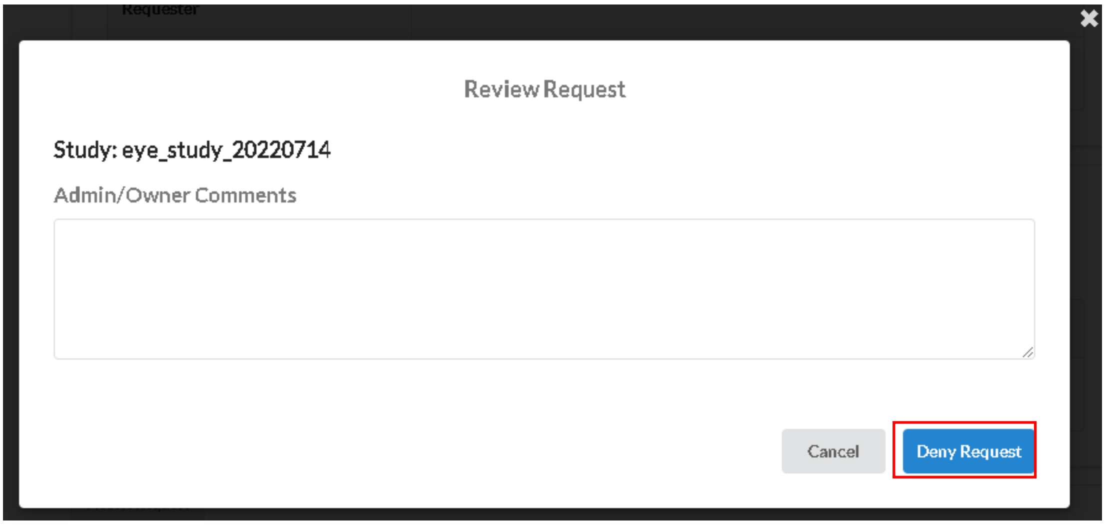


4. On clicking **Approve Request**, you will be presented with the request details and a field to provide your comments on approval (compulsory). Once ready, click **Approve Request**.

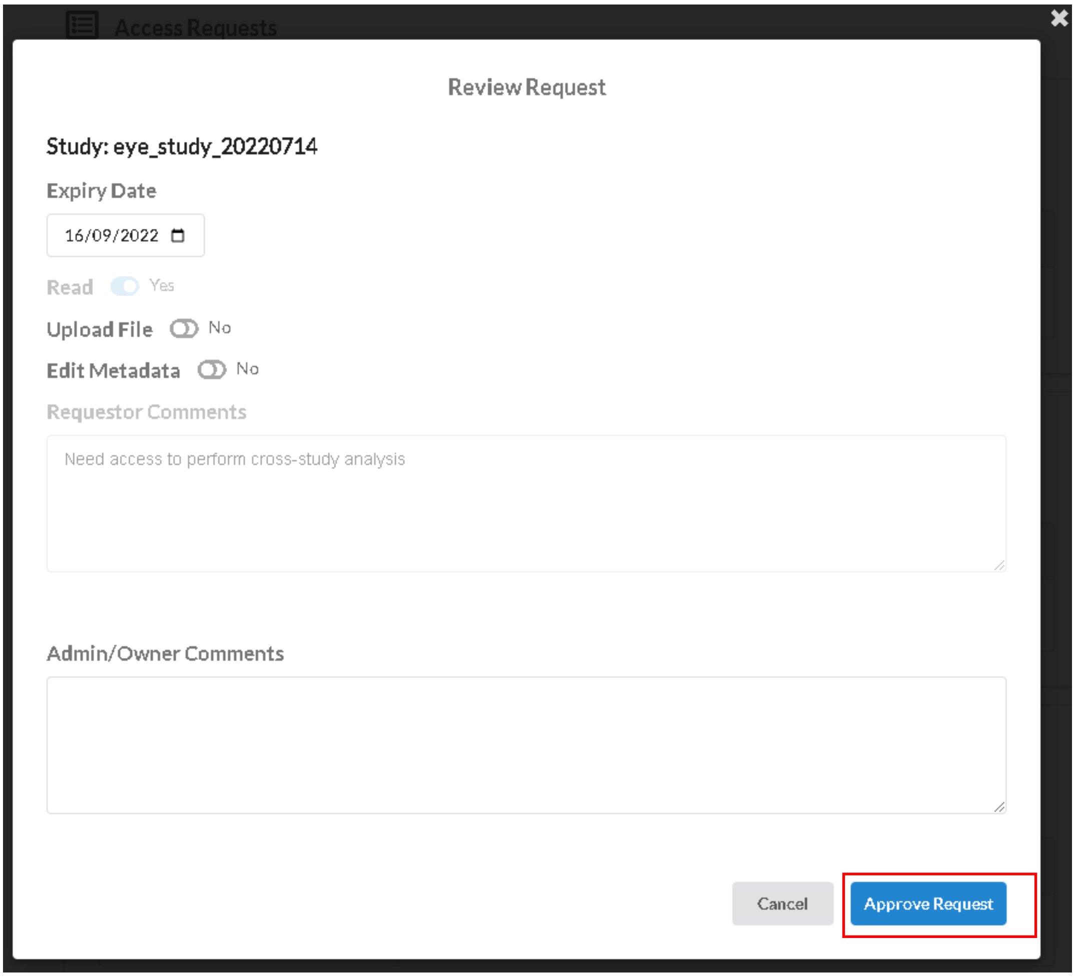


#### Update Study Public Status

Please note that you need to be an admin of the study to perform this action.

1. Go to the **Studies** tab

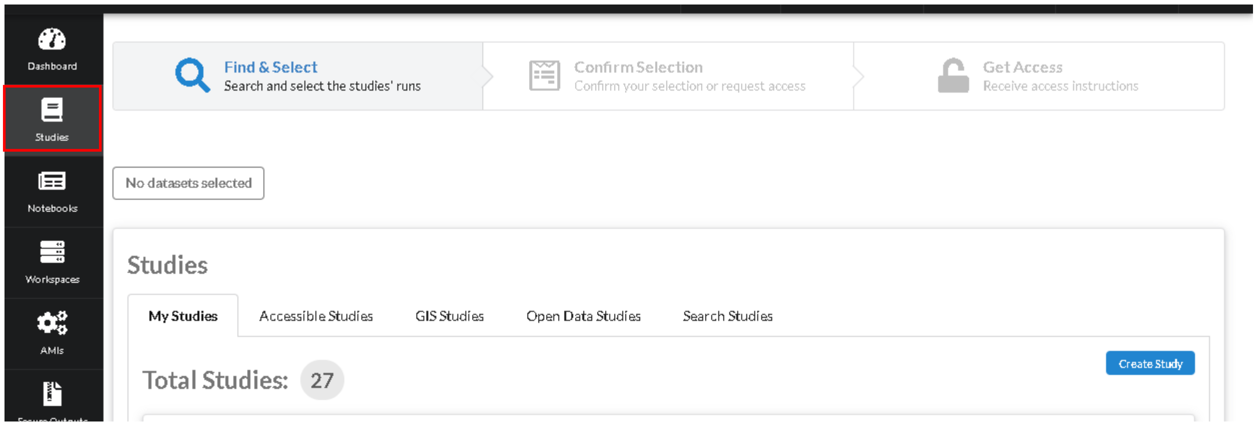


2. Under **My Studies** tab, locate the study card. Click on **Permissions** sub-tab, then click on the pencil icon beside **Users**.

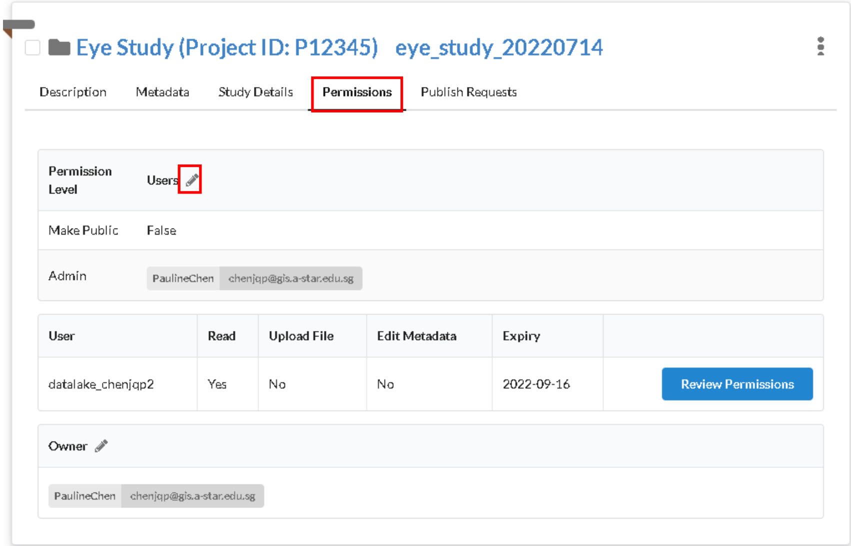


#### Changing permissions for an existing data set

Please note that you must be an admin of the study to perform this action.

##### Delegating project admin

1. Go to the **Studies** tab

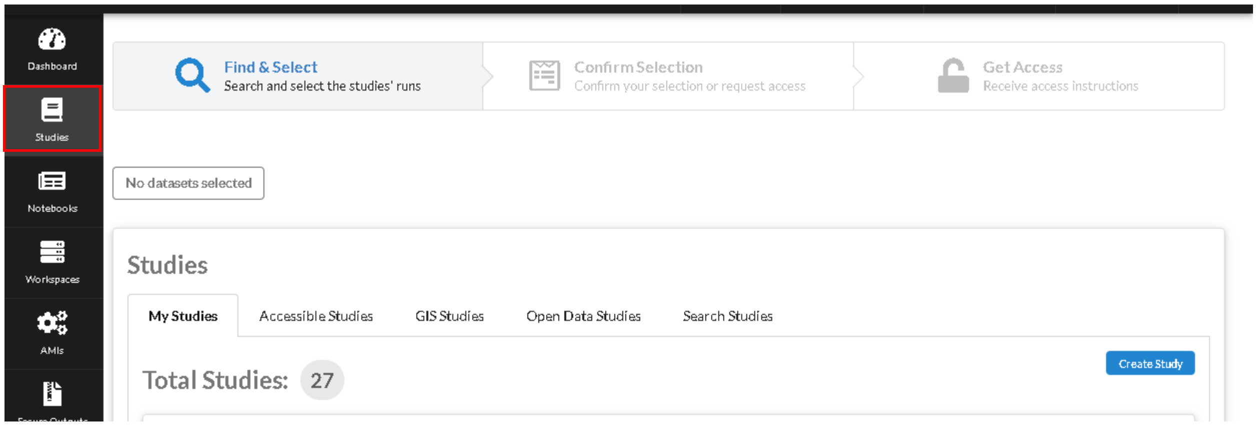


2. Under **My Studies** tab, locate the study card. Click on **Permissions** sub-tab, then click on the pencil icon beside **Users**.

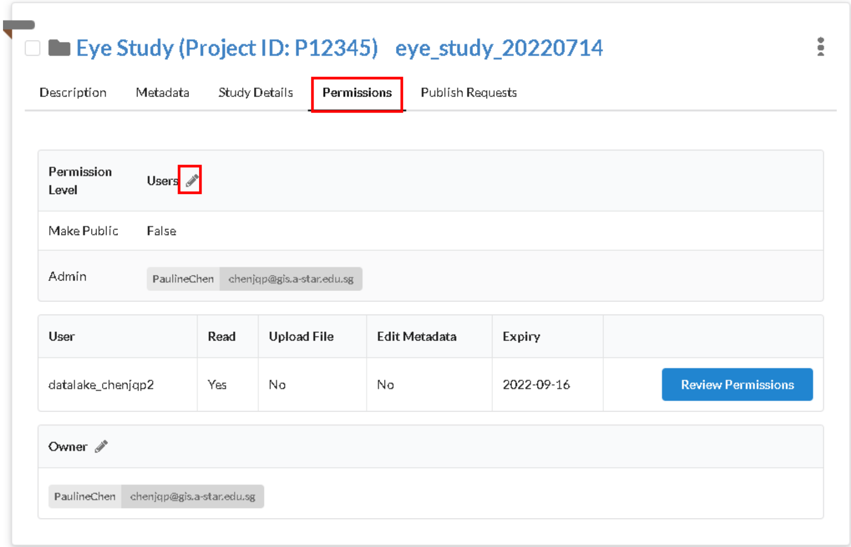


3. Add or delete users into the **Admin** permission level list, then click **Submit**.

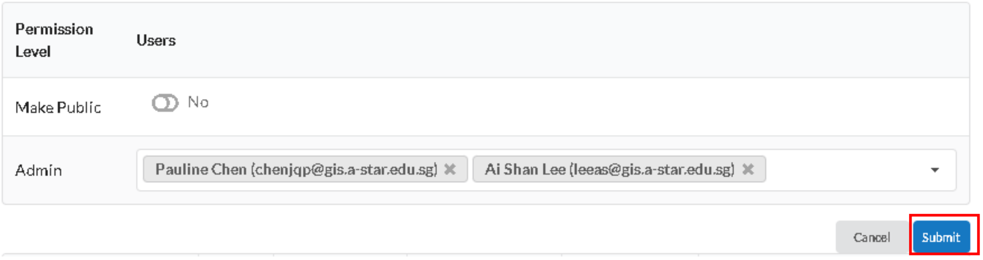

#### Modify existing permissions for a user on a data set

1. Go to the **Studies** tab

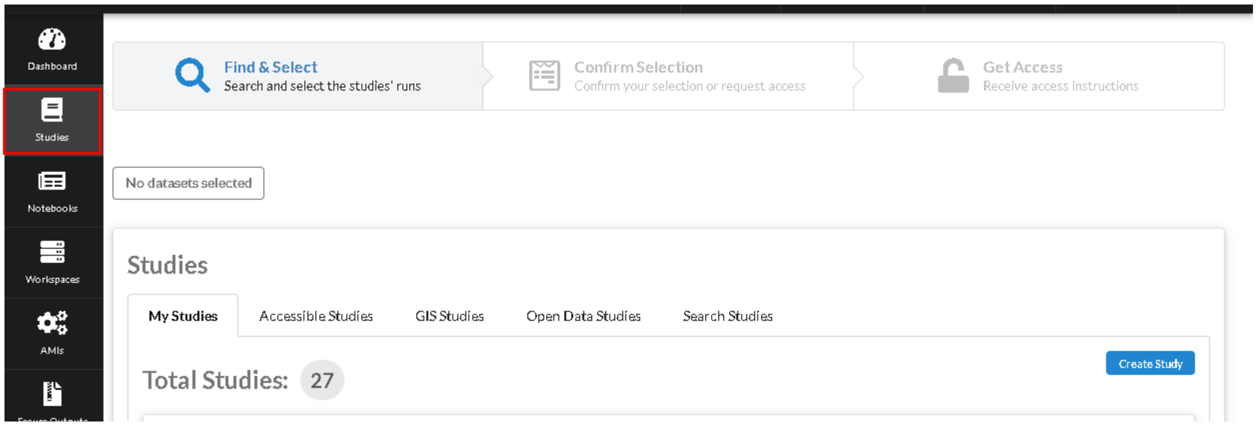


2. Under **My Studies** tab, locate the study card. Click on **Permissions** sub-tab, then click on **Review Permissions** button of the user whose permission you want to edit.

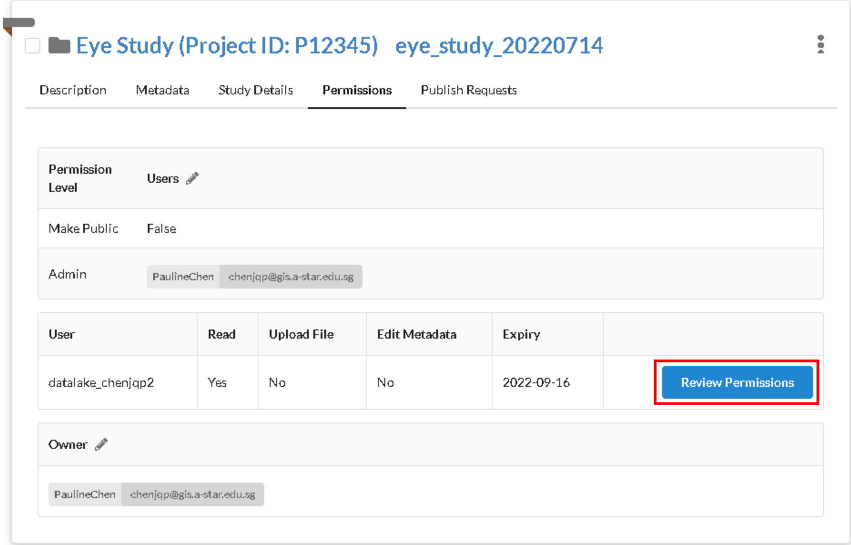


3. Edit the user's permission and click on **Update Permissions**. If you wish to revoke the user's permission completely, click on **Revoke**.

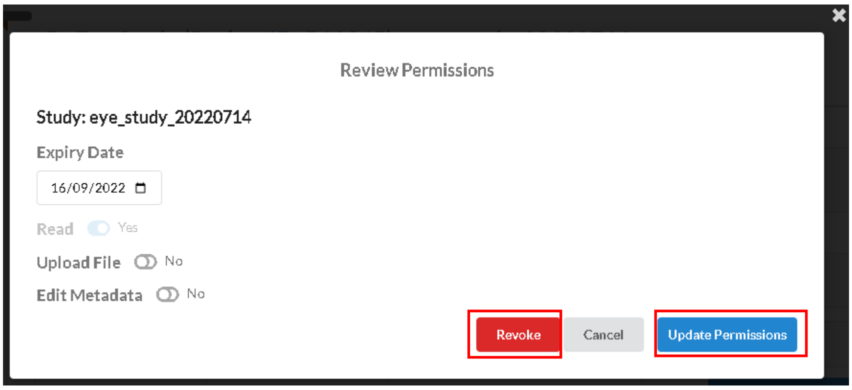

#### Revoke permissions for a user

4. Go to the **Studies** tab

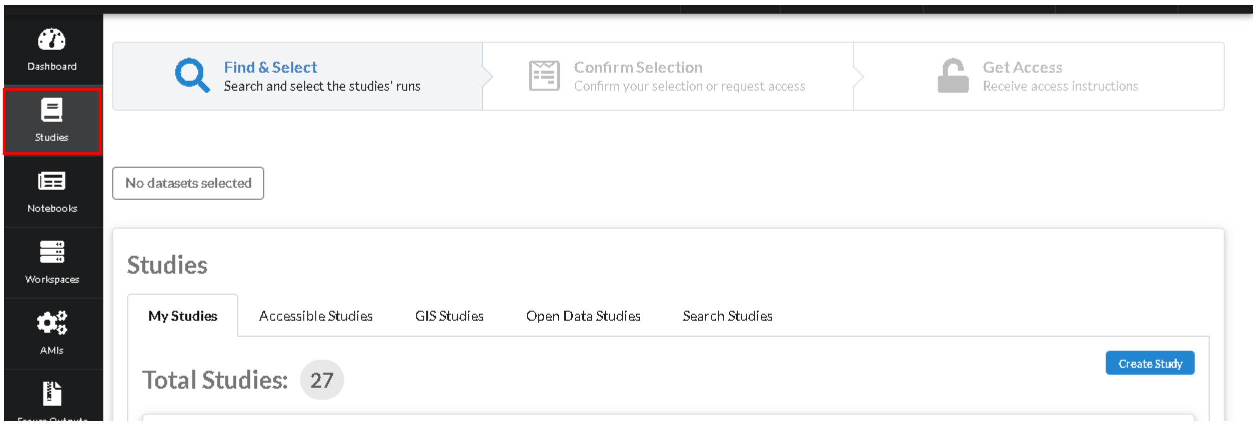


5. Under **My Studies** tab, locate the study card. Click on **Permissions** sub-tab, then click on **Review Permissions** button of the user whose permission you want to edit.

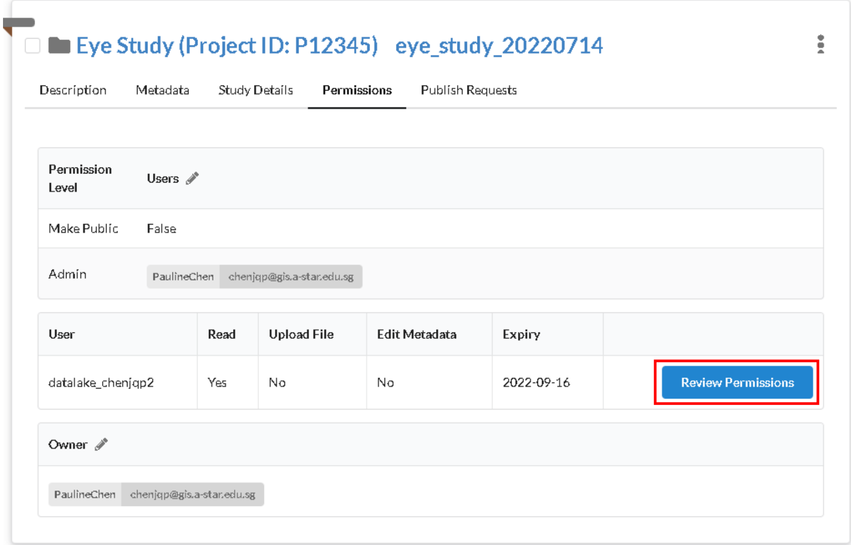


6. Edit the user's permission and click on **Update Permissions**. If you wish to revoke the user's permission completely, click on **Revoke**.

1. 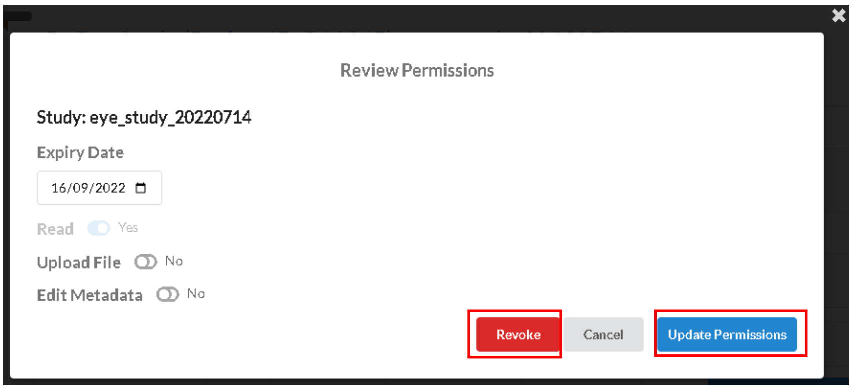

### Study Workspace

There are 2 types of workspaces:

- Private workspace: Used for private studies, public studies and API studies.
- Secure workspace: Used for secure private studies.

IMPORTANT: You should NEVER share workspaces with others as this may give access to studies that they may otherwise not have.

#### Create Workspace

1. Go to the **Studies** tab

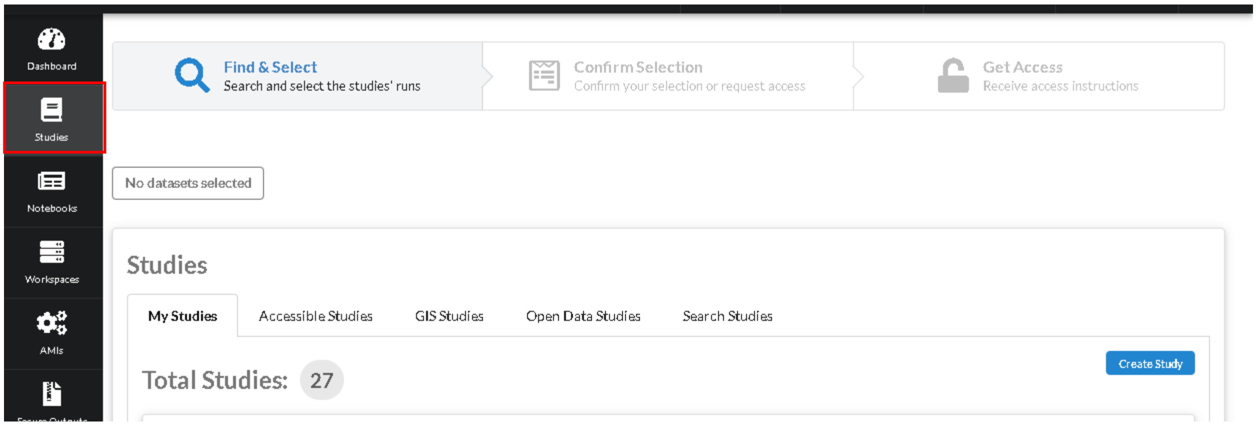


2. Select the studies you want to create a workspace with, then click **Next**.

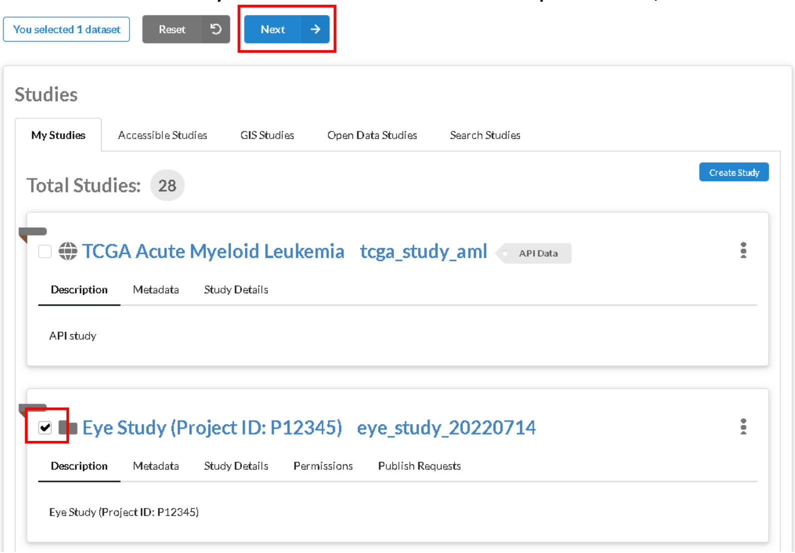

3. If you do not have sufficient permissions to all the selected studies, you will see the following:

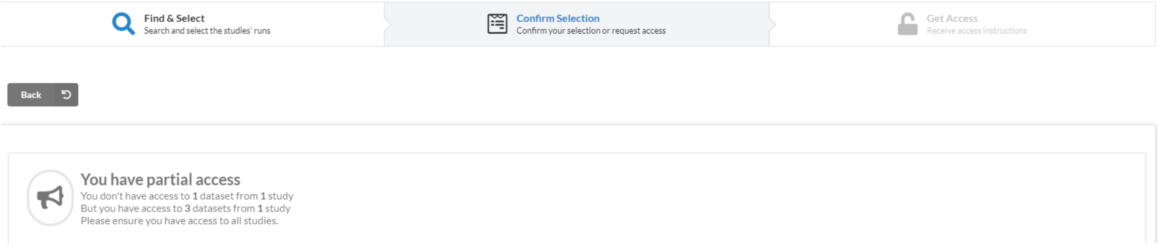


Please request for access for the affected studies if you see the above.

If you have all the required permissions, click **Set up a compute resource for me**.

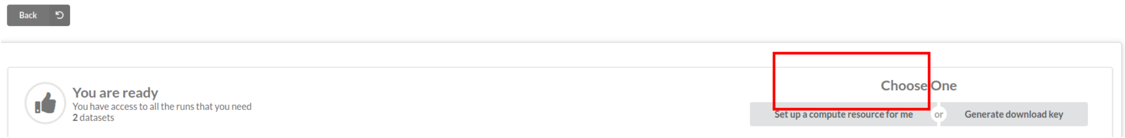


4. Select the resource type for your workspace based on your requirements and click **Configure**.

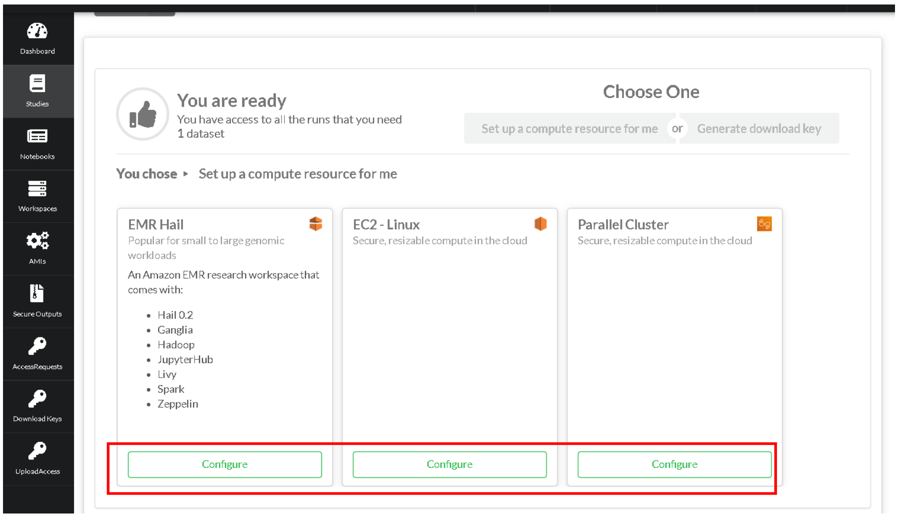

5. Fill up the workspace configuration form:
    a. Name: Workspace name, used to identify your workspaces
    b. Whitelisted CIDR: An IP CIDR value to whitelist access to your workspace. For security, it is highly recommended to set this value to your known IP range/address
    c. Project ID: Project to charge to (only 1 value available at the moment)
    d. Workspace size: Choose from the different sizes (Small, Medium, Large) or select your own (Custom – not available for EMR)
    e. Root Volume Size: The size (in GB) of the root volume.
    f. Workspace output study: Create a new output study or use an existing one that is not in use. (IMPORTANT: When creating an output study, you are required to select an **Output Storage Size**. Once created, this value cannot be changed.)

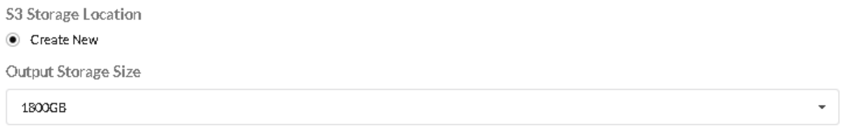


    g. (FOR PARALLEL CLUSTER) Min Counts: Minimum number of compute nodes
    h. (FOR PARALLEL CLUSTER) Max Counts: Maximum number of compute nodes
    i. Description: Workspace description

Once ready, click **Create Research Workspace**.

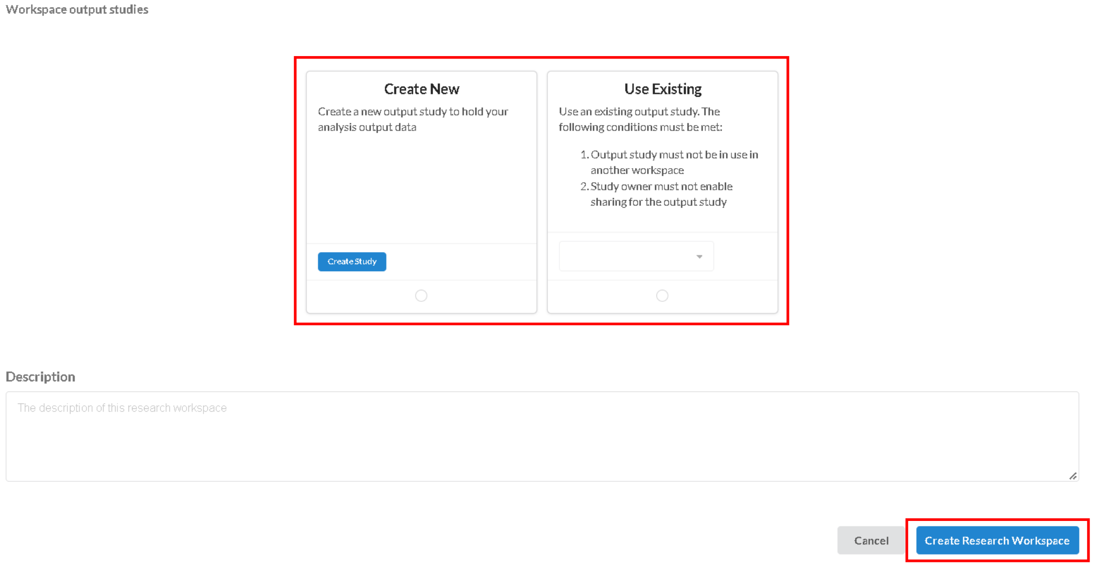


6. Once created, you will be redirected to the workspace page, where you will see your list of workspaces. While your workspace is being created, you will see the status as ‘Starting’. Please wait for the status to be listed as ‘Ready’.

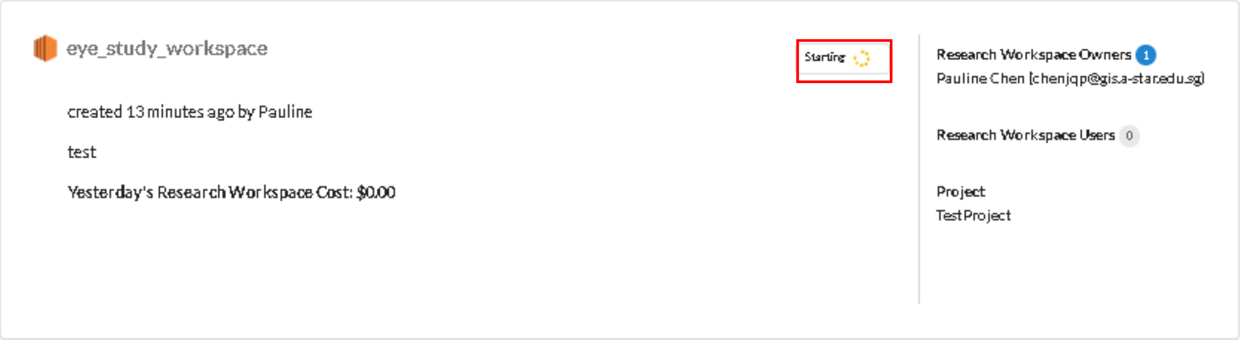
eye_study_workspace
Starting
created 13 minutes ago by Pauline
test
Yesterday's Research Workspace Cost: $0.00
Research Workspace Owners 1
Pauline Chen
Research Workspace Users 0
Project
Test Project

#### Create secure workspace

1. Go to the **Studies** tab

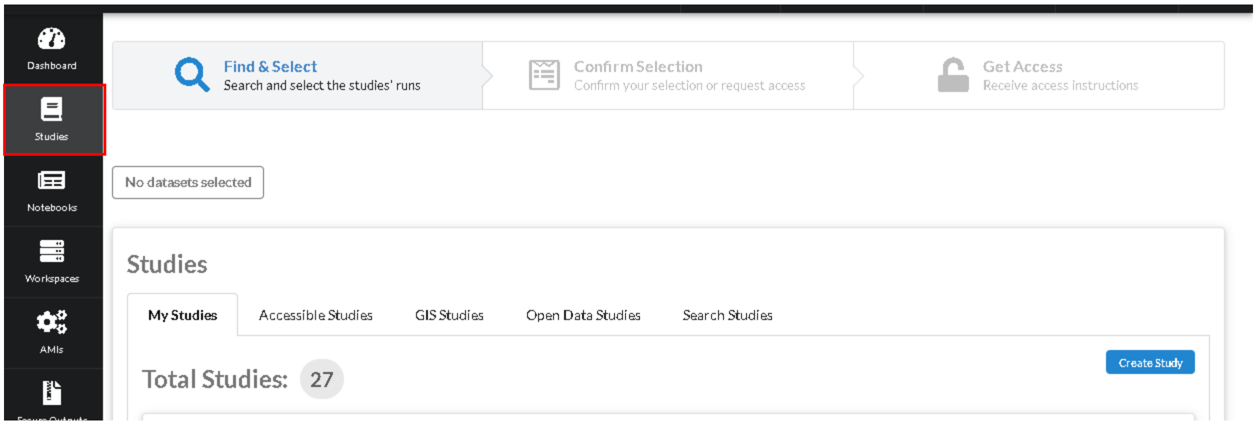


2. Select the **secure** studies (and any additional non-secure studies **excluding API study**) you want to create a workspace with, then click **Next**.

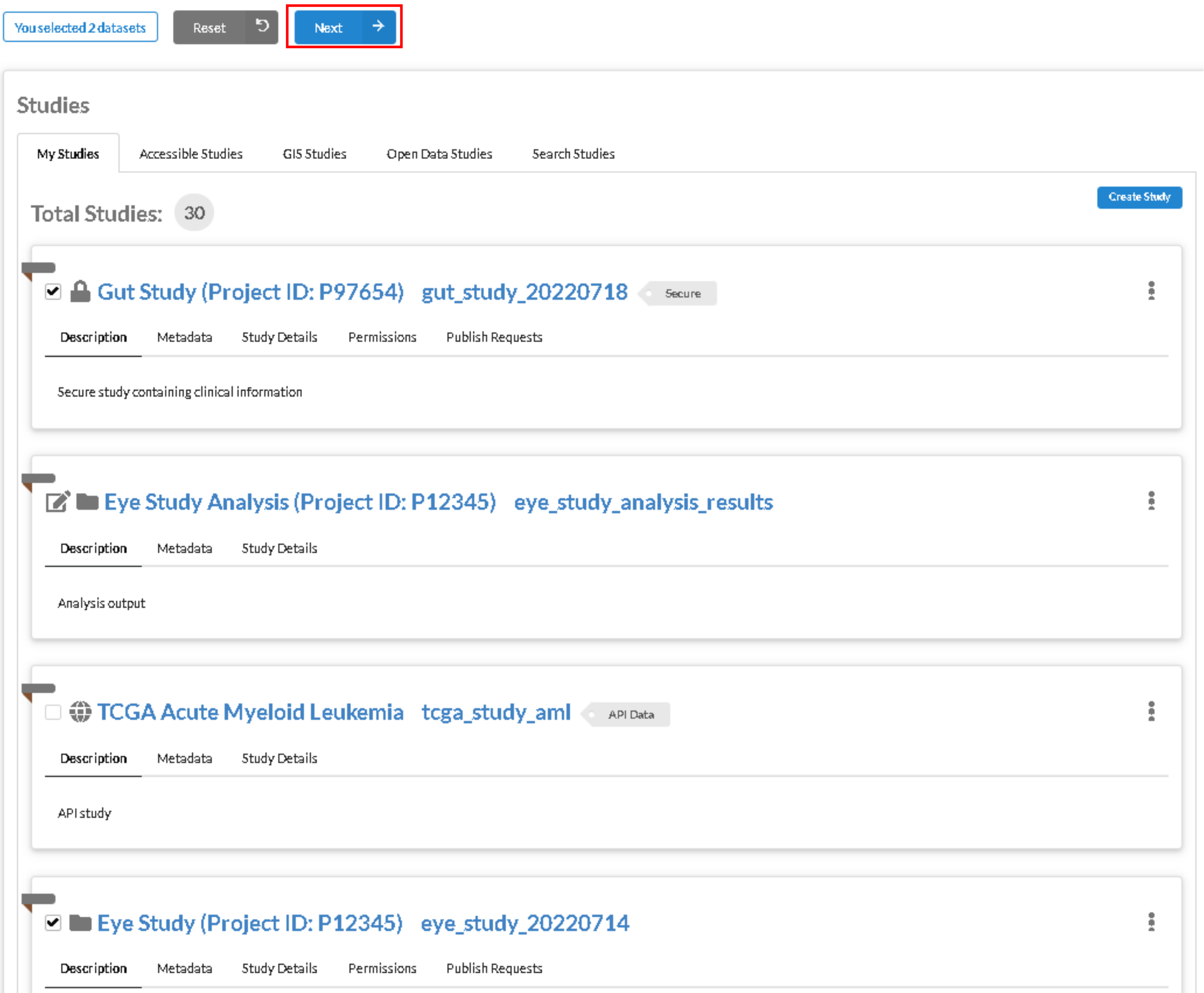


Selecting secure studies and API study will result in a conflict. Click **Back** to change your selection if you encounter the conflict.

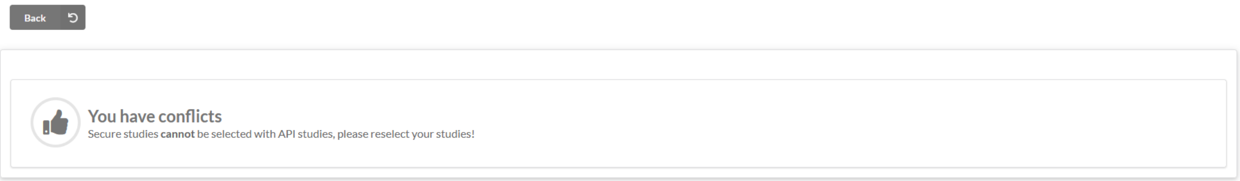


3. Once you have all the required study permissions, click on **Set up a SECURE compute resource for me** and select a workspace type available.

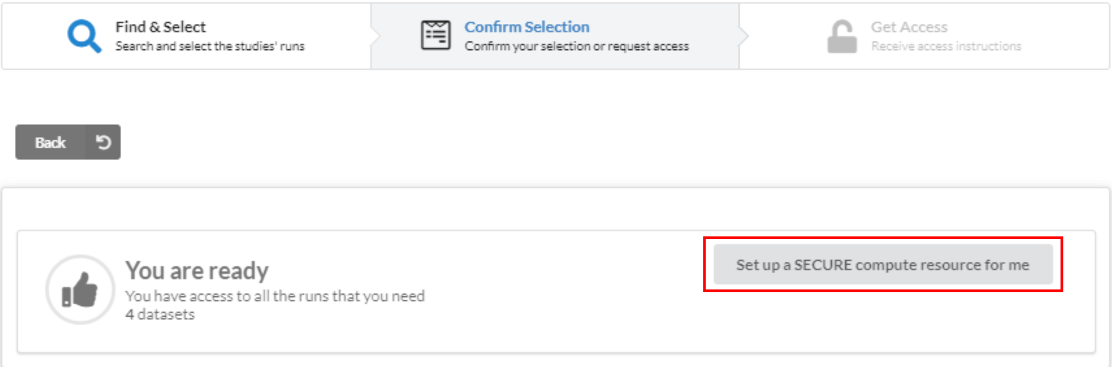


4. Select the resource type for your workspace based on your requirements and click **Configure**.

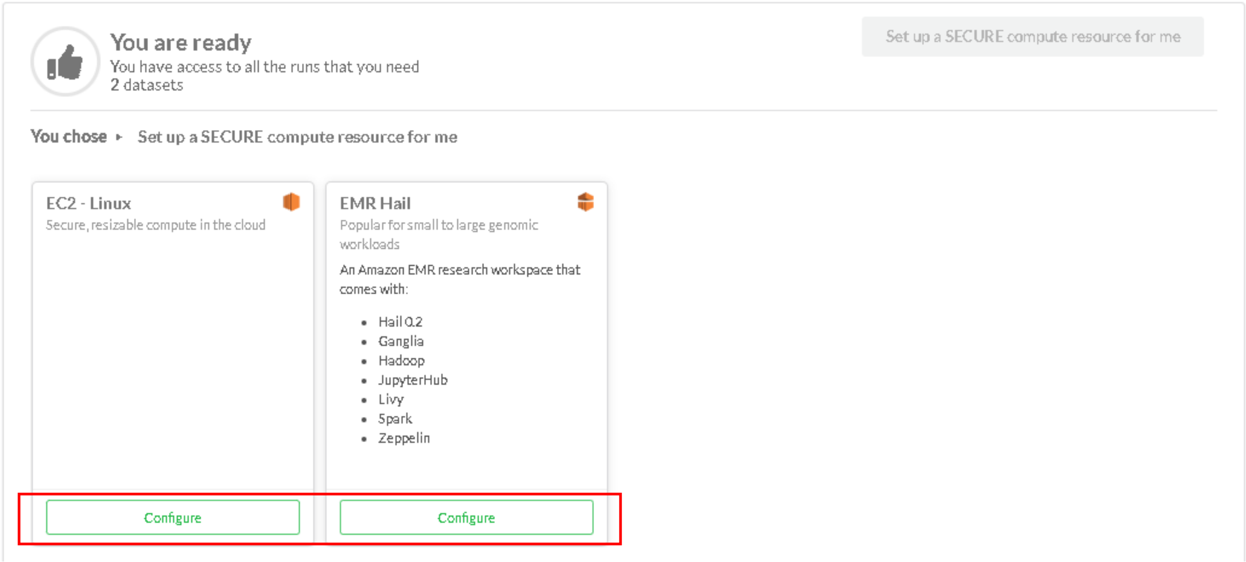


5. Fill up the workspace configuration form:
   a. Name: Workspace name, used to identify your workspaces
   b. Whitelisted CIDR: An IP CIDR value to whitelist access to your workspace. For security, it is highly recommended to set this value to your known IP range/address
   c. Project ID: Project to charge to (only 1 value available at the moment)
   d. Workspace size: Choose from the different sizes (Small, Medium, Large) or select your own (Custom – not available for EMR)
   e. Root Volume Size: The size (in GB) of the root volume.
   f. Workspace secure output: Create a new secure output or use an existing one that is not in use. (IMPORTANT: When creating a secure output, you are required to select an **Output Storage Size**. Once created, this value cannot be changed.) For more details on secure outputs, see Secure Output.

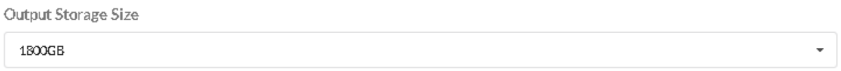


g. Description: Workspace description

Once ready, click **Create Research Workspace**.

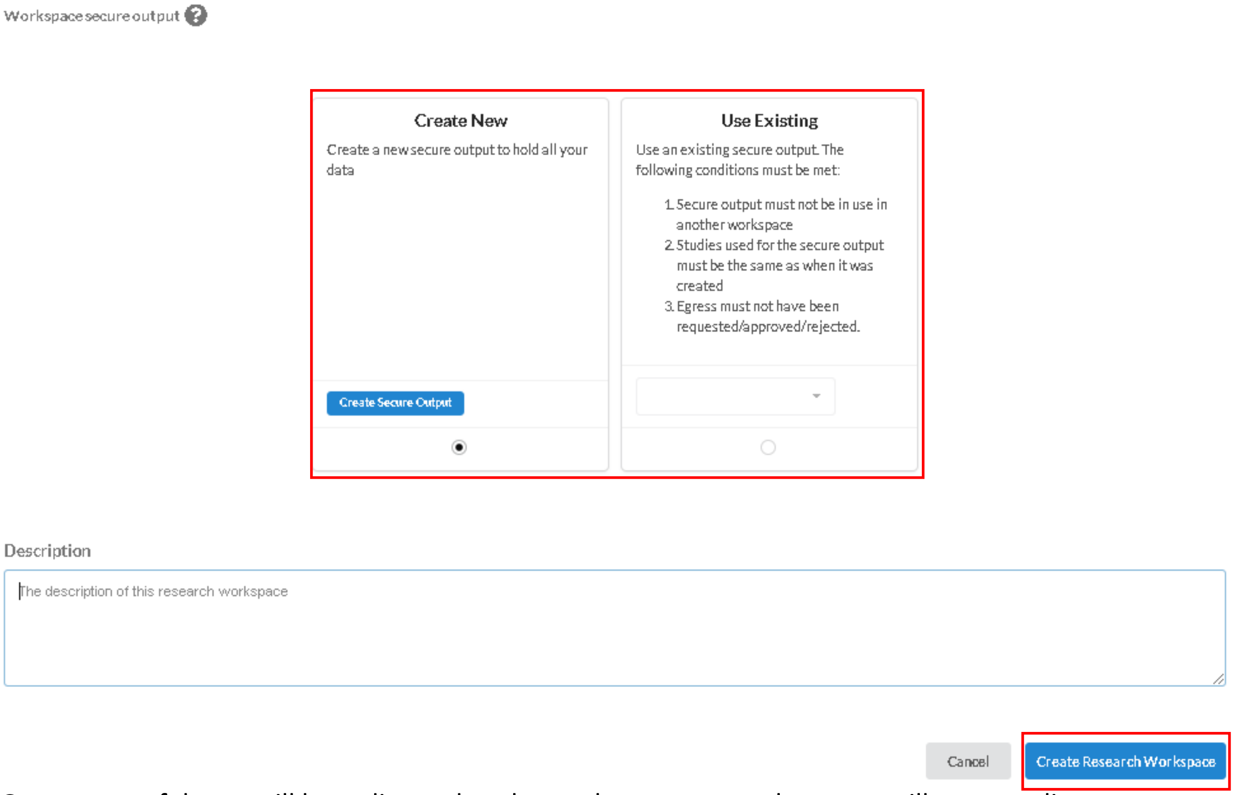


6. Once successful, you will be redirected to the workspace page, where you will see your list of workspaces. While your workspace is being created, you will see the status as 'Starting'. Please wait for the status to be listed as 'Ready'.

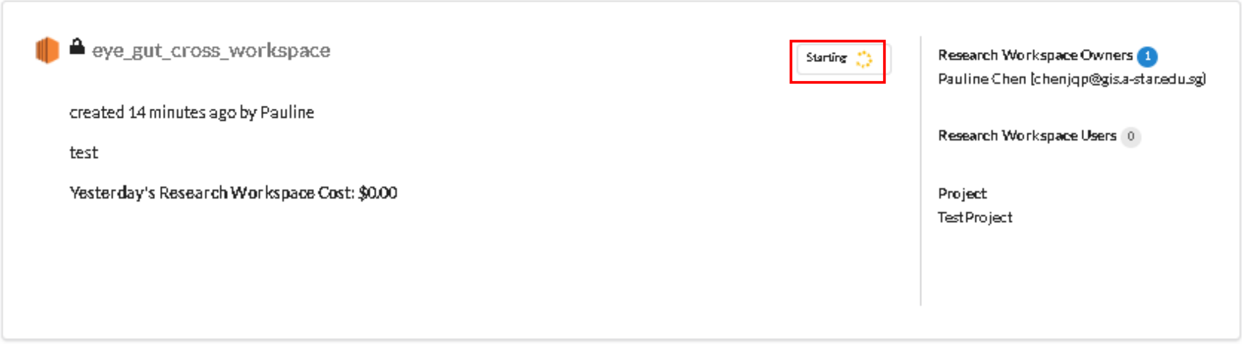


#### View Workspace Details

1. Go to the **Workspaces** tab

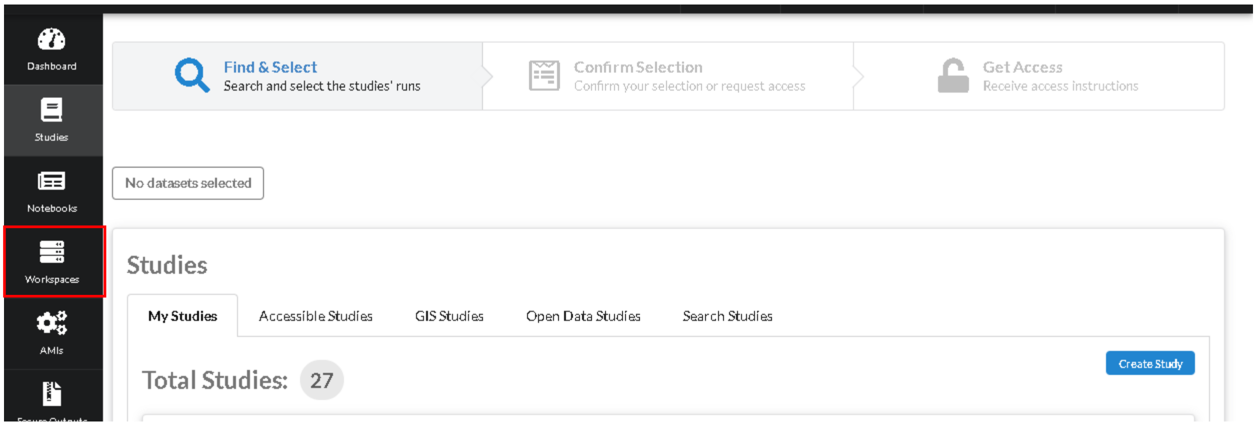


2. Search for your workspace card and ensure that the status of the workspace is **Ready**.

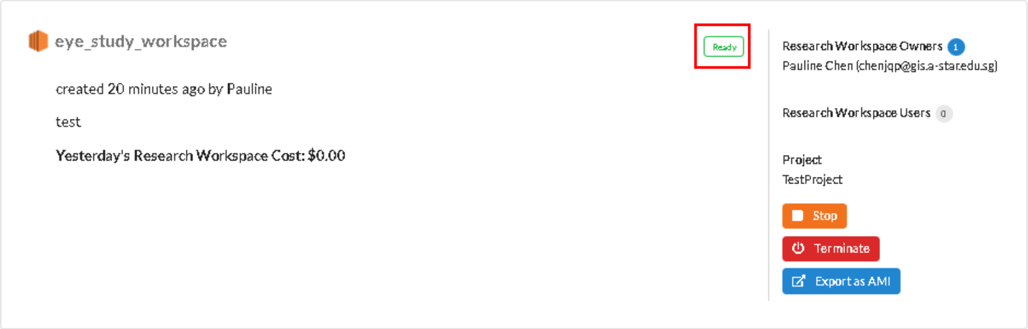


3. Click on the card to view details.

#### Access Instructions

Click on **Access Instructions** tab.

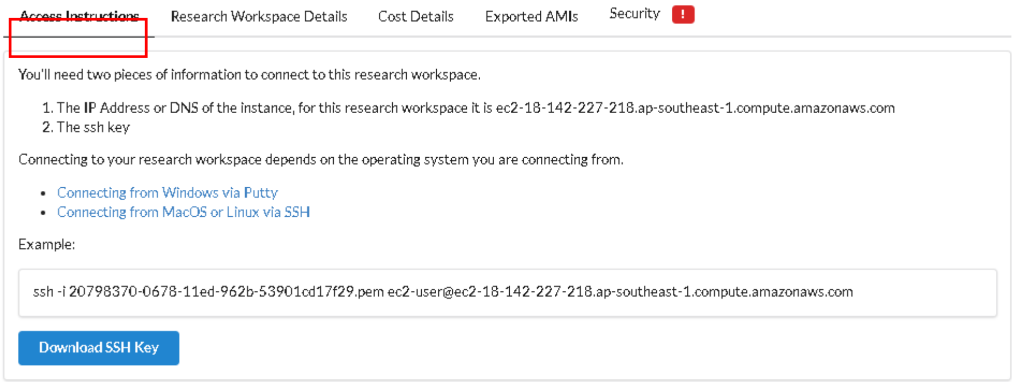

#### Workspace studies

Click on **Research Workspace Details** tab.

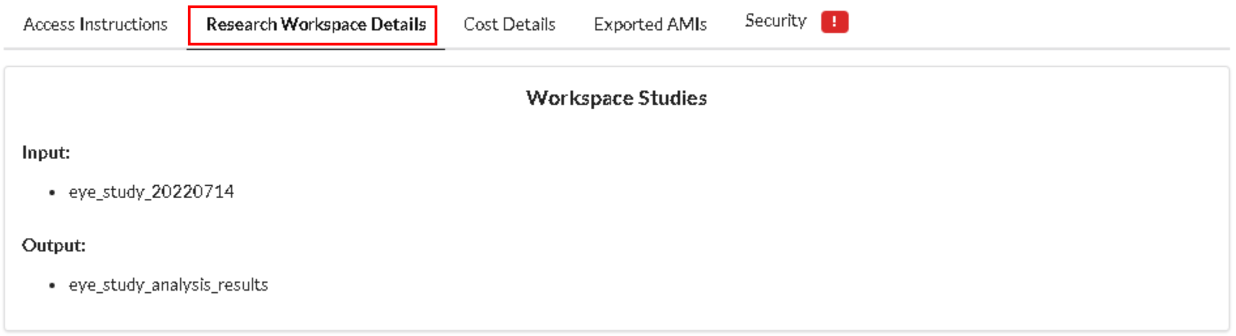


#### Costs for workspace

Click on **Cost Details** tab.

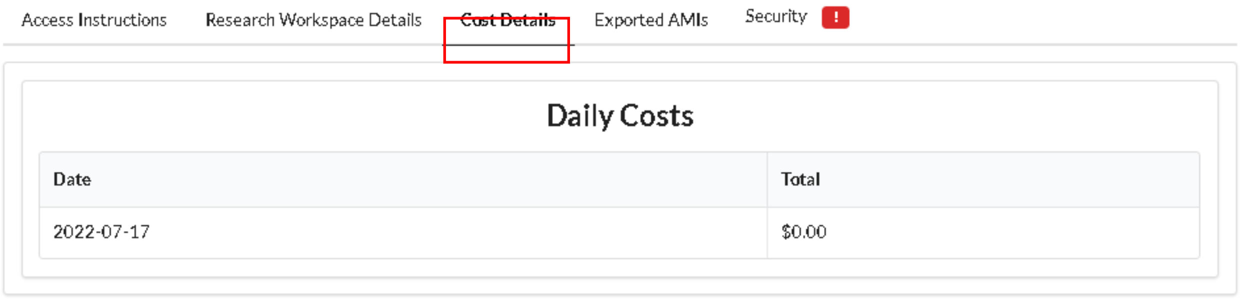


#### AMIs created from workspace

Click on **Exported AMIs** tab.

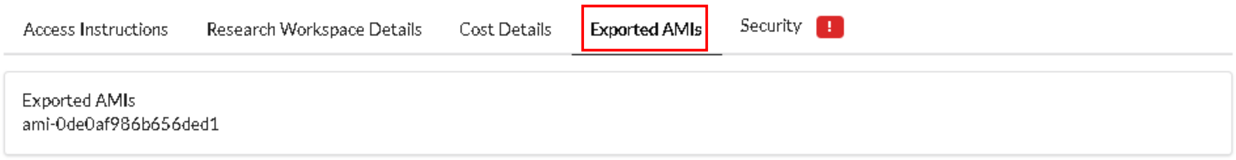

#### Security settings

Click on **Security** tab.

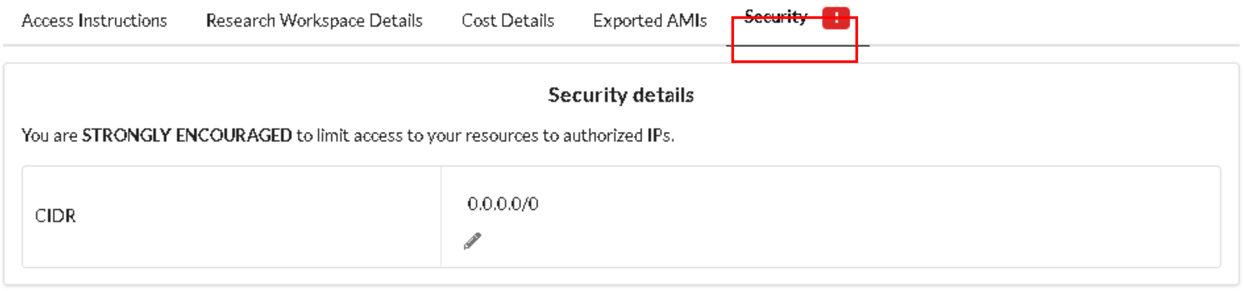


To update your security settings, click on the pencil button, update your CIDR in the text field then click **Submit**.

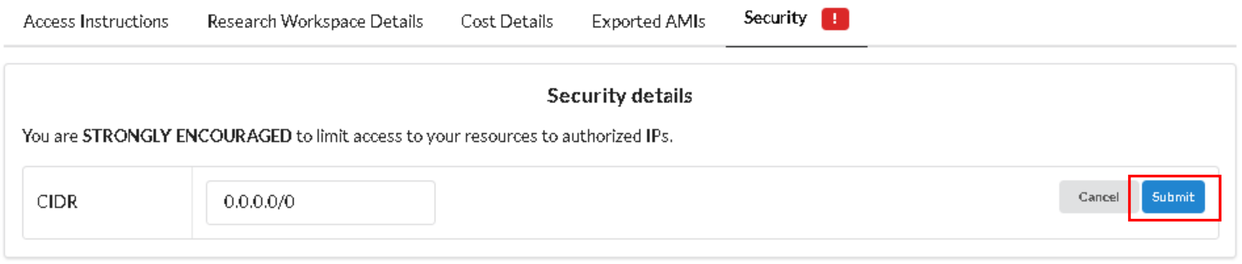


#### Access Workspace

See Access Instructions

1. For EC2/parallel cluster workspaces:

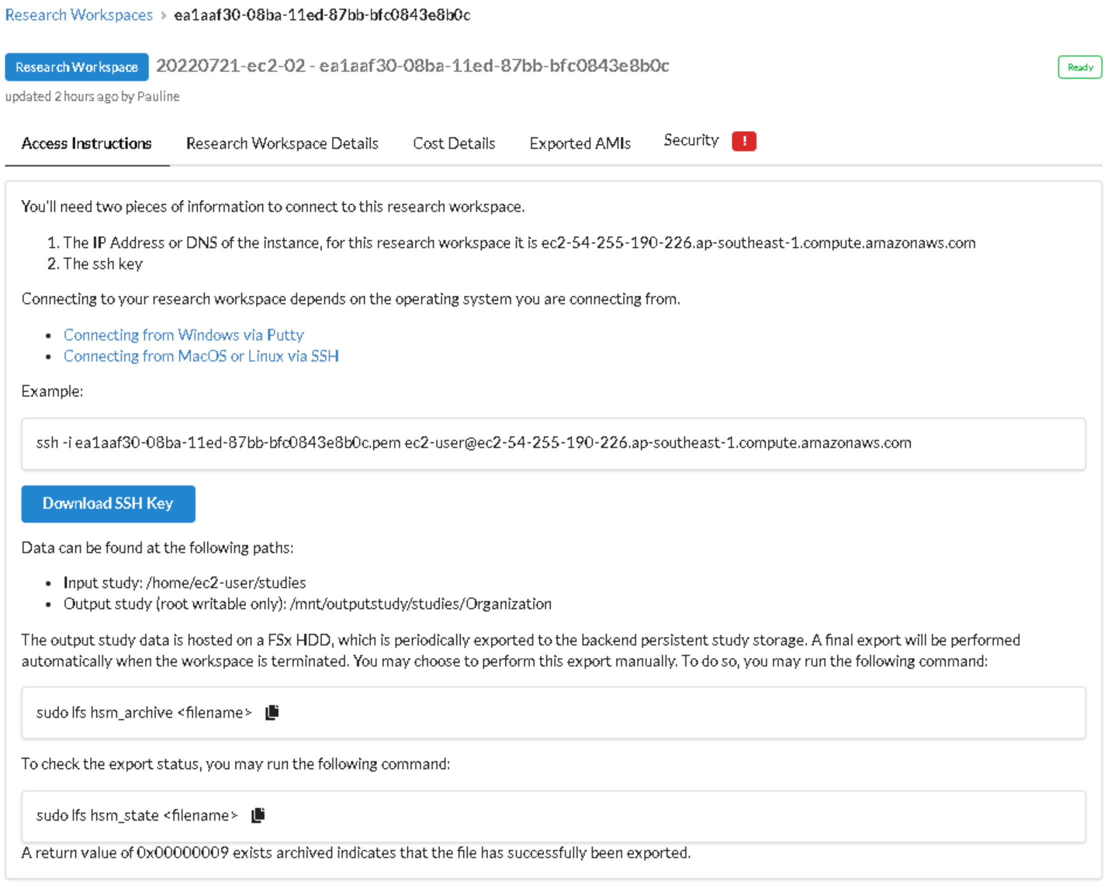


a. Click on **Download SSH Key**
b. Open a SSH client on your desktop (PuTTY, command prompt, powershell etc)

c. Navigate to the location of the key and run the command on your card

d. Once you've accessed your workspace, data can be found in the following paths:
   i. Study data (READ ONLY): ~/studies/
   ii. API data (READ ONLY) : ~/api/
   iii. Output study (READ-WRITE with root): /mnt/outputstudy/studies/Organization/

e. The output study data is hosted on a FSx HDD, which is periodically exported to the backend persistent study storage. A final export will be performed automatically when the workspace is terminated. You may choose to perform this export manually. To do so, you may run the following command:

```
sudo lfs hsm_archive <filename>
```

To check the export status, you may run the following command:

```
sudo lfs hsm_state <filename>
```

A return value of "0x00000009 exists archived" indicates that the file has successfully been exported.

2. For EMR workspaces:

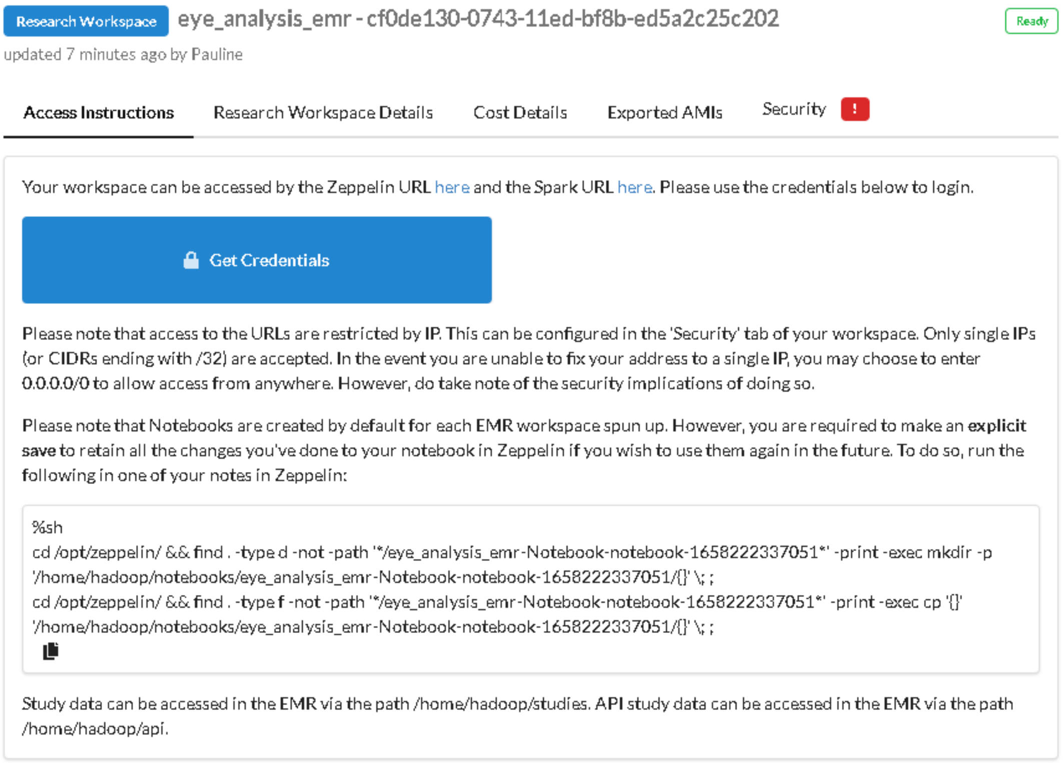


a. Click on the respective links to be directed to the Zeppelin workspace or the Spark page

b. Once you've accessed your workspace, data can be found in the following paths:
   i. Study data (including output study): /home/hadoop/studies/
   ii. API study data: /home/hadoop/api/
   iii. Notebook data: /home/hadoop/notebooks/

#### Access Secure Workspace

Secure workspaces can only be accessed through a bastion remote Windows host with restricted functionality.

1. If you're on a Windows machine, open 'Remote Desktop Connection'

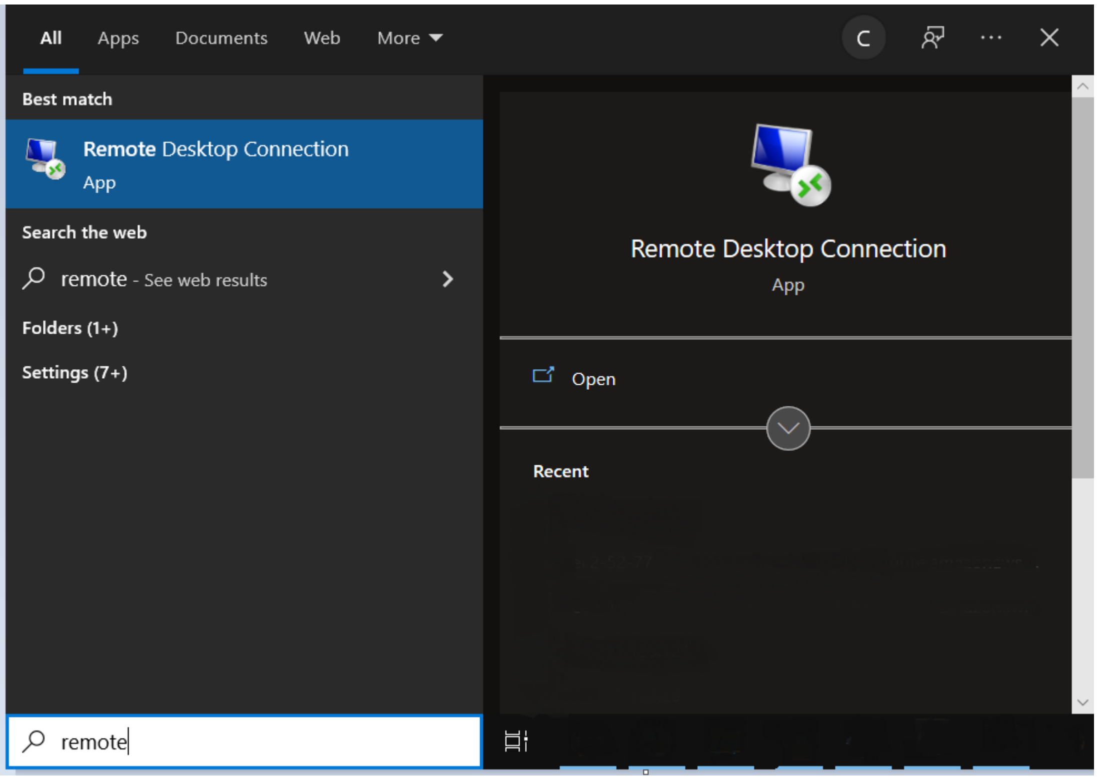


2. Enter the details specified in the **Access Details** tab of the workspace when prompted.

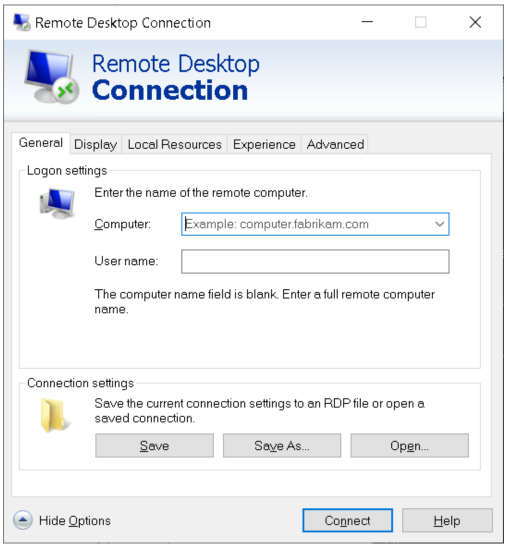


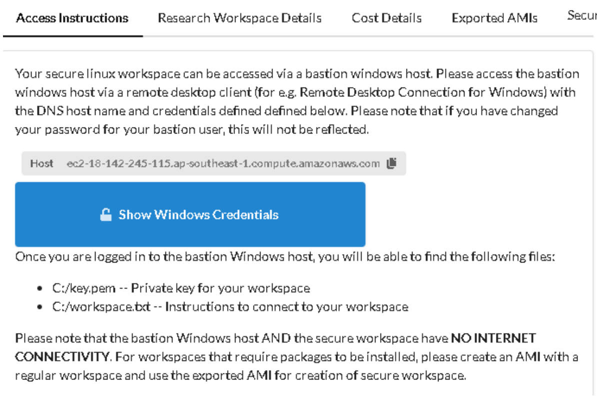


Computer: <Host>

Username: <Username>

Password (prompt): <Password>

3. For EC2 workspaces:
    a. In the bastion host, you will find the instructions to access the workspace node in C:/workspace.txt, with your key in C:/key.pem

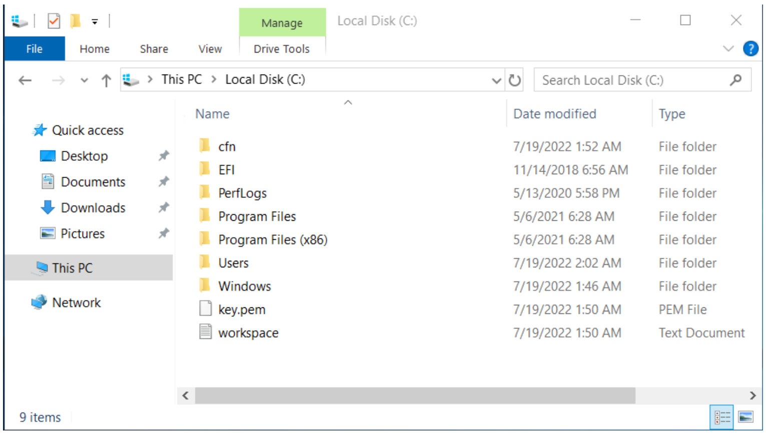


    b. Copy the C:/key.pem onto the bastion's desktop.
    c. Open command prompt or powershell on the bastion, navigate to the desktop and run the connect command as specified in the workspace.txt

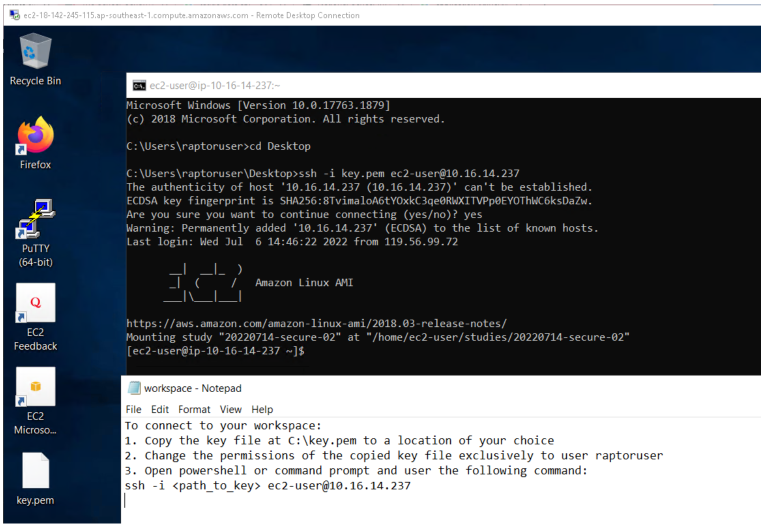


d. Once you've accessed your workspace, data can be found in the following paths:
   i. Study data (READ-ONLY): ~/studies/
   ii. Secure output (READ-WRITE with root): /mnt/outputstudy/secure-outputs

e. The secure output data is hosted on a FSx HDD, which is periodically exported to the backend persistent study storage. A final export will be performed automatically when the workspace is terminated. You may choose to perform this export manually. To do so, you may run the following command:
`sudo lfs hsm_archive <filename>`
To check the export status, you may run the following command:
`sudo lfs hsm_state <filename>`
A return value of "0x00000009 exists archived" indicates that the file has successfully been exported.

4. For EMR workspaces:
   a. In the bastion host, you will find the instructions to access the workspace in C:/workspace.txt

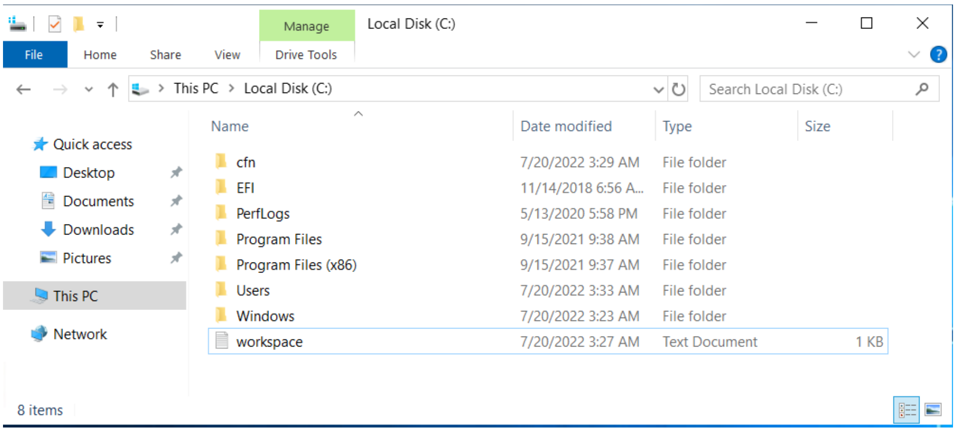


   b. Navigate to the URLs specified in the workspace.txt for Zeppelin and Spark.

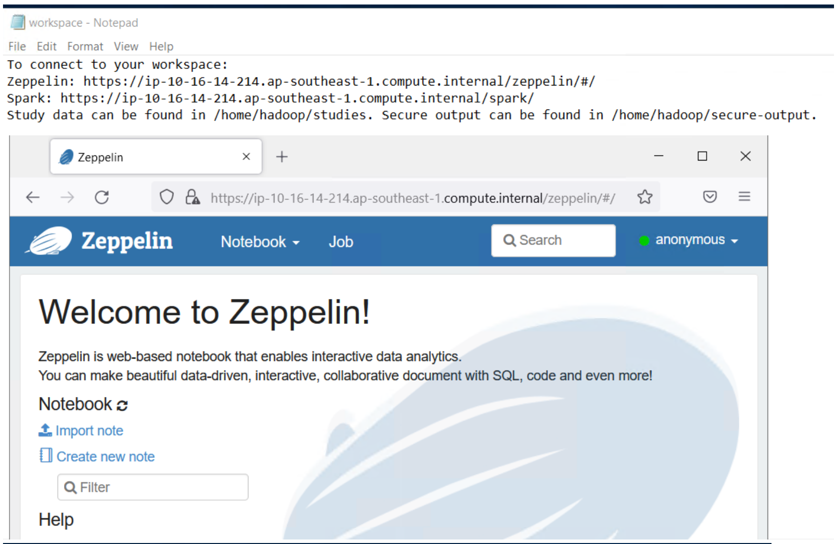


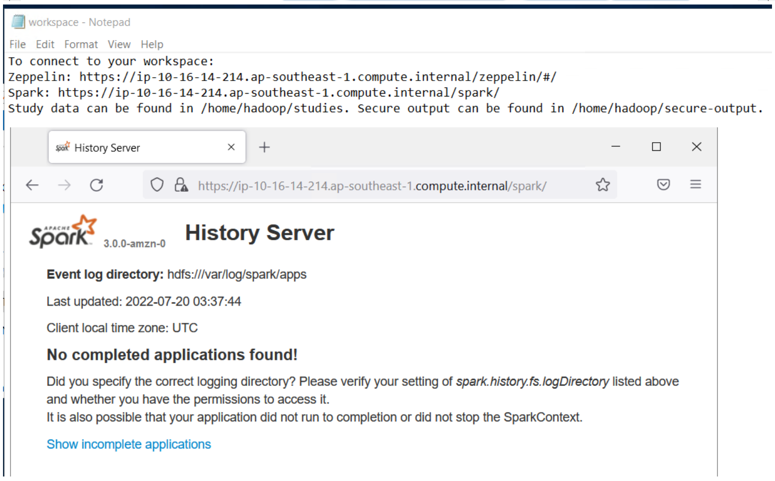


c. Once you've accessed your Zeppelin, data can be found in the following paths:
   i. Study data (READ-ONLY): /home/hadoop/studies/
   ii. Secure output (READ-WRITE): /home/hadoop/secure-output

#### Stop Workspace

Please note that not all workspaces support the start/stop function. You must be the owner of the workspace to perform this action.

1. Go to the **Workspaces** tab

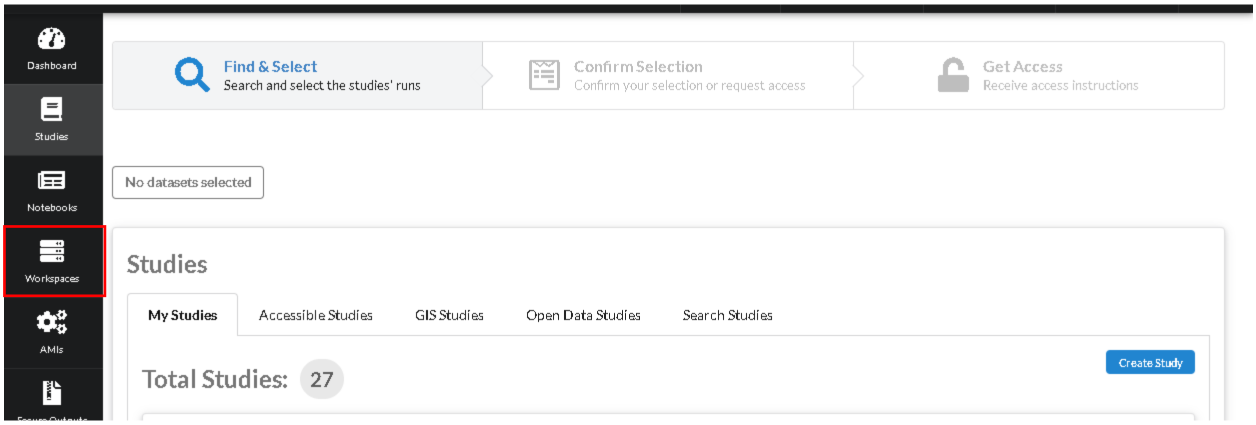


2. Click on the **Stop** button of the workspace.

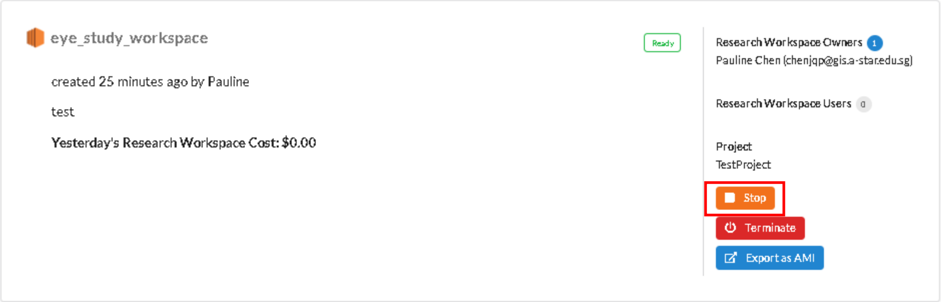


3. The workspace will change to a **Stopping** status

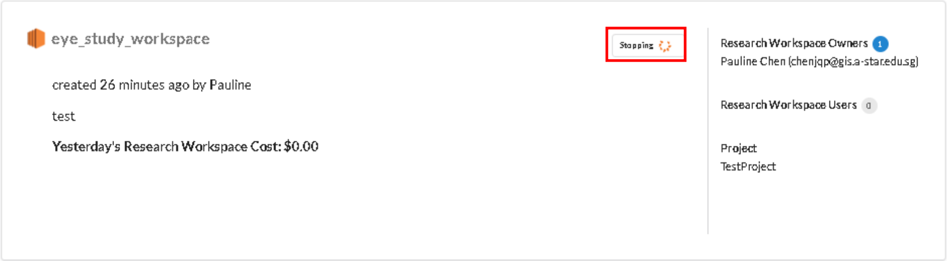


4. Once successful, the workspace will change to a **Stopped** status.

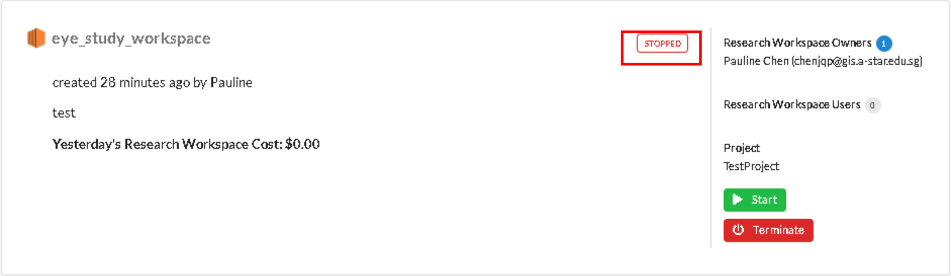

#### Start Workspace

Please note that not all workspaces support the start/stop function. You must be the owner of the workspace to perform this action.

1. Go to the **Workspaces** tab

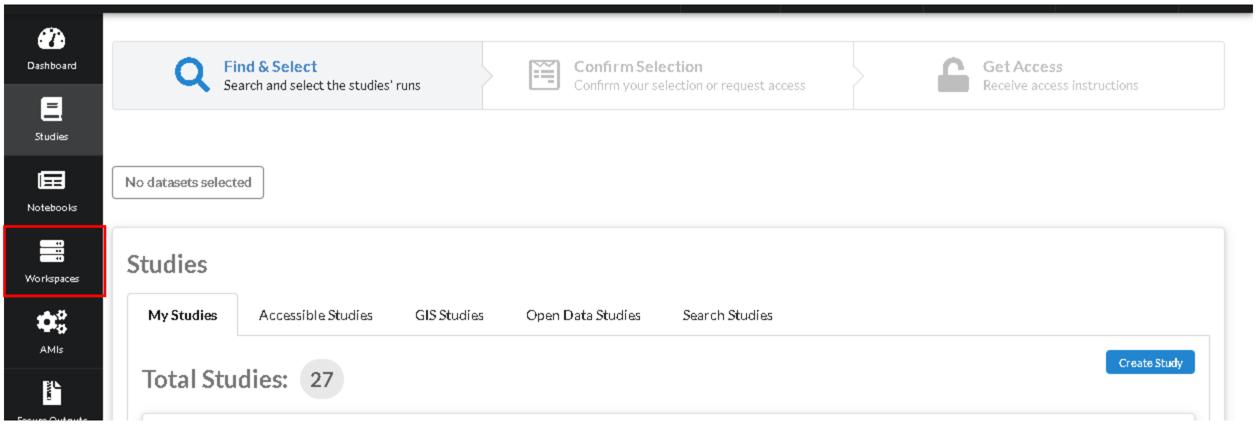


2. Click on the **Start** button of the workspace.

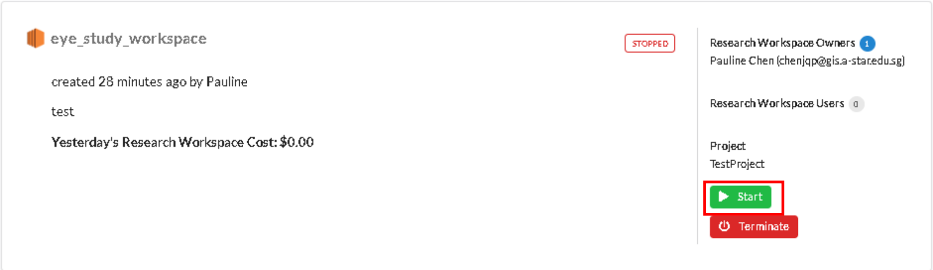


3. The workspace will change to a **Starting** status

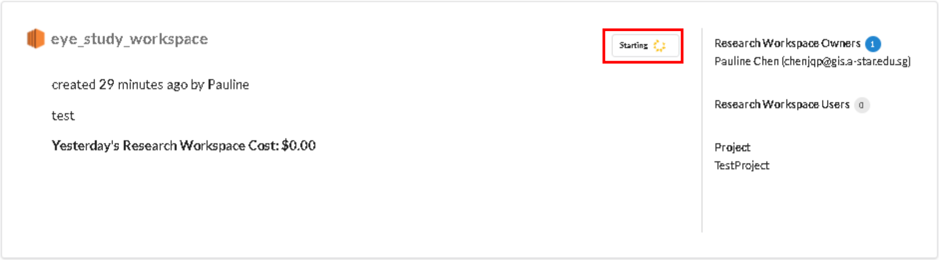


4. Once successful, the workspace will be in a **Ready** status.

#### Terminate Workspace

Please note that once a workspace is terminated, all unsaved data will be lost. You will be unable to restart the workspace. You must be the owner of the workspace to perform this action.

1. Go to the **Workspaces** tab

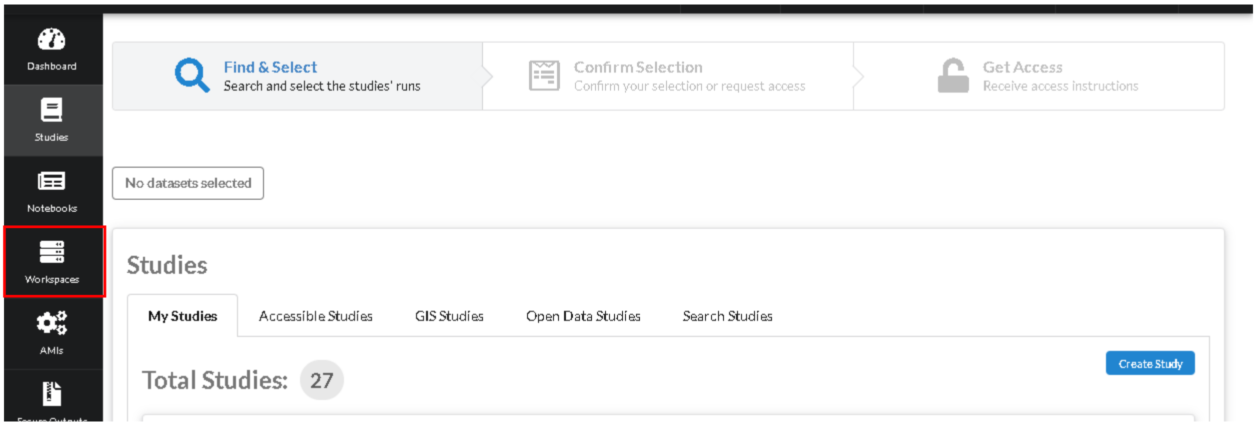


2. Click on the **Terminate** button for the workspace you want to terminate

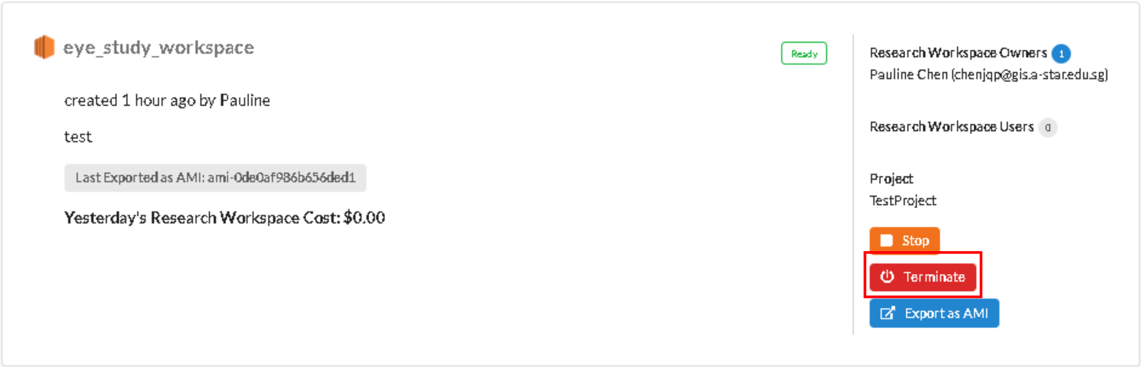


3. The workspace will change status to **Terminating**

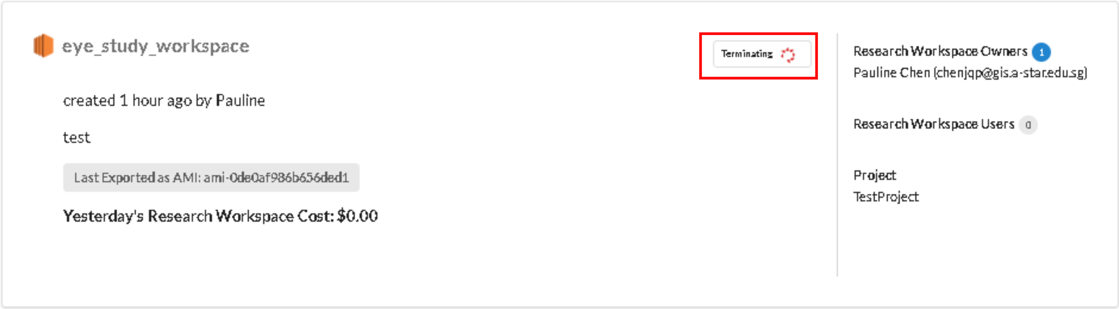

#### Create AMI

Please note that this action is only available for EC2-linux workspaces. To grant other users access to your AMI, please contact RAPTOR administrators.

1. Go to the **Workspaces** tab

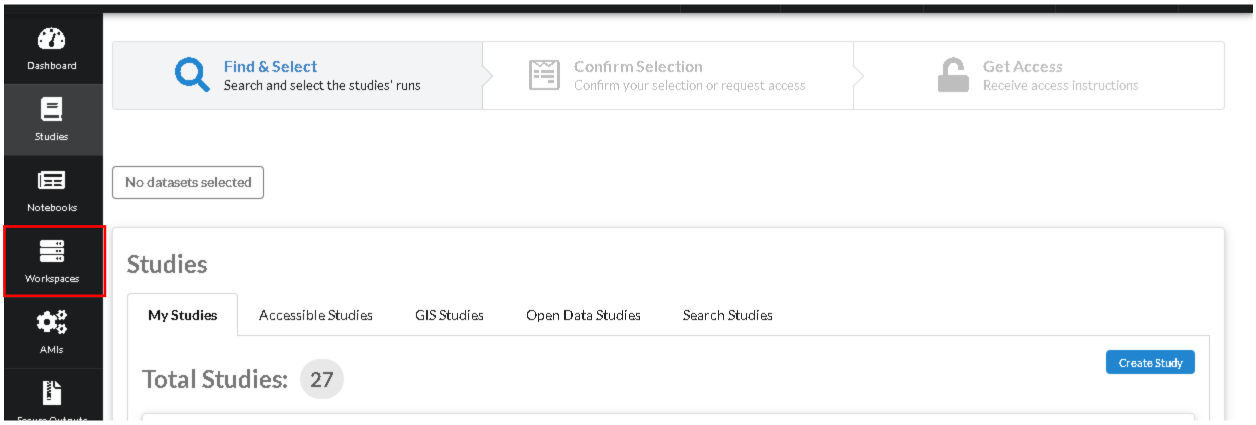


2. Click **Export as AMI** for the workspace.

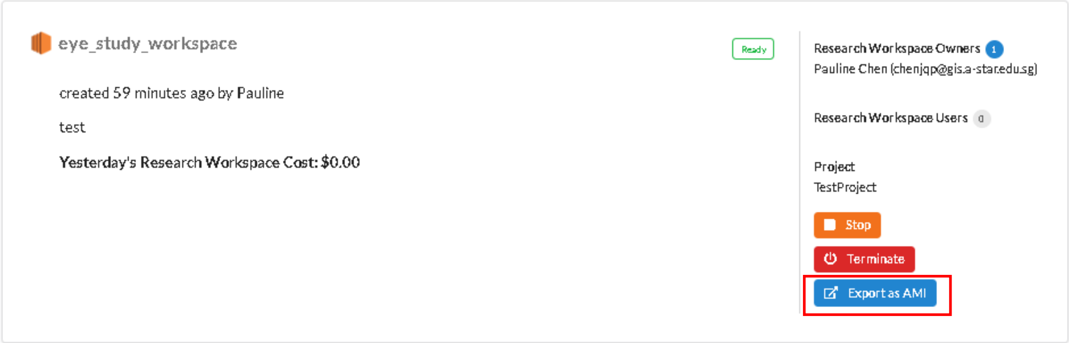


3. You will be redirected to your list of AMIs. Wait for the status of the AMI to turn to **Available**.

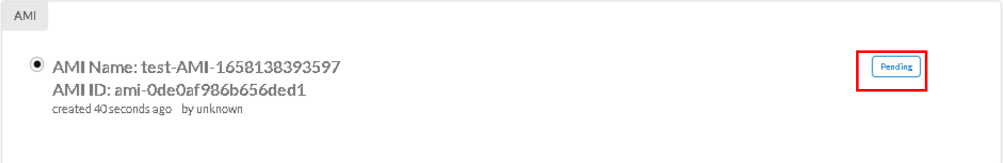


**Important note:** If you would like to import your own AMI, please contact RAPTOR administrators. Please ensure that the following **must be installed:**

- Jq
- Fuse
- Lustre client (https://docs.aws.amazon.com/fsx/latest/LustreGuide/install-lustre-client.html)

#### Create Workspace from AMI

Please note that the AMI must be in **Available** state to perform this action.

1. Go to the **AMIs** tab

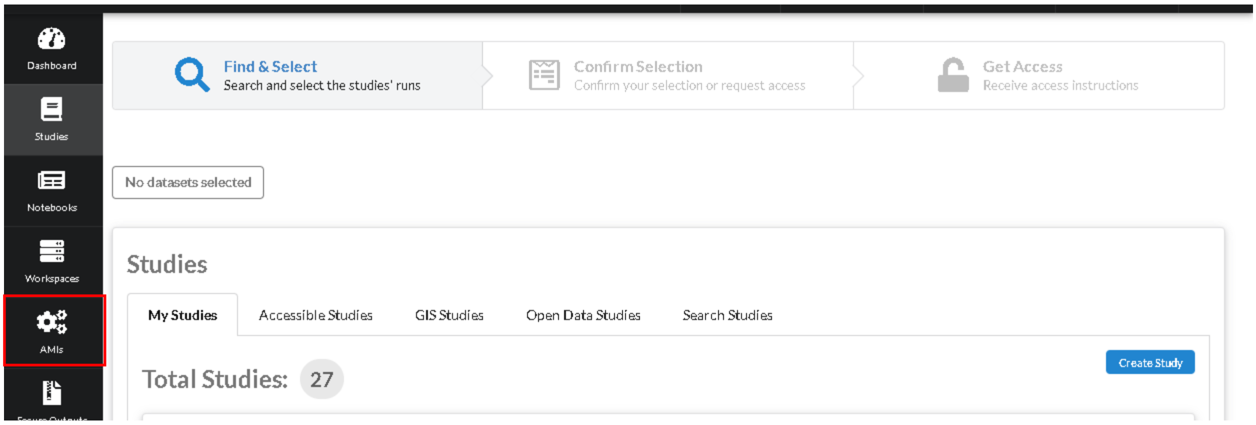


2. Select the AMI to use and click **Create Research Workspace**.

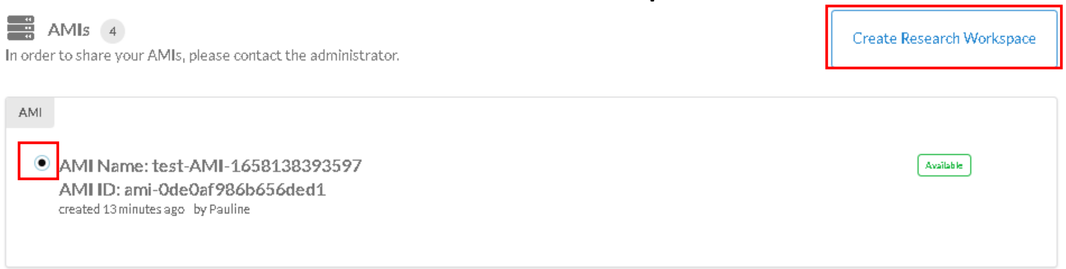


3. Continue from step 2 of Create Workspace

#### Secure Output

A secure output is used to hold any data derived from secure studies. For secure outputs that contain such data from secure studies not belonging to you, an explicit egress request needs to be submitted to the secure study owner via the **Freeze and Egress** button. The secure study owner will have to approve or reject your request after vetting data in your secure output. If you use secure studies that belong to you, you can immediately copy data out into a study of your choice after clicking **Freeze and Egress.**

##### Request Freeze and Egress

You can only request freeze and egress for secure outputs that are not attached to any active workspace. Secure outputs left idle for 30 days will have their data automatically deleted.

1. Go to the **Secure Outputs** tab.

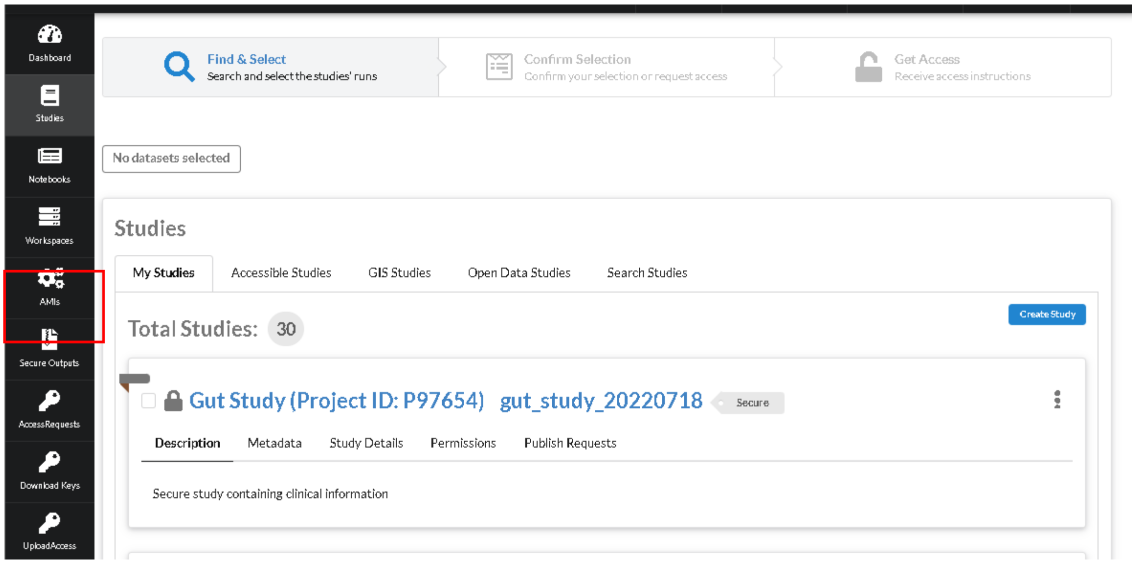


2. Under **My Secure Outputs**, locate the secure output card. Click on **Freeze and egress**.

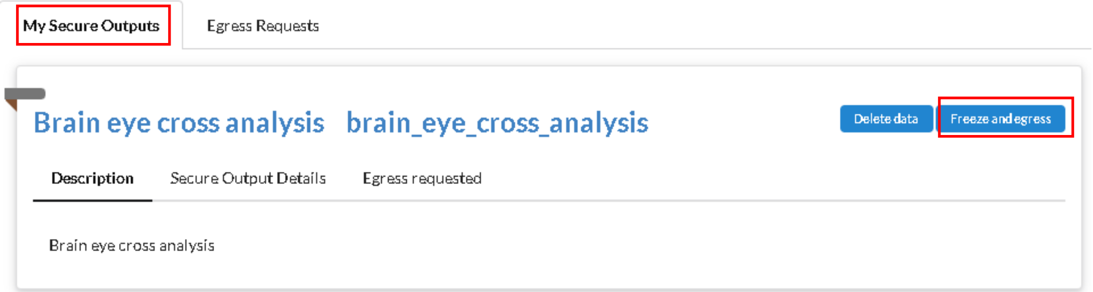


3. If the secure output has pending approvals from secure study owners (i.e. it contains data from secure studies that you are not an owner of), it will transit to a **Pending** status. You will now have to wait for the respective study owners to vet through and approve the data within the secure output.

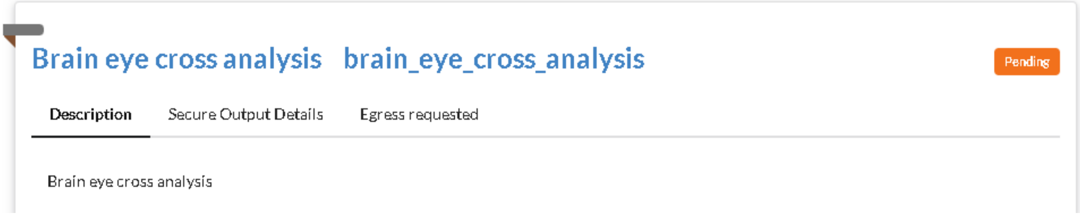

If the secure output contains data from secure studies that **<u>belong to you</u>**, you will immediately be able to copy the data out.

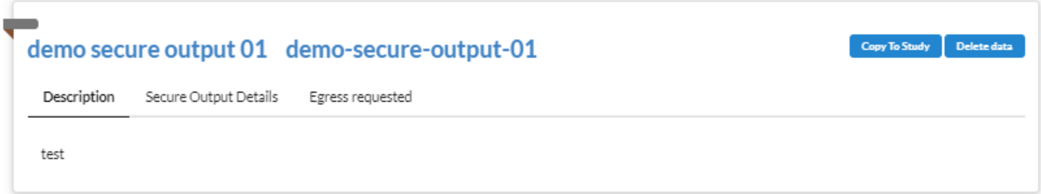

#### Vet Egress Requests

Upon receiving a request for egress from a secure output that may contain derived data from your secure study, you will need to vet the secure output for data that you may not want the requestor to take out (for e.g. sensitive clinical data).

1. Go to the **Secure Outputs** tab.

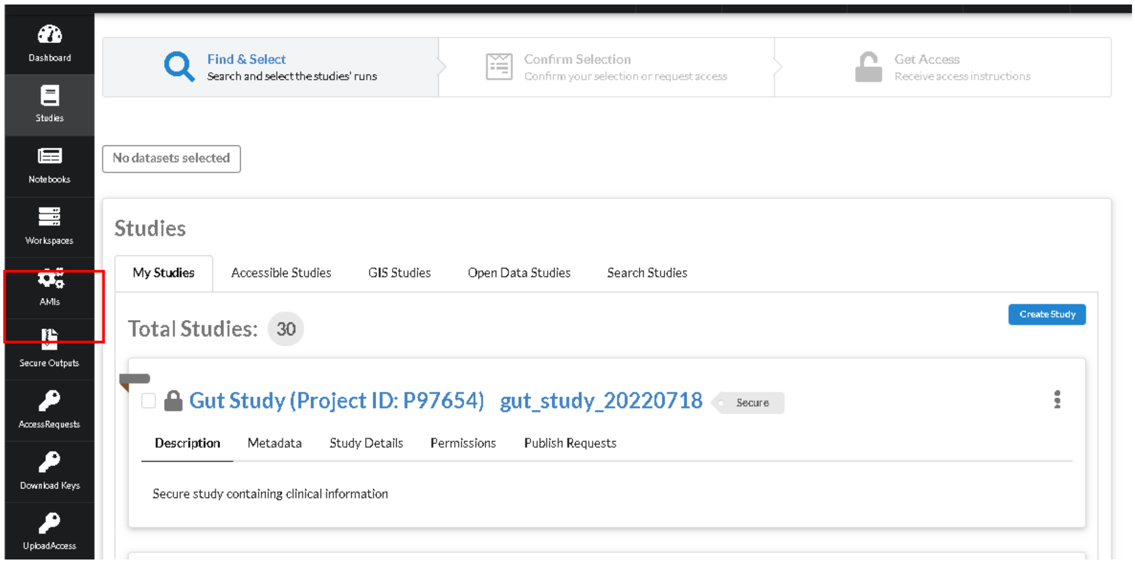


2. Under **Egress Requests**, select the pending secure outputs. Click **Next**.

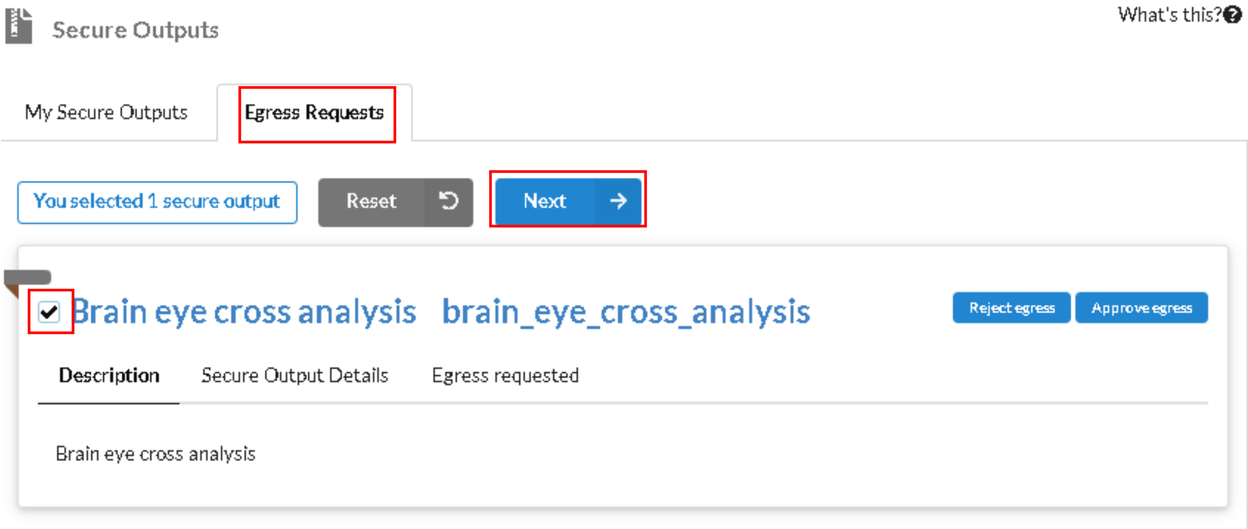

1. Fill up the workspace configuration form:
    a. Name: Workspace name, used to identify your workspaces
    b. Whitelisted CIDR: An IP CIDR value to whitelist access to your workspace. For security, it is highly recommended to set this value to your known IP range/address
    c. Project ID: Project to charge to (only 1 value available at the moment)
    d. Workspace size: Only 1 option available for **Small**
    e. Description: Workspace description

    Once ready, click **Create Vetting Workspace**.

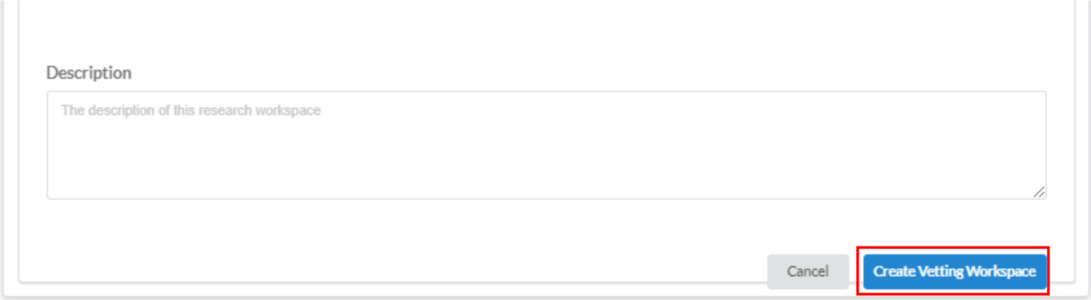


3. Your vetting workspace details will be available in the **Vetting Workspace** subtab of the **Workspaces** tab.

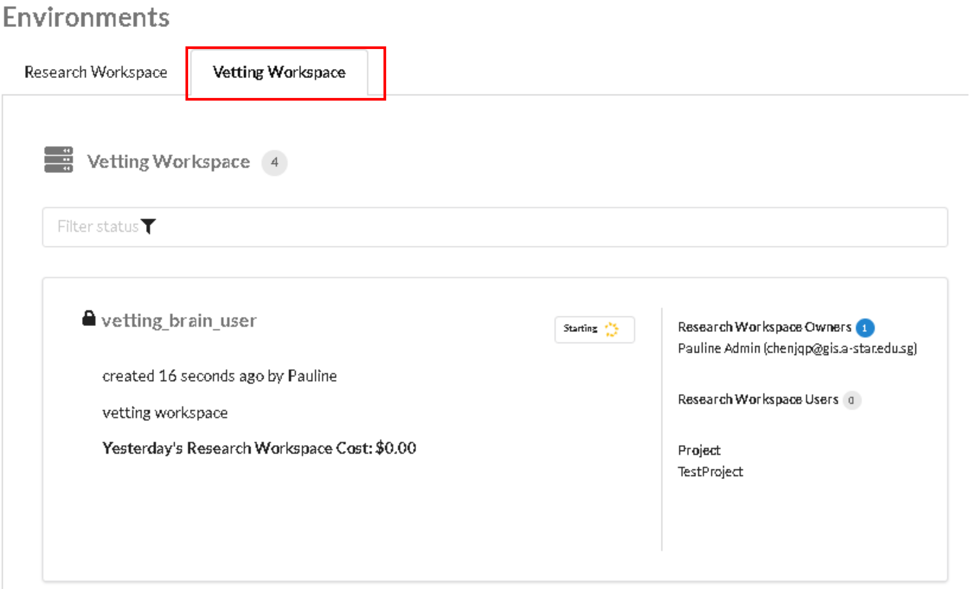


4. To access your vetting workspace, please refer to **Error! Reference source not found.**.

#### Approve/Reject Egress Requests

1. Go to the **Secure Outputs** tab.

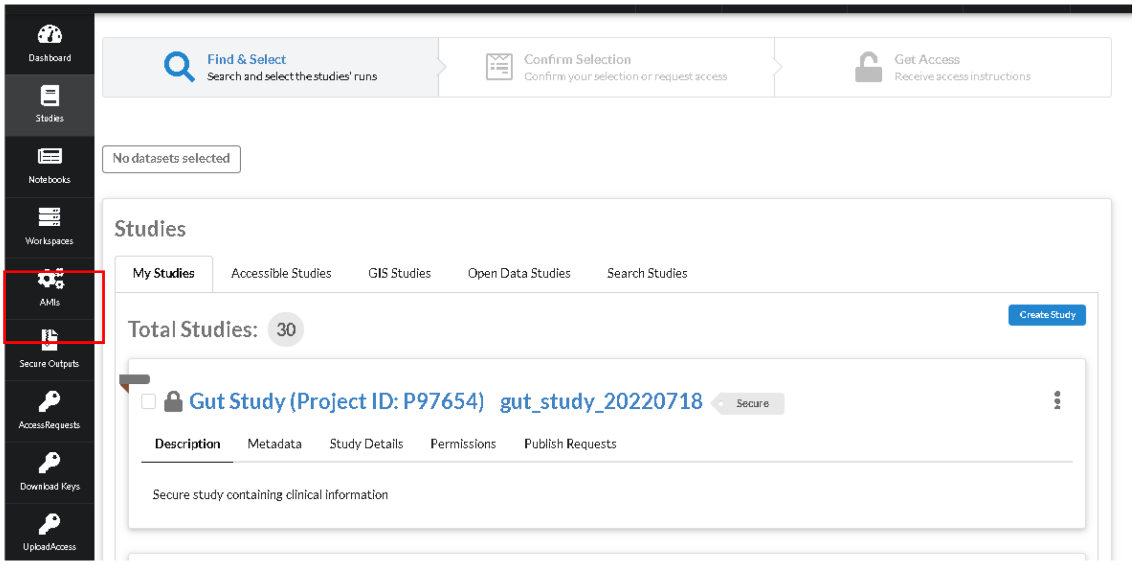


2. Under **Egress Requests**, locate the secure output. Click on **Reject egress** or **Approve egress**.

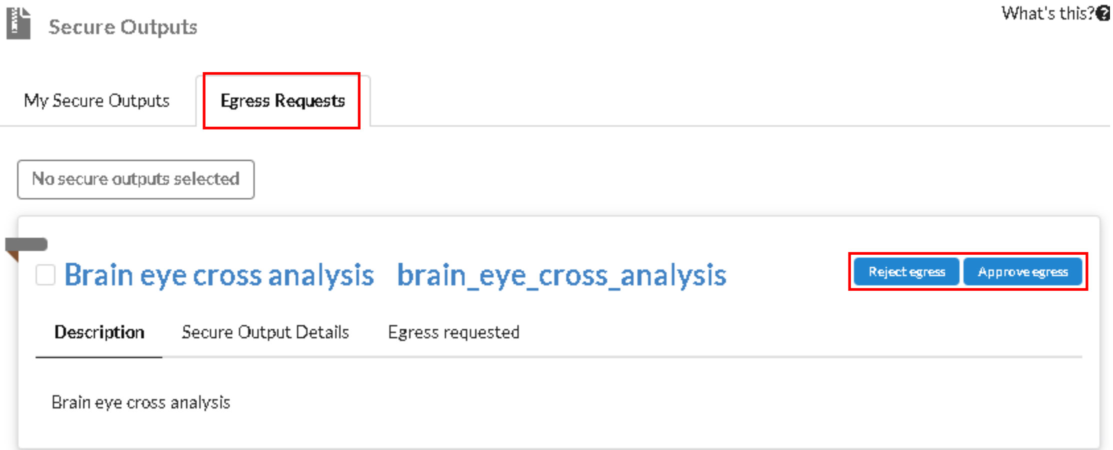


   a. On reject, you will be prompted to provide your comments for rejection. Once done, click **Reject egress**.

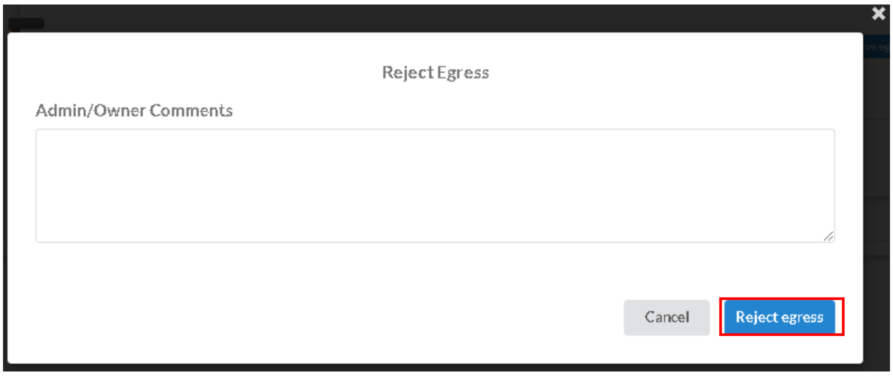


   b. On approval, you will be asked for a confirmation. Click **Approve egress** once you're ready.

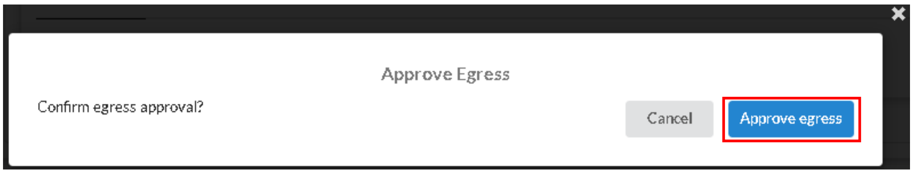
Approve Egress
Confirm egress approval?
Cancel
Approve egress

#### Copy Secure Output to Study

Please note that the secure output must have received approval from all secure study owners to be able to perform this action.

1. Go to the **Secure Outputs** tab.

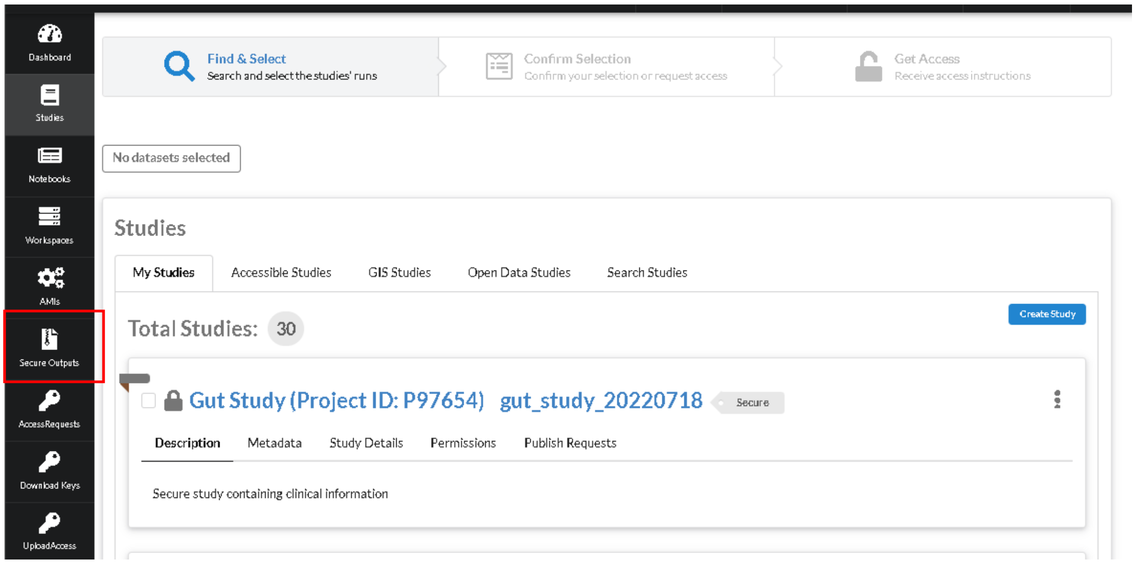


2. Under **My Secure Outputs**, click on **Copy to Study** for the secure output.

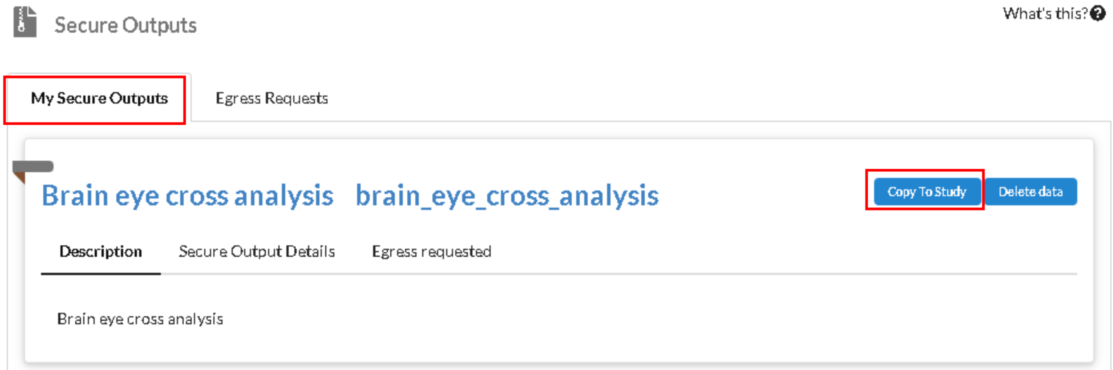


3. Specify the study and prefix to copy to and click **Submit egress**.

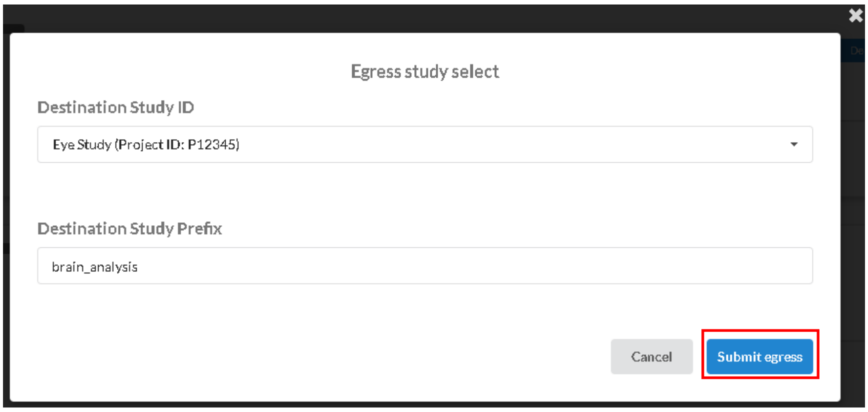

4. To keep track of the status of the egress, click on the **Egress requested** tab of the secure output card.

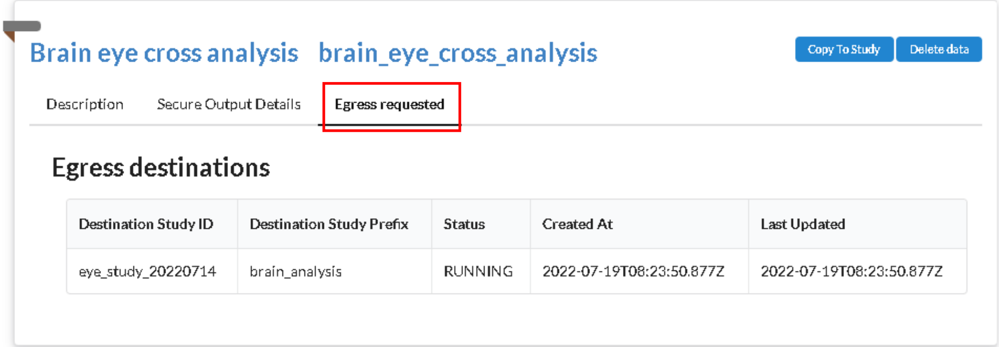


5. The data is fully copied once the status of the egress has changed to **SUCCEEDED**.

#### Delete Data

Please note that the secure output cannot be attached to any workspace or pending approval to perform this action.

1. Go to the **Secure Outputs** tab.

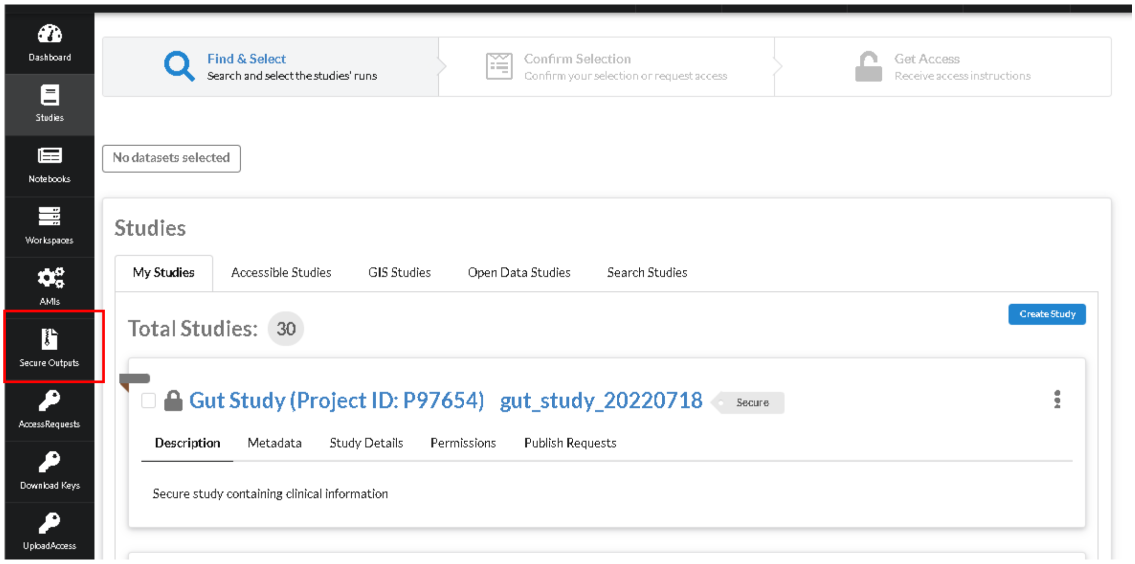


2. Under **My Secure Outputs,** click on **Delete data** for the secure output.

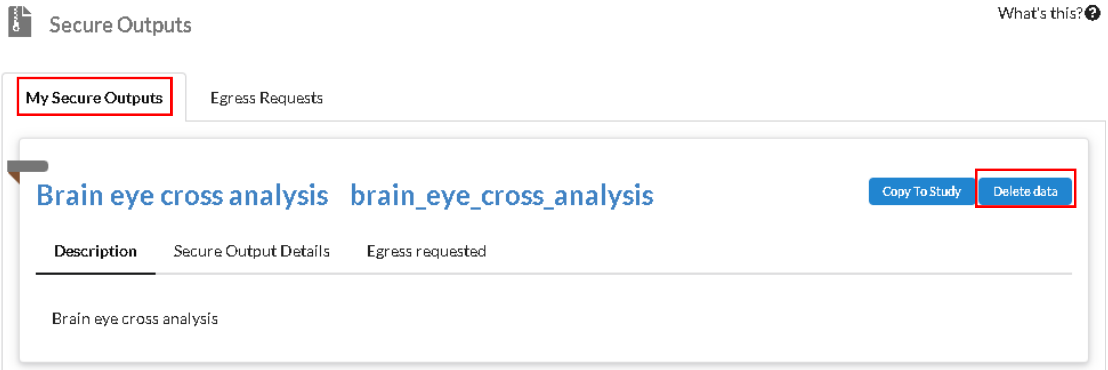

#### Notebooks

A notebook is created when an EMR workspace is spun up by default. The naming convention of the notebook is <workspace-name>-Notebook. Please note that local notebook data needs to be EXPLICITLY SYNCED to the notebook folder to be saved.

##### Request Notebook Access

1. Go to the **Notebooks** tab.

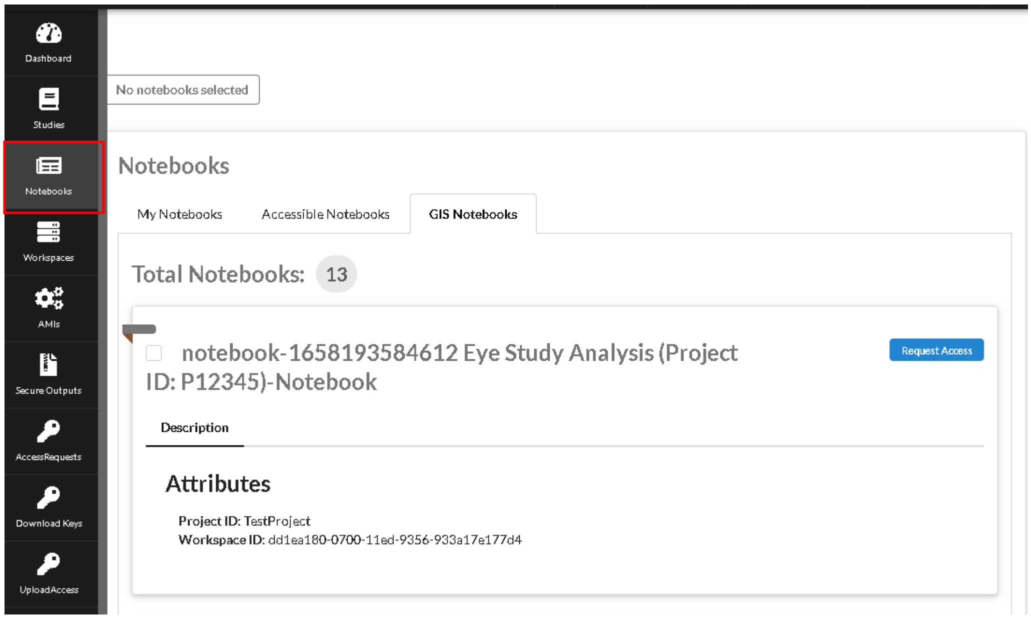


2. Under **GIS Notebooks,** locate the notebook and click on **Request Access**.

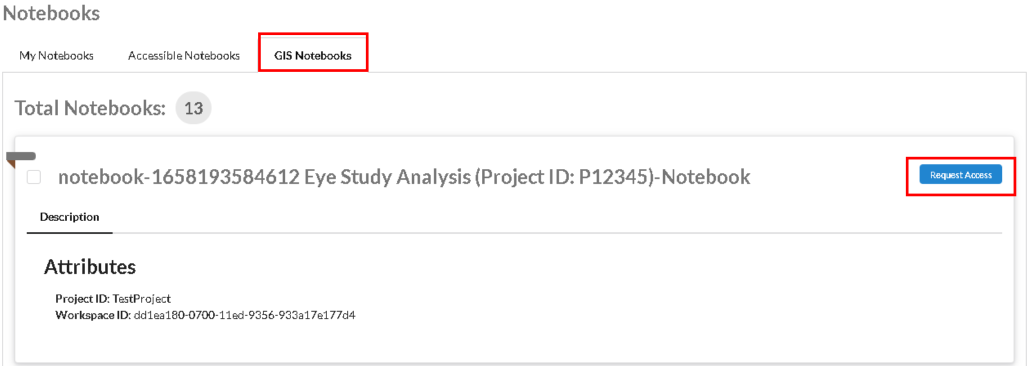


3. Fill up the form and click **Request Access**.

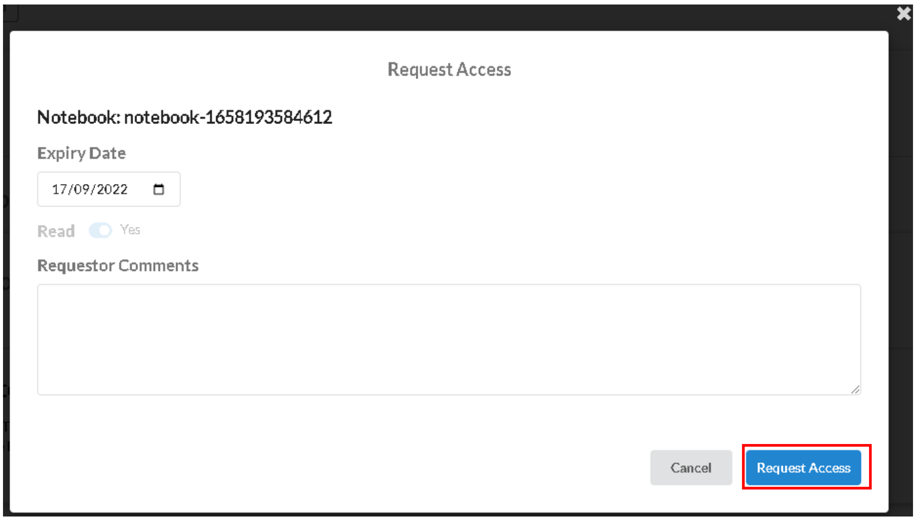

#### Approve/Reject Notebook Access Request

Please note that you will only see access requests of the notebooks you own.

1. Go to the **Access Requests** tab.

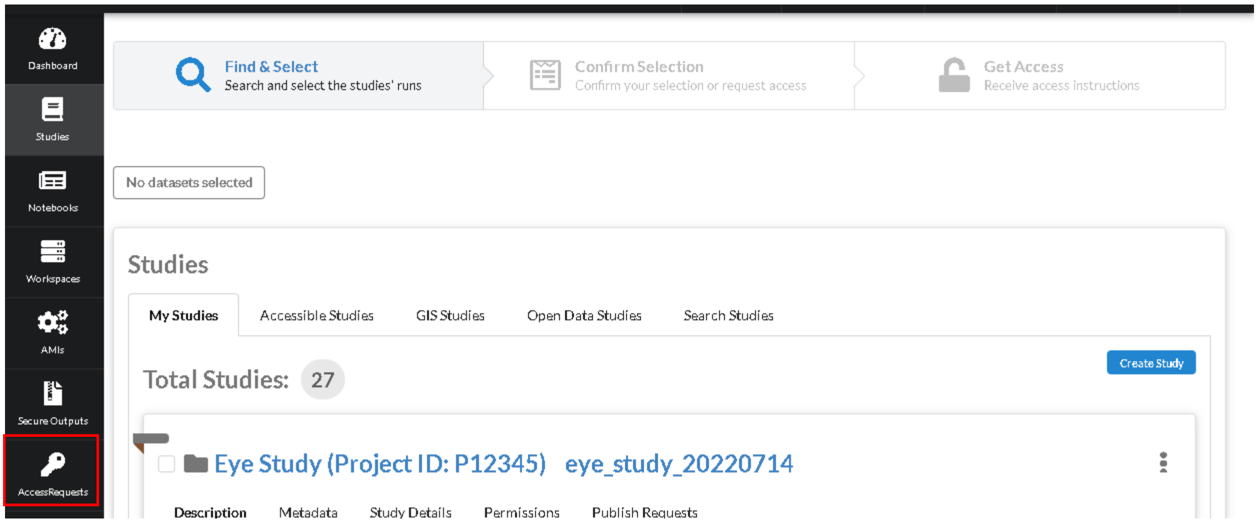


2. Navigate to the notebook. Click on the corresponding buttons to approve/deny the request.

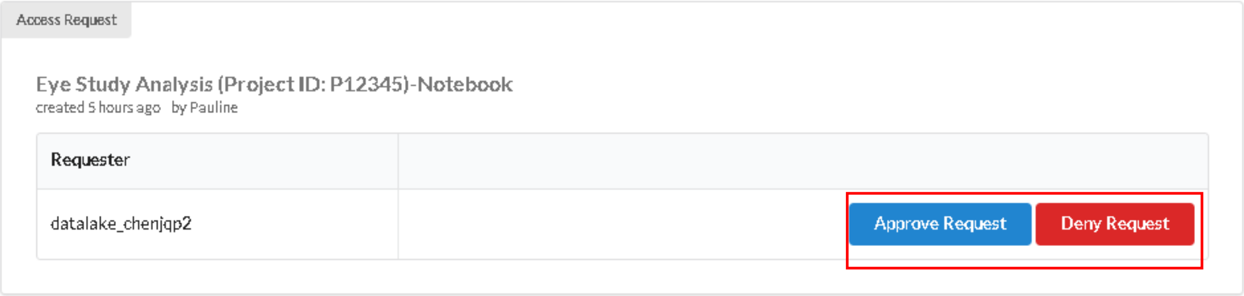

3. On clicking **Deny Request**, you will be presented with a field to provide your comments on rejection (compulsory). Once ready, click **Deny Request**.

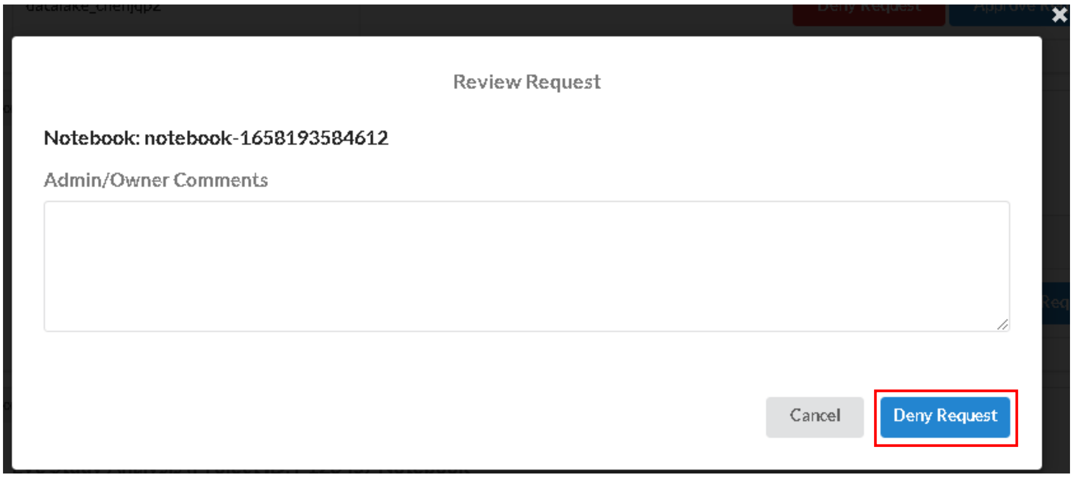


4. On clicking **Approve Request**, you will be presented with the request details and a field to provide your comments on approval (compulsory). Once ready, click **Approve Request**.

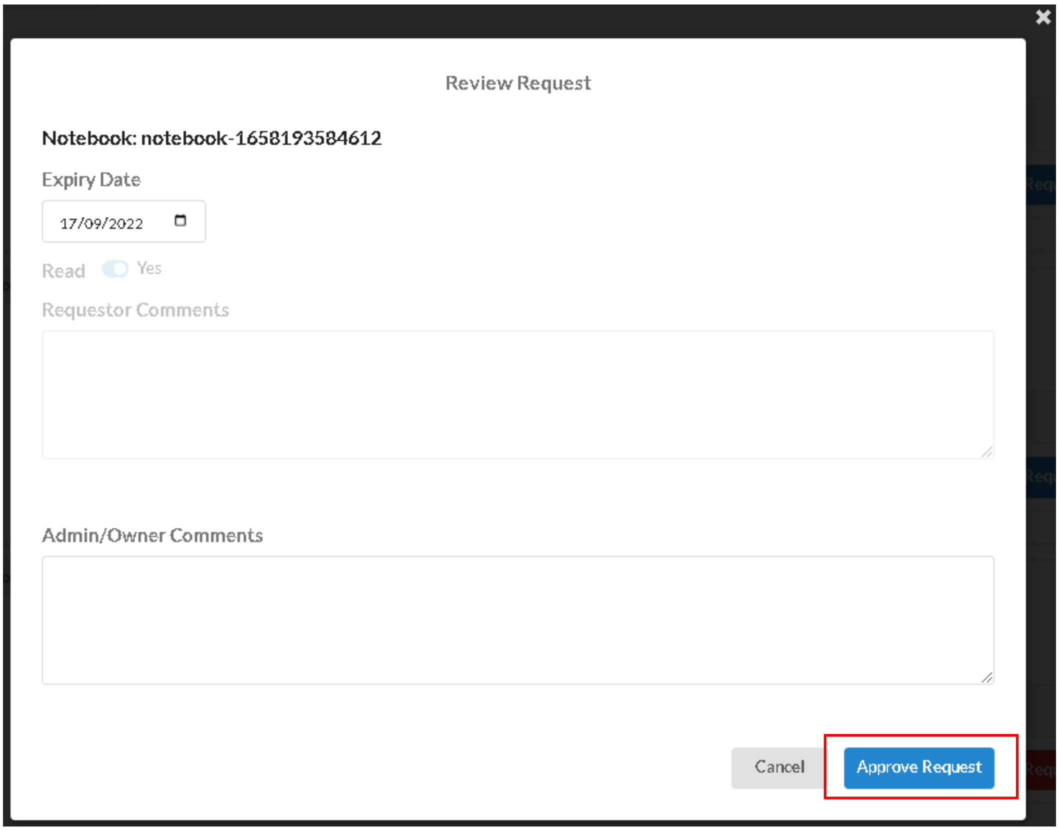

#### Edit Notebook Permissions

Please note that you must be an admin of the study to perform this action.

##### Admin permissions

1. Go to the **Notebooks** tab

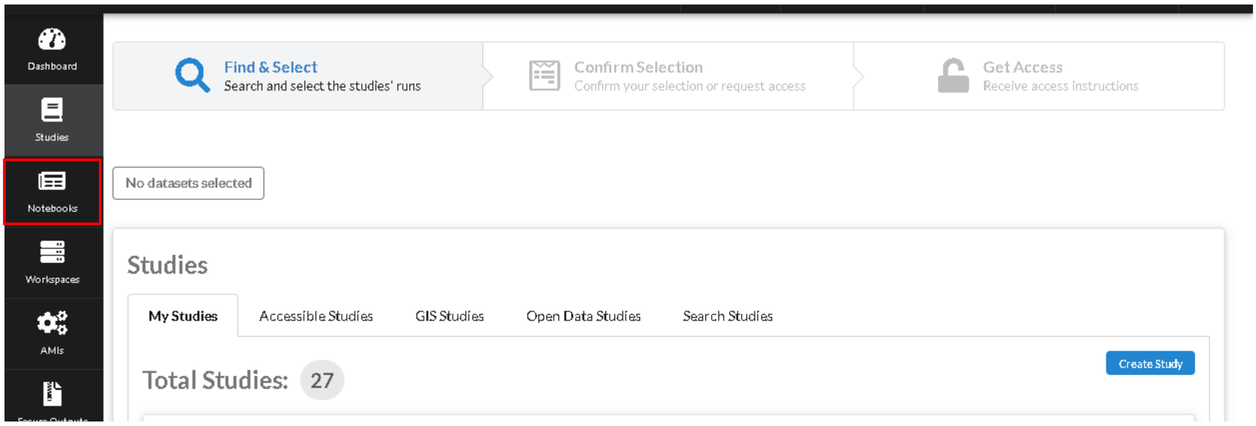


2. Under **My Notebooks** tab, locate the notebook card. Click on **Permissions** sub-tab, then click on the pencil icon beside **Users**.

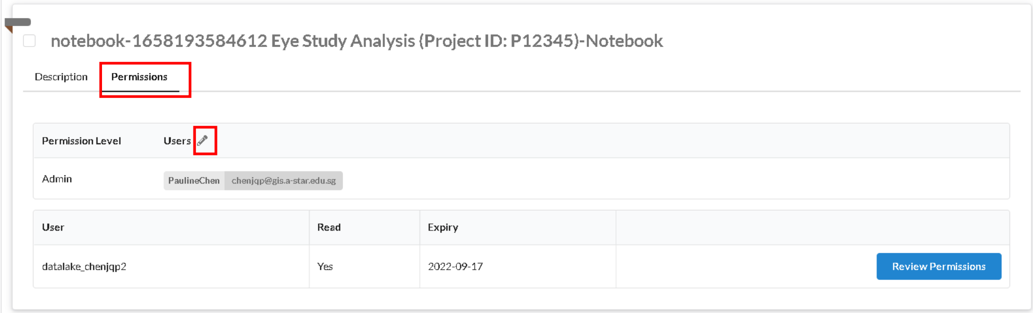


3. Add or delete users into the **Admin** permission level list, then click **Submit**.

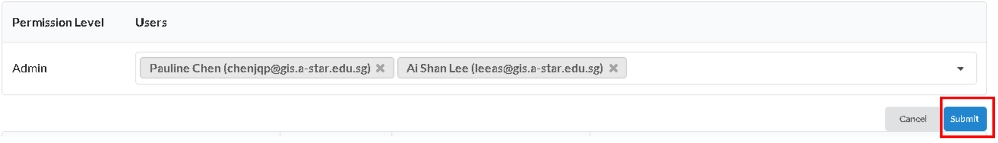

#### User permissions

1. Go to the **Notebooks** tab

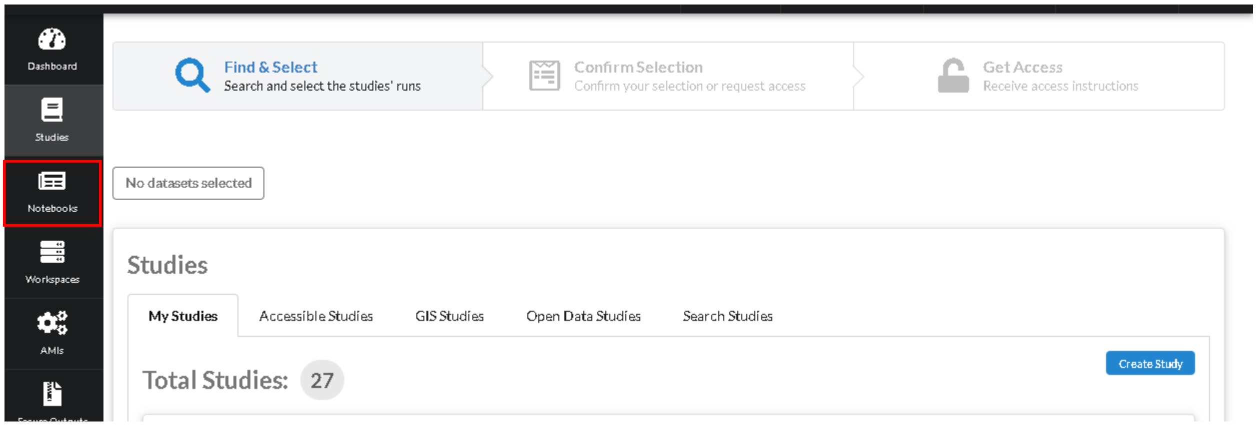


2. Under **My Notebooks** tab, locate the notebook card. Click on **Permissions** sub-tab, then click on **Review Permissions** button of the user whose permission you want to edit.

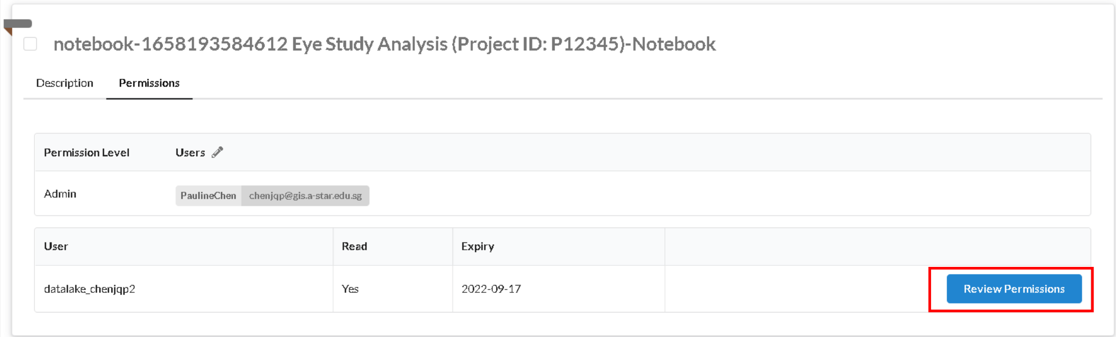


3. Edit the user's permission and click on **Update Permissions**. If you wish to revoke the user's permission completely, click on **Revoke**.

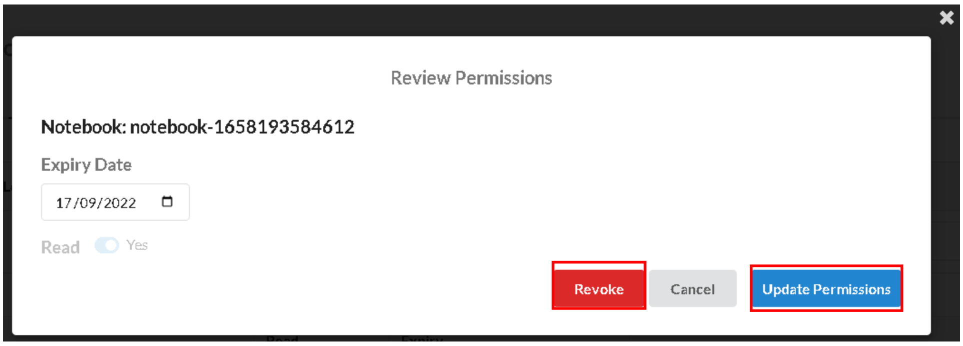

### Create Workspace from Notebook

1. Go to the **Notebooks** tab.

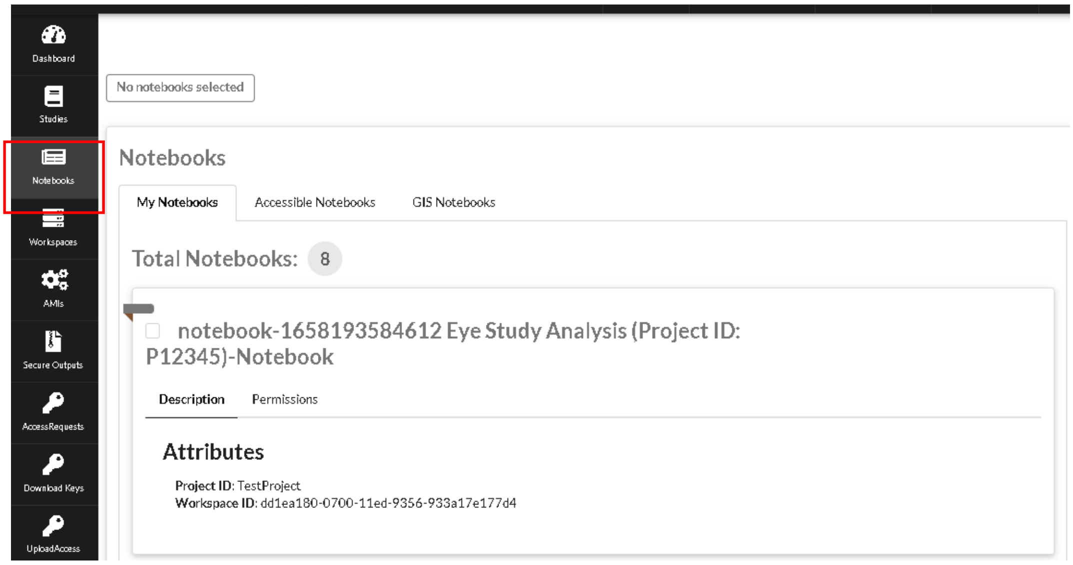


2. Select the notebook(s) and click **Next**.

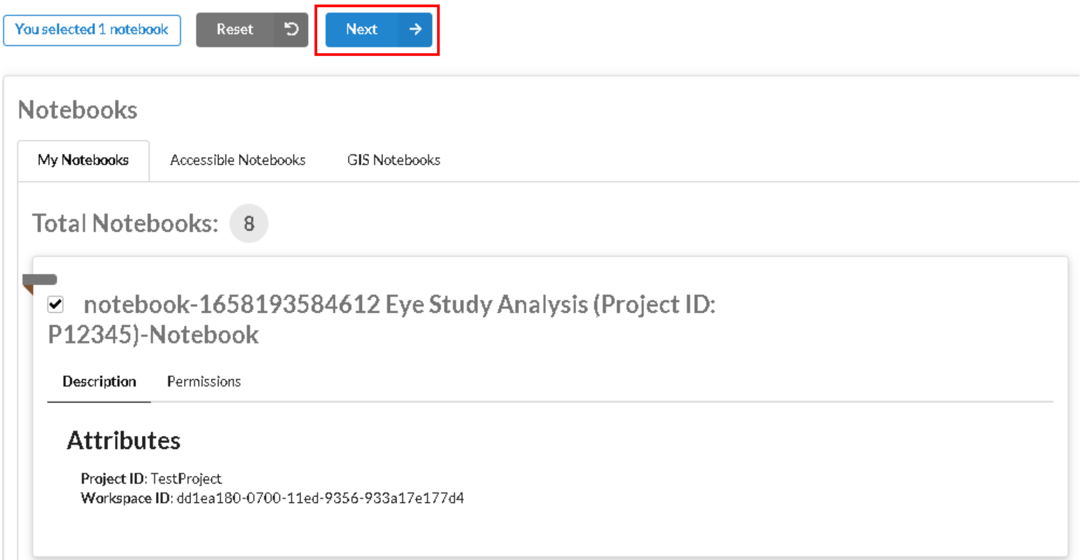

3. Continue from step 2 of Create Workspace. You will only be able to create EMR workspaces.

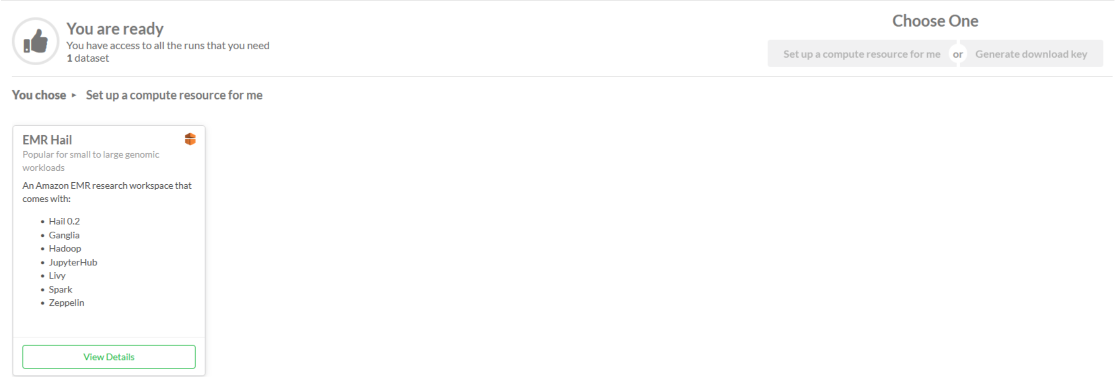
